# Lineage-specific lncRNAs critically determine cross-species differences in tumors

**DOI:** 10.64898/2026.05.01.722350

**Authors:** Jie Lin, Xiuying Liu, Hao Zhu

## Abstract

Diverse mouse models have been generated to study human tumors. Although mouse and human tumors share similar differentially expressed genes and cancer hallmarks, many drugs that work in mice fail in humans. What makes mouse models poorly recapitulate human tumors remains unclear. We postulate that transcriptional regulation by lineage-specific long noncoding RNAs (LS lncRNAs) critically determines cross-species and cross-tumor differences. To test this hypothesis, we identified LS lncRNAs, predicted their target genes, integrated 9,058 RNA-seq samples from 13 human tumors and their mouse counterparts, and analyzed transcriptional regulation by LS lncRNAs across cellular contexts. LS lncRNAs substantially and tumor-specifically reconfigure transcription and signaling, and strongly influence cancer immunity and anti-cancer drug efficacy. These results provide systematic information for exploring and interpreting human tumor mouse models and for identifying human- and tumor-specific diagnostic and therapeutic targets. They also present an analytical approach applicable to other human diseases and their mouse models.

## 1. Introduction

Despite the continuous generation of diverse mouse models to mimic and study human cancer, the gap between animal efficacy and human clinical success remains a persistent challenge (Arrowsmith and Miller, 2013; Day et al., 2015). Investigations of cross-species differences have yielded several key biological discrepancies. First, while fundamental “driver” mutations (e.g., *TP53, KRAS, MYC, APC*) are evolutionarily conserved, human cells require more genetic “hits” than murine cells to undergo malignant transformation (Rangarajan and Weinberg, 2003; Sansom et al., 2006). Second, while core proliferative pathways remain highly concordant between species, pathways governing immune interactions, inflammation, and the cellular microenvironment display strikingly weak conservation (Cheng et al., 2014; Mestas and Hughes, 2004; Seok et al., 2013). Third, many mouse models—ranging from xenografts to genetically engineered mouse models (GEMMs)—frequently fail to recapitulate the co-evolution of the tumor and the host immune system occurring on a human timescale (Fridman et al., 2017). Consequently, deciphering the precise mechanisms that make mouse models poorly reflect human tumors is vital for preventing therapeutic failures and for developing humanized mouse models.

Historically, pan-cancer analyses have illuminated commonalities and differences across tumors (Cancer Genome Atlas Research et al., 2013; Hoadley et al., 2018). However, most analyses examine tumors in humans or mice, prioritize protein-coding genes (PCGs), and emphasize evolutionary similarities but fail to capture the high-resolution, species-specific regulatory modules and networks governed by noncoding RNAs (Haemmerle and Gutschner, 2015; Sarropoulos et al., 2019; Schmitt and Chang, 2016).

Many systems biology studies also explore “network entropy” or “correlation network” primarily based on conserved proteins (Banerji et al., 2013; Becker et al., 2023; Teschendorff and Enver, 2017). Uncovering the causes of cross-species discrepancies and developing humanized mouse models for improving cancer therapy remain challenging (Hegde and Chen, 2020; Olson et al., 2018).

Many human and mouse long noncoding RNAs (lncRNAs) are lineage-specific (Derrien et al., 2012; Huarte, 2015; Yue et al., 2014), and many cell types express precise lncRNA signatures to control lineage-specific regulatory programs (Mattick et al., 2023). Because lncRNAs exhibit exquisite tissue-specificity and rapid evolutionary turnover, they are perfectly positioned to orchestrate the regulatory differences between orthologous cell types across species (Breschi et al., 2017; Hodge et al., 2019). In tumors, the rate of epigenetic changes can be orders of magnitude higher than that of genetic changes (Easwaran et al., 2014; Flavahan et al., 2017), and lncRNA-mediated gene regulation contributes critically to cancer metastasis and immunity (Guo et al., 2021; Liu et al., 2021). However, how lineage-specific (LS) lncRNAs influence cross-species discrepancies remains underexplored. Moreover, it remains poorly distinguished whether regulatory modules and networks are structurally rewired, merely hijacked, or completely invented *de novo* by these noncoding transcripts.

To address these distinct gaps, we developed an LS lncRNA-centered comparative pan-cancer framework. By identifying LS lncRNAs, predicting lncRNA-DNA binding, and integrating 9,058 RNA-seq samples from 13 human and mouse tumors, we systematically analyzed transcriptional regulation across distinct cellular contexts. Our results demonstrate that LS lncRNAs critically determine cross-species and cross-tumor divergence in transcriptional regulation and signaling. Ultimately, this framework sheds new light on the regulatory foundations of cross-species variations, providing a vital asset for identifying human- and tumor-specific therapeutic targets and translating pre-clinical mouse research into viable human therapies.

## 2. Results

### 2.1 Transcriptional divergent genes reveal cross-species differences in tumors

To systematically investigate how LS lncRNAs drive species-specific tumor biology, we first needed to define the baseline transcriptional divergence of their downstream protein-coding targets across species. We downloaded raw RNA-seq counts for human tumors and their mouse models from the UCSC Xena and EMBL-EBI websites (Supplementary Table 1) (Goldman et al., 2020; Petryszak et al., 2017). After quality control (Supplementary Fig. 1-1), we retained 13 matched tumor types with at least 20 cancer and 20 normal samples in both species (except for normal human bladder samples, n = 9). The evaluated tumor types included bladder, breast, kidney, liver, lung, pancreatic, prostate, ovarian, skin (melanoma), glioblastoma, glioma, leukemia, and lymphoma (comprising 5,077 human cancer, 1,867 human normal, 1,052 mouse cancer, and 1,062 mouse normal samples).

To identify differentially expressed genes (DEGs), raw counts were rescaled to z-scores across all samples (termed normalized expression, NX). Using the difference between the median NX values in cancer and normal samples (ΔNX), we identified DEGs within each species (|ΔNX| > 1 and false discovery rate (FDR) < 0.05). We then measured expression differences across species, specifically using one-to-one orthologous protein-coding genes (1:1 PCGs). Similar to recent comparative studies (Hodge et al., 2019), we computed cross-species differential expression (ΔNX = NX_human − NX_mouse) based on the median cancer sample values. Based on these metrics, we classified 1:1 PCGs into transcriptionally divergent genes (TDGs; |ΔNX| > 1, FDR < 0.05), transcriptionally intermediate genes (TIDs; 0.5 < |ΔNX| < 1.0, FDR < 0.05), and transcriptionally conserved genes (TCGs; |ΔNX| < 0.5, FDR < 0.05). This >1 cutoff for TDGs parallels the conventional DEG criteria used within a single species, enabling direct comparison between intra-species and cross-species variation. Across the 13 tumors, the NX-based differential expression of all 1:1 PCGs exhibited significant cross-species Pearson correlations (Supplementary Figs. 1-2, 3). Furthermore, analysis of similarities (ANOSIM) with 10,000 permutation tests revealed that intra-species cross-tumor variations were significantly smaller than inter-species variations (p < 10^−4^) (Supplementary Fig. 1-4) (Somerfield et al., 2021). This establishes that our identified TDGs more effectively capture species-specific tumor features than conventional intra-specific DEGs (Fig. 1A–D; Supplementary Figs. 1-5, 6; Supplementary Tables 2, 3).

**Figure 1.**
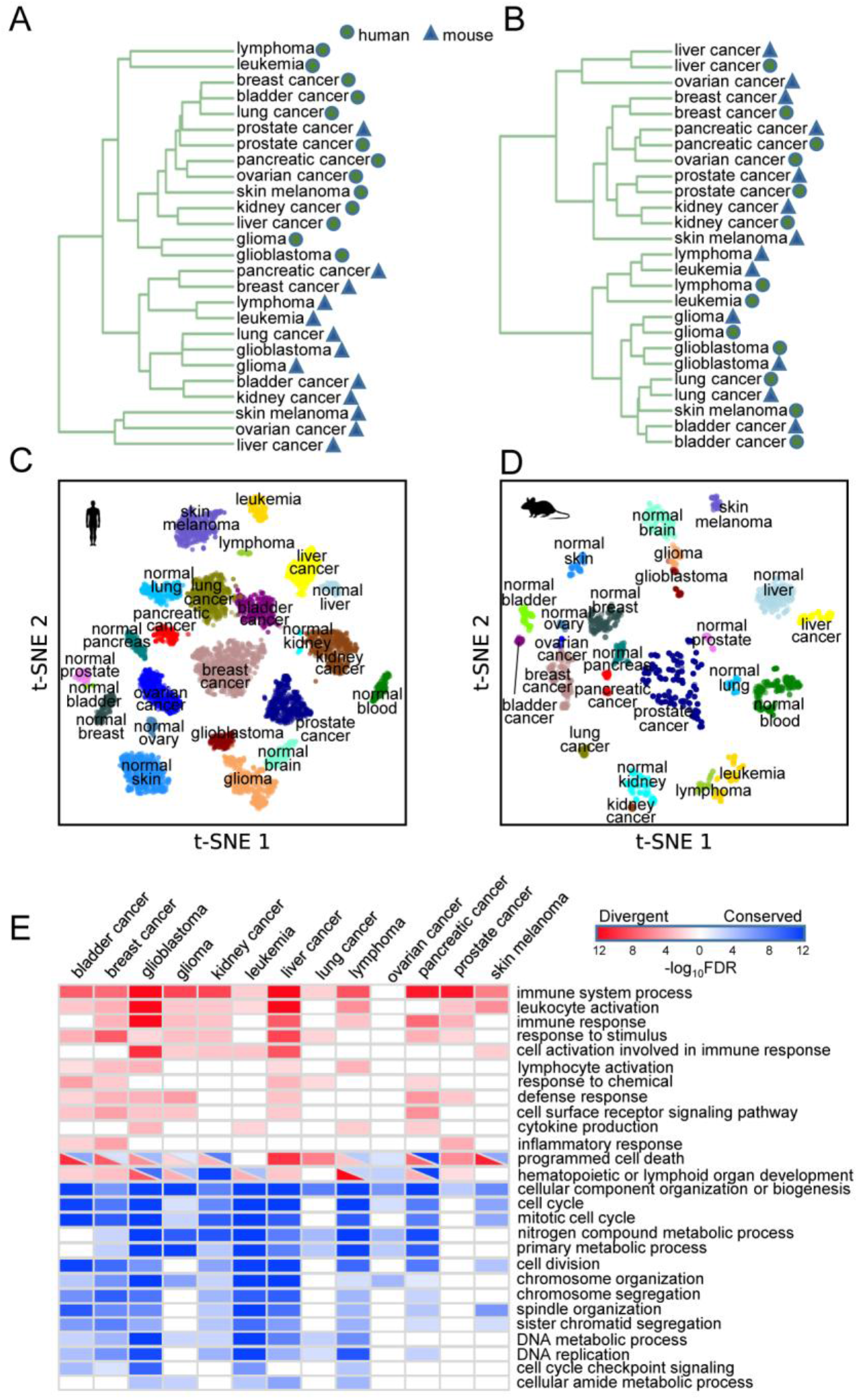
TDGs capture species- and tumor-specific expression features. (A) Hierarchical clustering of samples from 13 tumors based on the top 100 TDGs. (B) Hierarchical clustering of the same samples based on the top 100 TCGs. (C) t-SNE clustering of 6,944 human samples based on NX values. (D) t-SNE clustering of 2,114 mouse samples based on NX values. (E) TDGs (top) and TCGs (bottom) are enriched for distinct Gene Ontology (GO) terms.

Because bulk RNA-seq data reflects a mixture of cell populations, infiltrating immune cells intrinsically contribute to the detected expression levels. To ensure our cross-species divergence was not merely an artifact of differential immune infiltration between species, we evaluated the ΔNX values of established marker genes for ten infiltrating immune cell types (Zhang et al., 2019). In each tumor, approximately half of these immune markers showed human-biased expression (ΔNX > 0) while the other half showed mouse-biased expression (ΔNX < 0). This balanced distribution strongly indicates that the observed TDG variations are primarily driven by tumor-intrinsic, species-specific transcriptional regulation, rather than broad discrepancies in immune cell infiltration (Fig. 1E; Supplementary Fig. 1-7).

Having established the landscape of species-divergent protein-coding targets, we next evaluated the overarching transcriptional regulation mediated by LS lncRNAs. We identified LS lncRNAs based on orthologs of GENCODE-annotated human (17,958) and mouse (13,450) lncRNAs across mammals. This yielded 1,615 primate/simian-specific (PS/SS) lncRNAs and 1,377 rodent-specific (RS) lncRNAs. Differentially expressed LS lncRNAs were defined as those exhibiting transcripts per million (TPM) > 0.1 in at least 50% of samples and an absolute ΔNX > 1 (FDR < 0.05) in at least one tumor type (Supplementary Table 4). Distinctly separating the 13 tumors by species, t-Distributed stochastic neighbor embedding (t-SNE) mapping based on LS lncRNA expression suggests that these noncoding elements critically define cross-species tumor identity. Furthermore, Jaccard distance measurements confirmed that LS lncRNAs are significantly more cancer-specific than DEGs. To directly analyze their transcriptional mechanisms, we computationally predicted the corresponding DNA binding sites (DBSs) for these LS lncRNAs within the 5,000 bp promoter regions of the identified TDGs (Supplementary Tables 5, 6) (Wen et al., 2022).

### 2.2 LS lncRNAs species- and tumor-specifically regulate transcription

Because traditional transcription factors (TFs) are highly conserved across mammals, TF-based expression networks often fail to fully capture cross-species regulatory divergence. We bypassed this limitation by investigating transcriptional regulation mediated by LS lncRNAs. Specifically, we focused on genes exhibiting both differential expression and divergent transcription and possessing predicted LS lncRNA DBSs in their promoters. We applied the *eGRAM* program to identify distinct regulatory modules comprising LS lncRNAs as regulators and these DEG-TDGs (differentially expressed TDGs) as downstream targets (He et al., 2026).

During tumorigenesis, cancer cells can rewire or hijack existing transcription and signaling programs (e.g., basic metabolism, cell cycle machinery, lineage-survival signals) or create entirely new ones. However, prior comparative studies have poorly differentiated which specific programs change, remain unchanged, or are invented *de novo*, and crucially, whether mouse models utilize the same programs as human tumors or rely on distinct, species-specific modules. To address this, we identified regulatory modules jointly based on lncRNA-DBS targeting and expression correlation across three conditions: Normal (using only normal samples), Cancer (using only cancer samples), and Preserved (where LS lncRNA-target pairs maintain mathematically strong, directionally consistent expression correlations across both normal and cancer samples). Tracing modules across these conditions effectively distinguishes altered, newly invented, and hijacked programs. Because *eGRAM* calculates enriched KEGG and WikiPathways terms for each module, we mapped these modules to established functional axes by assigning a specific cancer hallmark to each module based on its dominant KEGG pathways (Hanahan, 2022, 2026).

To analyze these modules across the 13 tumors and three conditions, we projected them into a unified hallmark landscape using Uniform Manifold Approximation and Projection (UMAP). The resulting maps revealed striking topological properties of LS lncRNA-mediated regulation. First, total module numbers varied substantially across both species and tumors (Fig. 2A), and target gene sets exhibited varied Jaccard distances (Fig. 2B), indicating profound heterogeneity in LS lncRNA-mediated regulatory organization. (Because *eGRAM* merged modules sharing ≥80% of target genes, a tumor lacking lineage-specific regulation would inherently exhibit uniform module numbers across species).

**Figure 2.**
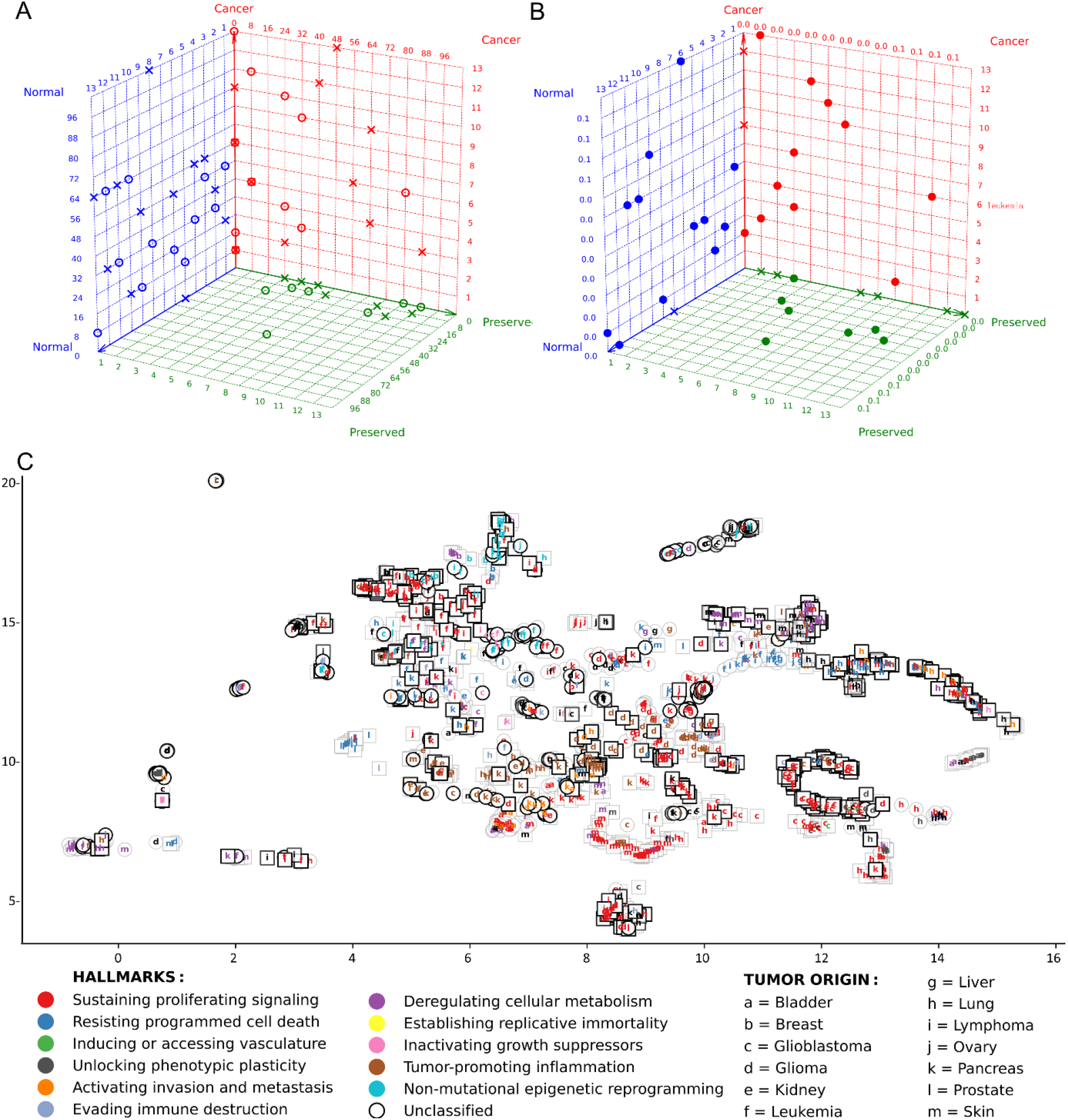
Differences in regulatory modules and cancer hallmarks between human and mouse tumors. (A) The number of identified modules in human and mouse tumors under Normal, Cancer, and Preserved conditions. Circles and crosses indicate humans and mice, respectively. (B) Jaccard distances between target gene sets across species. Dots represent the average human–mouse similarity score for each corresponding tumor; crosses indicate an empty result. (C) UMAP projection of hallmark-assigned modules across 13 tumors in both humans and mice under the Normal and Cancer conditions. Circular and square nodes map to human and mouse modules, respectively. Light-colored edges represent the Normal condition, while dark-colored edges denote Cancer conditions (Note: A remote, densely packed cluster of *Unclassified* modules was excluded for visual clarity).

Second, more regulatory modules were identified in the Normal condition compared to the Cancer condition. This suggests that oncogenic transformation forces cells away from diverse, specialized regulatory programs—which normally maintain tissue identities and respond to microenvironmental cues—and collapses them toward a restricted set of dominant oncogenic axes. These lost programs likely act as homeostatic barriers dismantled during tumorigenesis; their erosion may prevent the recovery of normal cellular fates.

Third, UMAP distances between modules highlighted vast architectural differences between human tumors and their mouse counterparts (Fig. 2C). Fourth, the Preserved landscape is nearly empty for several tumors (e.g., bladder, liver) but densely populated for others (e.g., glioblastoma, leukemia) (Supplementary Fig. 2-1). Because Preserved modules explicitly represent retained but altered programs, an empty Preserved landscape indicates a complete failure of the tumor to hijack existing homeostatic programs. Instead, the native regulatory syntax is completely eradicated during transformation (reflected by the dismantled Normal-condition modules), and the tumor is forced to rely exclusively on invented, *de novo* survival programs (captured by the independent Cancer-condition modules). In contrast, heavily populated Preserved landscapes (e.g., in human glioblastoma and leukemia) illustrate a different biological reality: these tumors survive by heavily hijacking pre-existing neurodevelopmental and germinal-center epigenetic programs, locking these altered regulatory programs into pathological overdrive (Wainwright and Scaffidi, 2017; Wu et al., 2025).

Finally, a few specific tumors (notably pancreatic and leukemic cancers) displayed a distinct cross-species co-localization of modules dominated by *Tumor-promoting inflammation* and *Sustained proliferative signaling*, pointing to a partial evolutionary conservation of their regulatory architecture. Together, these findings indicate that malignant transformation requires a systematic topological reconfiguration of a cell’s regulatory machinery, challenging the prevailing assumption that tumors simply “overdrive” normal pathways or exist primarily as a state of transcriptional “high entropy” or “chaos” (Banerji et al., 2013; Teschendorff and Enver, 2017).

### 2.3 Unified hallmark landscapes reveal cross-species and cross-tumor differences

We next examined the unified hallmark landscapes of the 13 tumors, where the dominant KEGG pathways defining each module’s cancer hallmark determine its distribution in the UMAP space (Supplementary Fig. 3-1∼13). In most solid tumors, human and mouse modules segregated into distinct clusters under both Normal and Cancer conditions, indicating substantial cross-species divergence in LS lncRNA-mediated regulatory organization. These divergences likely reflect the cumulative effects of species-specific stromal interactions, varying immune microenvironments, and distinct mutational trajectories. In sharp contrast, leukemia modules from humans and mice intermingled across all three conditions (Fig. 3A). This cross-species integration is consistent with established biology demonstrating that hematopoietic differentiation programs are evolutionarily conserved between humans and mice (Bradner et al., 2017; Krivtsov et al., 2006; Somervaille et al., 2009), and that leukemogenesis is driven primarily by cell-intrinsic epigenetic reprogramming and clonal evolution (Ferrando and Lopez-Otin, 2017; Shih et al., 2015). This functional convergence suggests that murine leukemia models far more closely recapitulate the core regulatory architecture of the human disease than many solid tumor models.

**Figure 3.**
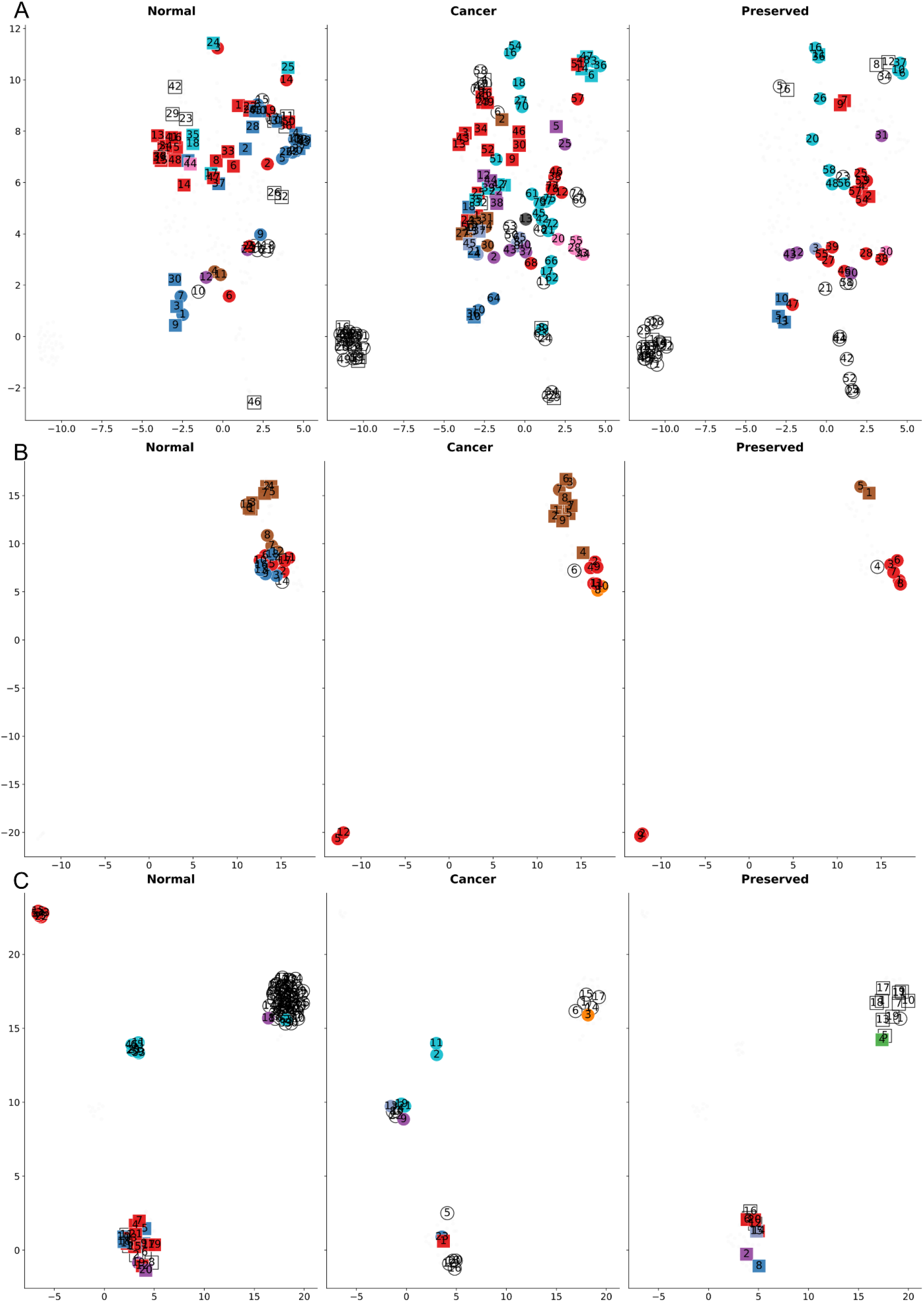
UMAP projection of leukemia, kidney cancer, and ovarian cancer modules based on hallmarks. Circular and square nodes map to human and mouse modules; colors designate hallmarks (see Fig. 2C); and numbers indicate modules. (A) Modules of leukemia. Human and mouse modules intermingle seamlessly, with a high density of modules maintained across both Cancer and Preserved conditions. (B) Modules of kidney cancer. The projection reveals multifaceted regulatory architecture in human tumors versus simple, inflammation-restricted architecture in mouse tumors. (C) Modules of ovarian cancer. In the Preserved condition, the *Unclassified* modules exhibit stark species-specific segregation, corresponding to the enrichment of entirely distinct WikiPathways terms between humans and mice.

In human and mouse kidney cancer, our analysis identified *Tumor-promoting inflammation* as a dominant hallmark among modules in the Preserved condition (Fig. 3B), indicating that this inflammatory program is heavily maintained during tumorigenesis in both species. This aligns with clinical observations that advanced clear cell renal cell carcinoma (ccRCC) is highly immunogenic and characterized by profound inflammatory infiltration (Jonasch et al., 2014; Thorsson et al., 2019). However, mapping this landscape simultaneously revealed a stark cross-species divergence: while murine modules remained restricted exclusively to the inflammatory hallmark across all three conditions, human modules systematically incorporated *Sustaining proliferative signaling* and *Resisting programmed cell death*. Pathway enrichment analysis corroborated this divergence. In the Preserved condition, human modules were uniquely enriched for specific pathways, including *Regucalcin in proximal tubule epithelial kidney cells* (WP4838) and *Focal adhesion* (hsa04510)—programs intimately tied to renal tubular cell survival and regeneration. These human modules likely capture the continuous VHL/HIF-driven proximal tubule regenerative program, which is constitutively hijacked during human ccRCC pathogenesis (Cancer Genome Atlas Research et al., 2013). This stark contrast between simple murine and multifaceted human regulatory architectures directly reflects a known translational limitation: standard murine models of kidney cancer (e.g., RENCA) are predominantly driven by rapid inflammatory responses, whereas human patients undergo slow, multi-hit metabolic shifts (Frew and Moch, 2015; Hsieh et al., 2017). Consequently, while murine models may mimic the gross inflammatory phenotype of RCC, they critically lack the LS lncRNA regulatory syntax required to faithfully model the complex proliferative and survival programs intrinsic to human kidney cancer progression.

Ovarian cancer provides a third compelling example of this divergence. The highly separated human and murine clusters across all three conditions mapped a clear topological dichotomy, implying fundamentally distinct functional programs. In the Preserved condition, we identified multiple murine *Unclassified* modules, but only one human *Unclassified* module. Notably, these modules were enriched for mutually exclusive, species-specific WikiPathways terms. The human module was uniquely enriched for *Homologous recombination* (WP186), *DNA repair pathways* (WP4946), and *Pathways affected in adenoid cystic carcinoma* (WP3651). Conversely, the murine modules were exclusively enriched for *EGFR1 signaling pathway* (WP572), *Estrogen signaling* (WP1244), and *IL-6 signaling pathway* (WP387). This topological divergence captures a widely recognized translational roadblock in ovarian cancer research. Preclinical murine models—including syngeneic ID8, transgenic, and xenograft models— exhibit a strict dependency on classical growth factor and hormone axes; blocking EGFR or estrogen signaling in these models effectively halts tumor progression (Laws et al., 2014; Yin et al., 2016). However, this therapeutic vulnerability fails to translate to human disease. While *EGFR* is overexpressed in up to 70% of human ovarian cancers and correlates with poor prognosis, recent cohorts confirm that true oncogenic *EGFR* driver mutations occur in only ∼0.08% of patients, readily explaining the failure of EGFR-targeted therapies in clinical trials (Gower et al., 2025; Vergote et al., 2014). Instead, human ovarian cancer is largely driven by genomic instability. Characterized by profound DNA repair dysfunctions (Miras et al., 2025), homologous recombination deficiency (HRD) occurs in approximately 50% of high-grade serous ovarian cancers (Khokhlova et al., 2025; Quesada et al., 2025). Thus, the functional pathways preserved within the human module accurately reflect the true, genome-instability-driven vulnerabilities of human ovarian cancer. Furthermore, the enrichment of WP3651 points toward ovarian adenoid cystic carcinoma (OACC), a specific “adenoid cystic-like” subtype of ovarian cancer (Eichhorn and Scully, 1995; Ho et al., 2013). The emergence of this highly specific signature confirms that the identified regulatory modules and their corresponding UMAP manifolds are sufficiently sensitive to capture both core and non-canonical regulatory syntaxes across the broader landscape of ovarian cancer.

### 2.4 Identifying therapeutic targets based on functional and module landscapes

These hallmark landscapes provide specific, testable predictions for therapeutic targets. For example, in human ovarian cancer, the Preserved landscape is reduced to a single *Unclassified* module (ME1) in which the regulator set comprises *AL513314.2* and *AC060780.1* and the target set contains key DNA repair genes *ATM, ERCC3, RAD52*, and *REV1*. While both *RAD51B* and *RAD52* were downregulated DEGs in the evaluated human ovarian cancer cohort—and canonical homologous recombination is frequently associated with *RAD51*—our algorithm specifically identified *RAD52* as the exclusive target regulated by this lncRNA axis. This regulatory specificity may be biologically profound: RAD52 operates as a critical, independent backup mechanism for homologous recombination in BRCA-deficient cells (Konstantinopoulos et al., 2015; Rossi et al., 2021). Because human ovarian cancers (especially high-grade serous ovarian cancers) are characterized by loss of p53 function and extensive chromosomal instability (Cancer Genome Atlas Research, 2011), we reason that the preserved, RAD52-dependent module regulated by *AL513314.2* and *AC060780.1* may be selectively maintained to buffer replication stress. We hypothesize that this regulatory axis may represent a context-dependent vulnerability, and perturbing it may induce synthetic lethality or re-sensitize resistant cells to PARP inhibitors—a therapeutic strategy recently supported by preclinical models demonstrating that RAD52 blockade successfully overcomes PARP inhibitor resistance in BRCA2-deficient ovarian cancer (Ota et al., 2025).

In sharp contrast, murine ovarian cancer models retain vastly more preserved modules, which are heavily enriched for the *Fanconi anemia pathway* (mmu03460) and *Nucleotide excision repair* (mmu03420). This comparative landscape reveals that, while both species must urgently manage genomic instability to survive, they exploit completely different LS lncRNA syntaxes to achieve it. For preclinical validation, the murine orthologous functional axis (*4930481A15Rik* →*Palb2, Brca2, Fancd2* →Fanconi Anemia) suggests that *4930481A15Rik* expression should be monitored as a biomarker of regulatory response in mice.

Our analysis also identified murine-specific artifacts. Specifically, the murine axis *AU020206* →*Cyp11a1, Star* →Steroidogenesis (mmu04925, mmu04927) may anchor a prominent regulatory program governing local hormone synthesis. This axis likely represents an artifact of the murine ovarian microenvironment, rendering the mouse tumor intrinsically dependent on local steroid hormones. This divergence elegantly aligns with the clinical observation that therapies targeting hormone signaling or growth factors often exhibit impressive preclinical efficacy in murine ovarian cancer models, yet fail to improve survival in human trials (Shao et al., 2026). Our data suggest that the human regulatory landscape has largely shed this steroidogenic dependency, offering a mechanistic explanation for translational failure that warrants experimental testing.

Because mapping hallmark landscapes relies on predefined hallmarks, it inherently introduces pathway biases—frequently resulting in *Unclassified* modules, particularly when these modules drive highly novel or tissue-specific functions. To bypass these biases and evaluate regulatory architecture at a higher, unbiased resolution, we projected the modules across all three conditions into a unified UMAP manifold based strictly on the Jaccard distances of their raw target gene sets (Supplementary Fig. 4-1∼13). These unguided landscapes reveal that human and murine modules generally maintain highly species-specific clustering, but there are notable exceptions. In kidney cancer (Fig. 4A; Supplementary Figure 4-5), three human modules (Normal ME15, Cancer ME7, and Preserved ME5) are clustered together with the murine cluster. In the hallmark landscape (Fig. 3B), human Cancer ME7 co-localizes with mouse Cancer modules and human Preserved ME5 co-localizes with mouse Preserved ME1, all with the *Tumor-promoting inflammation* hallmark. This dual-landscape concordance of these modules suggests that, while mouse models fail to recapitulate human tumor proliferation and survival mechanisms, they may utilize the same orthologous downstream genes to mirror the human inflammatory microenvironment. In ovarian cancer (Fig. 4B; Supplementary Figure 4-10), three human modules (Normal ME26, Normal ME61, and Cancer ME3) co-localize with the mouse cluster. Human Cancer ME3 contains classical Cancer-Associated Fibroblasts (CAF) markers (*ACTA2, PDGFRA*), master Epithelial-Mesenchymal Transition (EMT) transcription factors (*ZEB1, SNAI2*), and critical extracellular matrix (ECM) remodelers (*FBN1, TIMP2, SERPINE1, MMP19*), suggesting that it is the canonical signature of CAFs and EMT. Simultaneously, Human Normal ME26 and ME61 target *CD34, FLT4, ERG, CLDN5, EGFL7*, and *PECAM1*, indicating they are robust endothelial cell and angiogenic programs. These co-clustered modules suggest that while the malignant epithelial cells in human ovarian cancer possess shattered genomes and utilize regulatory architectures that are completely divergent from those in murine models, the host microenvironmental response—specifically the supporting stroma and vasculature—remains deeply conserved. Thus, while standard murine models may fundamentally fail to recapitulate human epithelial rewiring, they may recapitulate the human tumor microenvironment. Consequently, preclinical murine studies should be carefully prioritized toward the development and testing of stroma- and angiogenesis-targeted therapies rather than epithelial-intrinsic mechanisms.

**Figure 4.**
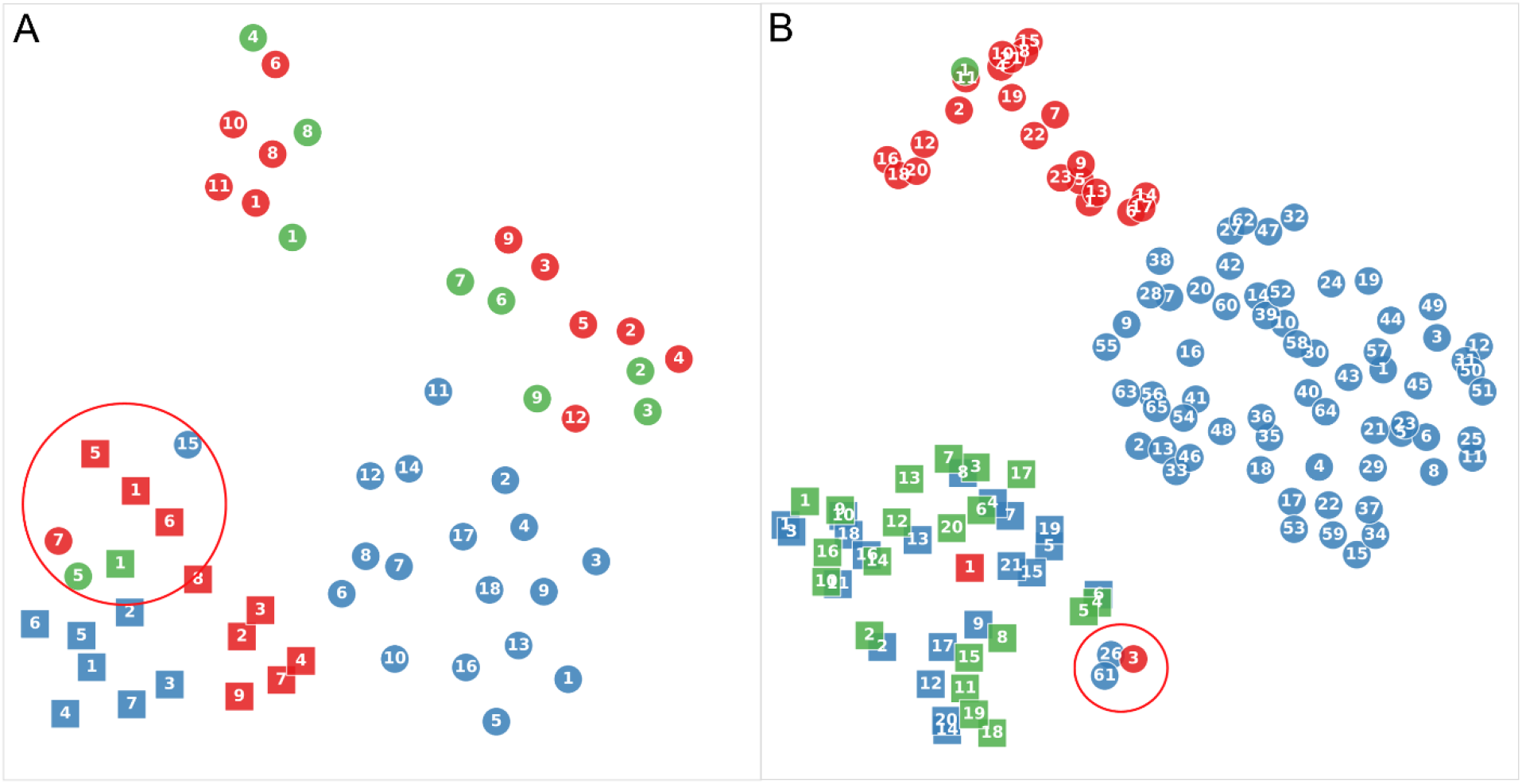
UMAP projection of kidney and ovarian cancer modules based on Jaccard distances between target gene sets. While human and mouse modules remain highly separated, several human modules were specifically co-clustered with mouse ones. Circular and square nodes map to human and mouse modules, blue, red, and green colors designate Normal, Cancer, and Preserved conditions, and numbers indicate modules. (A) Modules of kidney cancer. (B) Modules of ovarian cancer.

Together, the pathway-level hallmark landscape and gene-level Jaccard landscape provide objective, data-driven metrics for evaluating cross-species and cross-tumor differences. They demonstrate that while human and murine tumors share the same DEGs and cancer hallmarks, the underlying transcriptional regulation of these genes is highly divergent. This dual-landscape approach elucidates why certain cancer properties (e.g., *Sustaining proliferative signaling*) are more conserved across species than others (e.g., *Evading immune destruction*), and what modules are involved.

### 2.5 LS lncRNAs shape the tumor immune microenvironment

Immune cell infiltration is a critical determinant of tumor immune microenvironment (TIME) and heavily dictates cross-species discrepancies in cancer biology. Building on our initial observation of differential immune infiltration (Fig. 1E), we rigorously quantified these cross-species TIME discrepancies by deconvoluting bulk RNA-seq gene expression profiles using *CIBERSORTx* and *quanTIseq* (Finotello et al., 2019; Newman et al., 2019). The proportions of 10 major immune cell types across the 13 tumors indicate that immune infiltration exhibited significant species specificity in 54% of immune cell type-tumor type pairs (two-sided Mann-Whitney test, FDR < 0.01) (Fig. 5A).

**Figure 5.**
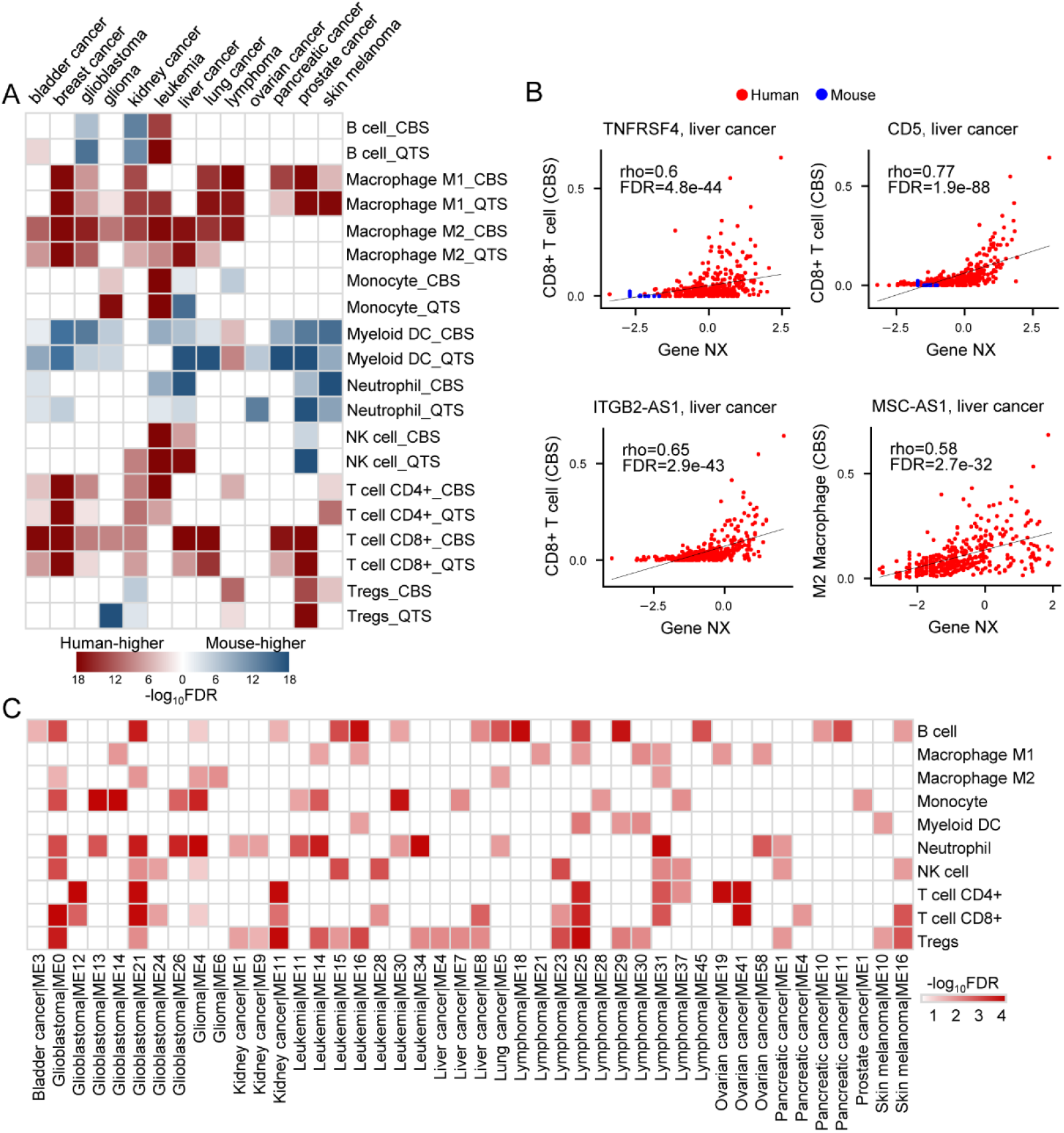
LS lncRNAs strongly correlate with TIME. (A) Heatmap depicting the enrichment of specific immune cell infiltration across 13 tumor types in both species. “_CBS” and “_QTS” denote measurements computationally extracted using *CIBERSORTx* and *quanTIseq*, respectively. (B) Scatter plots illustrating the direct correlations between the normalized expression of LS lncRNA-target pairs and the physical infiltration proportions of specific immune cells in liver cancer. Highlighted examples include *TNFRSF4* (an immune checkpoint gene in CD4+/CD8+ T cells) and *CD5* (whose expression correlates with increased CD8+ T cells penetration), plotted alongside *ITGB2-AS1* (previously implicated in regulating T and B cell activation) and *MSC-AS1* (which promotes hepatocellular carcinoma oncogenesis). (C) Heatmap demonstrating the enriched distribution of immune cell marker genes within 44 of the 54 ID-associated modules (hypergeometric distribution test, FDR < 0.1).

Next, we examined divergence at the module level by computationally isolating cross-species “divergent modules”—groups of functionally linked genes whose expression diverges between humans and mice— to systematically test whether LS lncRNAs drive tumor-immune discrepancies. First, for each tumor, we classified cross-species TDGs into modules using agglomerative hierarchical clustering, and subsequently, mapped LS lncRNAs to these groups (Supplementary Table 7). To identify directional regulatory relationships, we calculated a standard intra-species Spearman correlation (*ρ*) alongside an extended cross-species correlation metric (*ρ*′). *ρ*′>0 strictly indicates that the intra-species correlation (*ρ*) between an LS lncRNA and a TDG aligns logically with the cross-species transcriptional divergence of that TDG. This metric systematically links LS lncRNA activity to divergent gene expression. An LS lncRNA was formally assigned as a module regulator if its *ρ*′ values with internal TDGs were significantly greater than with external TDGs (one-sided Kolmogorov-Smirnov test, FDR < 10^−5^). Through this methodology, 74% of PS/SS lncRNAs and 49% of RS lncRNAs were assigned to at least one of the 299 modules, with PS/SS lncRNAs enriching a greater number of modules than RS lncRNAs. This bottom-up approach successfully recovered multiple experimentally validated LS lncRNA-TDG pairs from prior independent studies, including *AC011899.2*-*LPAR5* and *U62317.3*-*HLA-DRA* in human tumors, and *Gm42595*-*PPP2CB* in murine tumors (Jacob et al., 2021; Kremer et al., 2022; Liu and Hofman, 2022).

Then, we sought to directly connect these lncRNA-regulated modules to the actual physical imbalances observed in the TIME. We fitted a multiple linear regression model to test whether fluctuations in TDG expression within each module could mathematically predict shifts in the proportions of the 10 immune cell types. Ultimately, we identified 54 immune divergence-associated (ID-associated) modules based on three joint criteria: (a) robust predictive correlation (*R* > 0.5) with specific cell infiltrates, (b) significant cross-species divergence of at least one linked immune cell type, and (c) significant functional enrichment for the “*immune response*” (GO:0006955) GO term (FDR < 0.05). The 54 ID-associated modules demonstrate that LS lncRNAs strongly influence the recruitment or exclusion of immune cell infiltration across species, aligning with prior studies (Supplementary Tables 8, 9) (Kadian et al., 2024; Zhao et al., 2021).

Finally, we quantified this regulatory influence by computing Spearman correlations between expression profiles within each module and infiltrated immune cell proportions. Based on both deconvolution algorithms, 98% of target TDGs and 70% of assigned LS lncRNAs significantly correlated with the infiltration density of at least one immune cell type (FDR < 0.05) (Fig. 5B; Supplementary Fig. 5-1∼2). We further prioritized these regulatory elements by calculating an immune regulation (IR) score that estimates the overall likelihood that a specific LS lncRNA robustly governs an ID-associated module. Among the 273 LS lncRNAs ranking in the top 10% of IR scores (Supplementary Table 10), PS/SS lncRNAs governed significantly more modules on average than RS lncRNAs (two-sided Mann-Whitney test, p < 1.99×10^−45^). This disparity highlights an evolutionary expansion of the immunological regulatory repertoire specifically mediated by primate- and simian-specific lncRNAs (Fig. 5C).

### 2.6 The intrinsic links between LS lncRNAs, immune divergence, and immunotherapy efficacy

As an increasing number of anticancer therapies target the tumor immune response, we investigated whether LS lncRNAs modulate patient responses to pharmacological interventions. We analyzed RNA-seq clinical response data from the CTR-DB database, which encompasses 28 tumor types (including 10 of the 13 evaluated in our study) treated with 123 distinct therapeutic agents (Liu et al., 2022). Patients in each drug treatment cohort were stratified into clinical response and non-response groups, and genes were subsequently classified as differentially expressed (|log_2_FC| ≥ 1, FDR < 0.1) or non-differentially expressed between these groups. Within the CTR-DB dataset, 338 TDGs and 44 LS lncRNAs (derived from our previously identified 299 modules) exhibited significant differential expression associated directly with drug response. Notably, 78% of these drug resistance-associated TDGs and LS lncRNAs demonstrated significantly higher expression in the non-response patient cohorts (Supplementary Fig. 6-1; Supplementary Table 11). These findings indicate that transcriptional regulation by LS lncRNAs is strongly associated with clinical resistance to anticancer therapies.

To determine whether these clinical associations reflect a direct physical regulatory architecture rather than secondary statistical correlations, we examined DBSs for LS lncRNAs within promoters of TDGs in each ID-associated module. Of the 273 LS lncRNAs, 143 in humans and 101 in mice possessed strong DBSs (averaging ∼136 bp in length), and most DBSs (86% in humans and 80% in mice) overlapped with at least one ENCODE-annotated candidate cis-regulatory element (cCRE) (Encode Project Consortium et al., 2020). Mapping these DBSs revealed a highly disproportionate many-to-many regulatory topology— specific “hub” lncRNAs possessed DBSs across a vast array of TDGs, and specific “hub” targets harbored binding sites for a multitude of different LS lncRNAs. Crucially, the functional identities of these hub targets diverge profoundly across species. In humans, widely validated immune-regulatory and apoptotic hubs—such as *IFNAR1, CFLAR*, and *BTN2A2*—emerge as densely targeted nodes regulated by multiple LS lncRNAs. Conversely, the murine architecture centers its highest targeting density on completely distinct receptors and signaling molecules, such as *Il18r1* and *Notch1* (Supplementary Fig. 6-2).

To determine whether these observed cross-species differences in cancer immunity and therapeutic efficacy are strictly limited to the specific RNA-seq cohorts we analyzed, or are instead intrinsically hardwired in the genome, we evaluated the most extreme subset of entirely species-specific transcripts (28 human-specific [HS] and 39 mouse-specific [MS] lncRNAs) across the 54 ID-associated modules. We systematically compared the density of their predicted DBSs across TDGs annotated under the “*immune system process*” GO term (GO:0002376) versus a genome-wide background of all Ensembl-annotated PCGs. We found that genes within these specific immune-related GO terms are, on average, targeted by a significantly higher number of HS and MS lncRNAs compared to the genome-wide PCG background (one-sided Kolmogorov-Smirnov test, FDR < 0.05) (Fig. 6). These results suggest that the distinct regulation of immune programs by LS lncRNAs is physically encoded within the foundational genomic architecture of each species instead of merely an adaptive consequence of TIME.

**Figure 6.**
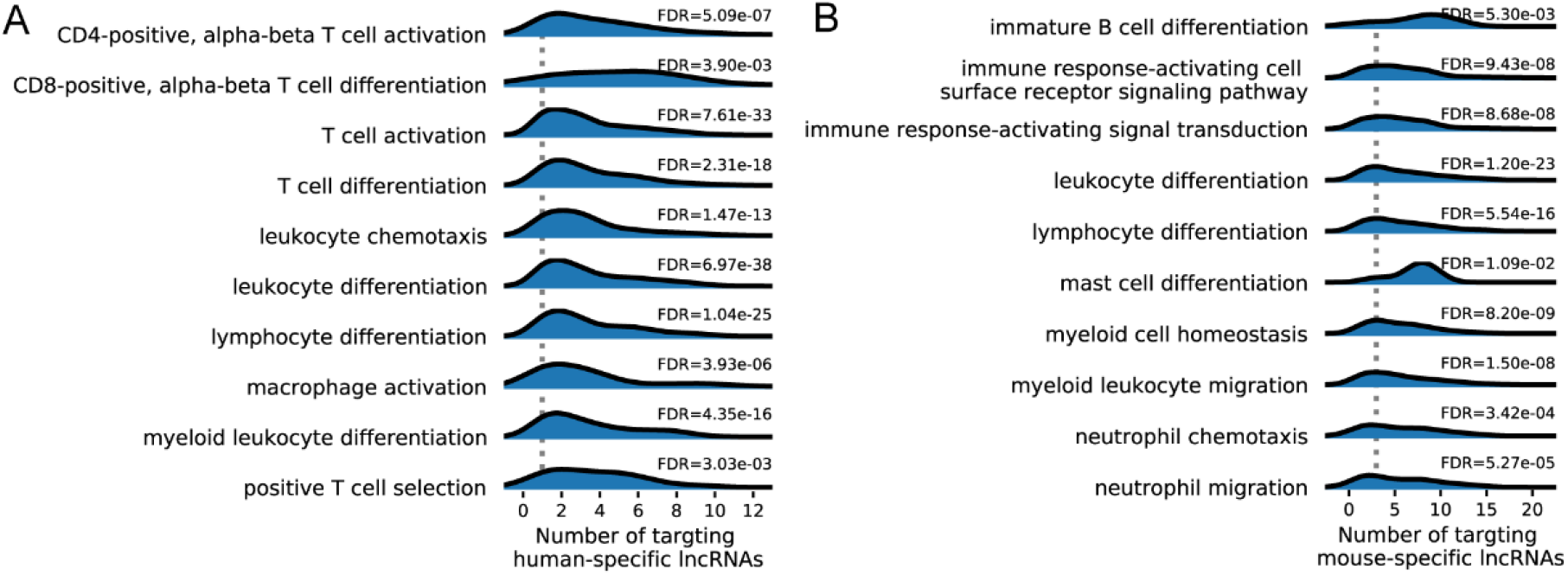
Immune-related genes are disproportionately targeted by LS lncRNAs. Density plots illustrate that downstream target genes heavily involved in immune-related GO terms are targeted by a greater volume of HS and MS lncRNAs than the general genome-wide PCG background. Panels display binding distribution data for (A) humans and (B) mice. X-axes represent the numbers of HS and MS lncRNAs that possess predicted DBSs within genes in the listed immune-related GO terms. Dashed vertical lines indicate the median numbers of targeting HS or MS lncRNAs calculated across the entire genome-wide PCG background.

## 3. Discussion

### 3.1 Methodological innovations

Recent studies have begun to examine lncRNAs as regulators and indicators of cancer hallmarks (Gutschner and Diederichs, 2012; Schmitt and Chang, 2016), particularly in the context of cancer immunity and immunotherapy resistance (Chiu et al., 2018; Guo et al., 2021; Iaccarino and Klapper, 2021; Raimondo et al., 2023; Winkle et al., 2021). However, pan-cancer analyses have mainly centered on DEGs and PCGs, emphasizing commonalities across tumor types (Cancer Genome Atlas Research et al., 2013; Hoadley et al., 2018). None has systematically examined LS lncRNAs, which greatly outnumber LS TFs and may more critically determine cross-species and cross-tumor gene expression differences.

The omission of physical regulatory evidence and the failure to account for many-to-many regulator-target relationships in transcriptional regulatory analysis have been seriously critiqued (Bar-Joseph et al., 2003; Saelens et al., 2018). While it is well established that mammalian genomes possess substantial LS lncRNAs—and multiple methods have been developed to predict lncRNA DBSs—whether transcriptome-wide analyses can be reliably conducted based on genome-wide prediction of LS lncRNA DBSs has remained unclear. Consequently, most transcriptomic analyses continue to infer transcriptional regulation solely from correlated expression and to detect disjoint modules using unsupervised clustering. This study combines lncRNA DBS targeting with correlated expression, systematically identifies modules comprising LS lncRNAs and target genes, and strictly distinguishes TDGs from traditional DEGs. Further, the identified modules were projected into UMAP manifolds, compared under three cellular conditions, and mapped directly to pathways and hallmarks. This framework helps reveal what transcription and signaling programs may be altered, invented, or hijacked via transcriptional regulation by LS lncRNAs. Examining the convergence of the two classes of manifolds perfectly captures the duality of animal models: what they fail at (the widely separated modules) and what they excel at (the tightly co-clustered modules). In support of these LS lncRNAs and DBSs, our companion study mapped the TE-derived origins of LS lncRNAs and their DBSs and investigated their cross-species effects in distinct biological contexts, including Alzheimer’s disease and spermatogenesis.

### 3.2 Conclusions, significance, and insights

Although human and mouse tumors broadly employ the same signaling pathways and display the same cancer hallmarks, LS lncRNAs can drive their underlying transcriptional regulation and molecular signaling to differ substantially. The dual-landscape representation provides an objective molecular framework for interpreting a substantial body of cancer research, evaluating preclinical mouse models, and identifying highly specific diagnostic and therapeutic targets. For example, the dense preservation of *Non-mutational epigenetic reprogramming* modules in glioblastoma and leukemia aligns with observations that these tumors survive by hijacking conserved neurodevelopmental and germinal center epigenetic programs (Wainwright and Scaffidi, 2017; Wu et al., 2025). Conversely, our mapped cross-species divergence in kidney and ovarian cancers functionally recapitulates known translational limitations in murine modules (Frew and Moch, 2015; Gower et al., 2025; Hsieh et al., 2017). By demonstrating that certain human tumors preserve modules based on extensive genomic instability, while their corresponding mouse models depend strictly on *Tumor-promoting inflammation*, our data provide a tangible structural explanation for why researchers observing success in mice often see distinct biological responses in humans. Conversely, co-clustered human and mouse modules in the Jaccard landscape clearly indicate the most highly translatable biological axes for robust therapeutic intervention.

Furthermore, our spatial UMAP landscapes generate multiple novel biological insights. First, the stark gain or loss of modules between Normal and Cancer conditions indicates a transition from functional complexity in normal cells toward a tight molecular convergence on dominant oncogenic axes. Second, tracking cross-species module concordance allows tumors to be categorized into two distinct archetypes: *Convergent*, where humans and mice share deep regulatory logic; and *Divergent*, where murine models achieve the same superficial hallmarks using entirely distinct LS lncRNAs. Third, examining sparse versus dense Preserved modules uncovers two fundamental etiologies of oncogenesis: “regulatory co-option” (where dense Preserved modules indicate tumors hijack and lock heavily altered developmental programs into a permanent active state) versus “topological obliteration and invention” (where an empty Preserved landscape indicates native regulatory networks are completely dismantled, forcing the invention of distinct, de novo Cancer modules to survive). Fourth, as Jaccard distance landscapes reveal, murine Normal and Cancer modules are cleanly separated in multiple tumors, whereas human modules intermingle; this reflects their distinctly different origins, mirroring how standard mouse models are generated via artificially forced mutations on homogeneous genetic backgrounds, as compared to the chaotic, naturally accumulated mutations intrinsic to human disease.

### 3.3 Broader significance and future directions

The analytical framework introduced here can readily be extended to additional human diseases, diverse animal species, and single-cell transcriptomics. Ultimately, our results suggest that translational studies must systematically evaluate the impact of LS lncRNA topology. For instance, specific cells possessing high developmental plasticity in mouse lung cancer are known to rapidly drive cancer progression and therapy resistance (Chan et al., 2026). In line with this, we identified significantly more accessible regulatory modules in the murine Cancer condition for lung cancer than in human lung cancer, directly offering a noncoding, mechanistic rationale for this murine plasticity.

Several important questions remain for future investigation. First, whether the cross-species differences mapped here are partially influenced by intrinsic differences in sample collection pipelines (e.g., TCGA vs. GTEx) warrants further verification using carefully matched, prospective cross-species datasets. Second, the therapeutic predictions computationally generated by our framework—such as targeting the LS lncRNA-RAD52 axis to disrupt human ovarian cancer, or utilizing *4930481A15Rik* expression as a regulatory biomarker in murine cohorts—must be advanced to preclinical validation. Finally, extending this module-centric methodology to incorporate other non-coding RNA classes (e.g., circular RNAs) will likely unmask further dimensions of cross-species divergence.

### 3.4 Limitations

Several technical limitations should be carefully considered. First, standard RNA-seq datasets prioritize polyadenylated transcripts, inherently underestimating the full repertoire and influence of non-polyadenylated lncRNAs. Second, TCGA and GTEx samples are not strictly paired, and adjacent normal tissues may not represent fully healthy states. Third, certain evaluated tumor types share the same normal reference samples, potentially introducing minor mathematical bias during contrastive analysis. Fourth, while highly stringent parameters were utilized to suppress computational false positives, alternative significance thresholds could inevitably yield partially different hierarchical module structures. Finally, as genome annotation improves, the catalog of LS lncRNAs will expand.

## 4. Materials and methods

### 4.1 Data resources and data pre-processing

Because cross-species transcriptomic comparisons are susceptible to technical artifacts and batch effects, we enforced a highly rigorous, standardized data processing pipeline across all sample cohorts. We retrieved sample annotations for publicly available murine experiments using the RNASeq-er API (24-June-2020 version) (Petryszak et al., 2017). The RNASeq-er API used a standardized pipeline to reassemble RNA-seq reads deposited on the ENA website, thereby eliminating batch effects caused by different data processing protocols. Samples from nude mice were excluded because of the lack of a normal immune system. Mouse RNA-seq data were further processed using the following steps to eliminate batch effects, remove outliers, and make gene expression comparable across humans and mice (Supplementary Fig. 1:1∼8). (1) Raw counts were normalized using the trimmed means of M values (TMM) and transcripts per million (TPM) methods to remove sequencing library-size biases prior to expression quantification (Abrams et al., 2019; Robinson and Oshlack, 2010). (2) Genes with TPM < 0.1 in over 80% of samples were excluded. (3) log-transformed TPM values were adjusted using the *ComBat* algorithm (Leek et al., 2012), taking “*experiment”* as the batch parameter to remove batch effects between experiments. (4) A median absolute deviation (MAD)-based method was further deployed to remove statistical outliers. Based on the *ComBat*-adjusted gene expression matrices, we computed the pairwise Pearson correlation distance (*d*_*ij*_) between samples *i* and *j* within each cancer and normal group using the *distance.pdist* function in the *SciPy* package. The isolation level (*IL*) of sample *i* was defined as the median of all pairwise distances between sample *i* and all other samples in that group (*IL*_*i*_ = median_*j*_ (*d*_*ij*_)). Letting 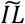 represents the overall median of all sample IL values within the group, the MAD was computed as: 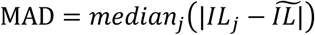. (5) Based on this MAD, an isolated score was computed for each sample: 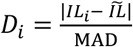. After removing extreme outliers (*D*>4), we retained only highly cohesive cohorts: specifically, 13 murine tumors with an effective sample size ≥20 and 11 corresponding normal tissues.

For human analysis, raw transcriptomic counts representing tumors and corresponding normal tissues were obtained from the UCSC Xena platform. This database has re-assembled the sequencing reads from the tumor-centric TCGA project with normal tissues from the GTEx project (Goldman et al., 2020; GTEx Consortium, 2015). Raw counts were processed using the same steps described above, including using the *ComBat* correction to remove potential non-biological differences between TCGA and GTEx datasets.

After preprocessing, 13 tumor types were selected, each comprising ≥20 tumor and ≥20 normal samples across both species (the sole exception was the human bladder normal group, n = 9). The finalized dataset encompassed 5,077 human tumor and 1,867 human normal samples, alongside 1,052 mouse tumor and 1,062 mouse normal samples (Fig. 1AB; Supplementary Table 1). The *ComBat*-adjusted gene expression of the 13 cancers and 11 normal tissues was normalized to z-scores (termed normalized expression, NX) using the *preprocessing.StandardScaler* function in the *scikit-learn* package (van den Berg et al., 2006). The 13 human and mouse cancers with abbreviated names are detailed in the supplementary file (Supplementary Table 1). Some cancer types (leukemia and lymphoma, glioma and glioblastoma) use the same normal samples as the control.

### 4.2 Data quality control

We evaluated whether factors, including individual characteristics (e.g., age and sex) and tumor characteristics (e.g., histological subtype, tumor evolution), may significantly induce transcriptomic variations between tumor samples by performing an Analysis of Similarities (ANOSIM)—a non-parametric test designed to evaluate differences between defined groups based on specific categorical covariates (Clarke, 1993). Briefly, for each tumor, we computed intra- and cross-species pairwise Euclidean distances based on orthologous gene NX values. For all ortholog pairs, these distances were converted into standardized ranks to compute the segregation index 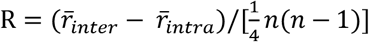. Here, 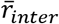 is the mean of cross-species ranks, 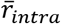 is the mean of intra-species ranks, *n* is the total sample size, and *R* indicates whether species divergence distances are significantly larger than internal cohort variance. To validate the robustness of *R*, we executed a 10,000-round permutation test. In each round, real sample labels were randomly shuffled between the human and murine groups to generate a mock baseline metric (R′). The probability of the randomized R′ values surpassing the biologically genuine R value was <10^−4^ for all tumors evaluated (Supplementary Fig. 1-3). This establishes that disparate intra-species covariates are negligible, and that the fundamental variation measured in this study is intrinsically driven by cross-species biology.

To empirically confirm that the 13 tumor sets differentiate cleanly according to biological identity, we used t-Distributed Stochastic Neighbor Embedding (t-SNE) projections. Post-PCA dimensionality reduction (reducing to 50 dimensions), the high-dimensional NX profiles distinctly clustered corresponding tumor and normal samples across species, verifying strong preprocessed signal integrity (Fig. 1C-D). Furthermore, to demonstrate that our divergence metrics successfully captured functionally distinct properties, we clustered the tumor samples explicitly using our top 100 computationally derived TDGs versus the top 100 TCGs using the *hierarchy.linkage* function (*method*=“ward”) in the *SciPy* package. The generated dendrograms clearly demonstrated fundamentally opposed spatial grouping, corroborating that these metrics identify biologically unique and opposite properties of inter-species tumor progression (Fig. 1A-B). Finally, mathematical similarities and divergences across modular gene sets were measured using classical Jaccard distances, which quantify the absolute scale of coordinate overlap.

### 4.3 Identify orthologous genes and define different kinds of genes

To prevent spurious transcript mapping or duplicated paralogs from confounding cross-species comparison, we identified one-to-one orthologous PCGs by extracting gene pairs from the Compara API on the EMBL-EBI website and downloading the list of human-mouse one-to-one orthologs from the MGI website (Supplementary Table 2, 3) (Bult et al., 2019; Vilella et al., 2009). The intersection of the two gene sets (from the EMBL-EBI and MGI websites) contains 14,988 pairs of orthologs.

Within-species DEGs were identified across the 13 tumors via the two-sided Mann-Whitney test based on |ΔNX = NX_cancer_ – NX_normal_| > 1 with FDR < 0.05 (Benjamini-Hochberg method), where NX_cancer_ – NX_normal_ reflects the medians derived from respective internal sample populations (Li et al., 2022). We also classified the human-mouse one-to-one orthologs into distinct categories based on their respective expression values within corresponding *cancer* populations (ΔNX = NX_human_cancer_ − NX_mouse_cancer_). Genes meeting stringent divergence (|ΔNX| > 1, FDR < 0.05) were categorized as Transcriptionally Divergent Genes (TDGs). Genes displaying moderate variance (0.5 ≤ |ΔNX| ≤ 1.0, FDR < 0.05) were classed as Transcriptionally Intermediate Genes (TIDs), while strictly stable orthologs (|ΔNX| < 0.5, FDR < 0.05) were categorized as functionally rigid Transcriptionally Conserved Genes (TCGs). Using these predefined definitions ensures high analytical reproducibility and clear stratification between core cancer homologies (TCGs) and lineage-adaptive biology (TDGs).

### 4.4 Identify LS lncRNAs

Some methods and databases detect cross-species lncRNA orthology relying on primary sequence conservation (Degalez et al., 2024; Pignatelli et al., 2016). The *Infernal* program performs RNA homology search using covariance models that integrate primary sequence and secondary structure information (Nawrocki and Eddy, 2013; Nawrocki et al., 2009). This method is more reliable because lncRNAs frequently exhibit rapid sequence turnover while conserving their secondary structure (Ulitsky, 2016), but it is also more time-consuming. *Infernal* has been used to construct the Rfam and Dfam databases.

We previously conducted exon-level searches for 13,562 GENCODE-annotated human lncRNA genes (GENCODE V18) across 16 representative mammalian genomes using *Infernal* (Lin et al., 2019). To minimize false orthology assignments from distant paralogs or random genome homologies, all exon searches were directionally constrained within syntenic regions predefined by UCSC pairwise genome alignments, with aligned boundaries extended at both ends to capture complete structural contexts. Using this identical methodology, we searched the orthologous sequences of the 4,396 GENCODE V36-updated human lncRNAs’ exons in these 16 mammals and the orthologous sequences of the 13,450 GENCODE-annotated mouse lncRNAs’ exons (GENCODE M22) across 4 selected reference genomes covering Muridae and Lagomorpha: rat, Ryukyu mouse (*Mus caroli*), shrew mouse (*Mus pahari*), and rabbit (as a non-rodent outgroup). To conservatively define true orthologous genes, we required that the number of identified orthologous exons in the target species must exceed 50% of the total exon count of the query human (or mouse) lncRNA gene. Human lncRNAs and mouse lncRNAs lacking any mammalian orthologs were defined as human-specific (HS) or mouse-specific (MS), respectively. Human lncRNAs with orthologues only in simians (chimpanzee, macaque, and marmoset), and with orthologues only in primates (simians plus tarsier and mouse lemur), were defined as simian-specific (SS) and primate-specific (PS), respectively. Mouse lncRNAs with orthologues only in rodents were defined as rodent-specific (RS). Homology and species-specificity of these LS lncRNAs were re-checked using UCSC *liftover*, and candidates disagreeing with our lineage-specific definitions were discarded. This pipeline identified 66 HS, 2006 SS, 2858 PS, 212 MS, and 3913 RS lncRNAs. LS lncRNAs with TPM>0.1 in at least 50% of samples of a tumor were considered robustly expressed in that tumor.

### 4.5 Predict lncRNAs’ DBDs and DBSs

ncRNA-DNA binding follows experimentally characterized rules governing specific base pairs to form RNA-DNA triplexes (Abu Almakarem et al., 2012), allowing lncRNAs to recruit epigenomic modification enzymes to binding sites (Chu et al., 2011; Engreitz et al., 2016; Yap et al., 2010). Multiple methods predict lncRNAs’ triplex-targeting sites (TTSs) using substring search based on individual rules (Buske et al., 2012; Kuo et al., 2019), but this approach has drawbacks: (a) predicted TTSs are short, disconnected, and (b) triplex-forming oligonucleotides (TFOs) are not identified simultaneously. Thus, TFOs and TTSs do not faithfully capture densely populated lncRNAs at binding sites, and their reliability needs additional statistical tests (Kuo et al., 2019). The *LongTarget* program takes another approach by integrating rules into 24 rule-sets that each cover all four kinds of nucleotides, translating the DNA sequence into 24 RNA sequences based on rule-sets, applying a variant of Smith–Waterman local alignment to 24 pairs of RNA sequences to identify all high-scoring local alignments, and identifying a set of densely overlapping triplexes (TFOs and TTSs) at each binding site as the DBD and DBS (He et al., 2015; Lin et al., 2019). Sequence alignment naturally tolerates the flexible distribution of sequence mismatches, insertions, and deletions that occur *in vivo*, identifies much longer DBDs and DBSs, and calculates the percentage of matched base pairs. We defined *binding affinity = Length* (length of local alignment) *× Identity* (percentage of matched base pairs) as a continuous metric for comparing lncRNA-DNA binding across lncRNAs and binding sites. All computational methods predict multiple TFOs or DBDs. *LongTarget* automatically ranks all DBDs upon predicted DBSs. Since DBD1 (the top-ranked DBD) generates substantially greater DBS coverage than other DBDs, which suggests statistically and biologically more reliable results, our downstream analyses were restricted to DBD1s and their DBSs. This method was validated using multiple methods. Especially, deleting DBD1 using CRISPR/Cas9 resulted in systemic transcriptomic alterations and measurable changes in cellular phenotype across distinct human cell lines (He et al., 2026; Lin et al., 2026). Benchmarking using simulated and real data also supports the reliability and biological relevance of predicted DBSs (He et al., 2015; Liu et al., 2017; Wen et al., 2022). The key drawback of excessive time consumption was effectively solved by the *Fasim-LongTarget* version (Wen et al., 2022).

For HS and MS lncRNAs, we used *LongTarget* (*Length* = 50, *Identity* = 60%) to predict DBSs within promoter regions (3,500 bp upstream to 1,500 bp downstream of the annotated transcription start site, TSS) across the entire annotated transcriptomes: 179,128 human transcripts (GRCh38.79) and 95,052 mouse transcripts (GRCm38.87). For broader PS/SS and RS lncRNA, we constrained DBS prediction using *Fasim-LongTarget* (*Length* = 50, *Identity* = 60%) to promoter regions (3,500 bp upstream to 1,500 bp downstream of TSS) of TDGs (5,593 in human and 5,882 in mice). These annotated TSSs in human and mouse genomes were obtained from the EPDnew website (Dreos et al., 2017).

### 4.6 Identify transcriptional regulatory modules

Expression levels (log_2_TPM) of DEGs in the 13 human tumors and 13 mouse tumors were organized into 26 matrices. These expression matrices, along with the two previously generated LS lncRNA-TDG DBS physical targeting matrices, served as inputs for the *eGRAM* program (*DBS affinity* ≥ 80, *Spearman correlation* ≥ 0.6, *module size* ≥ 50, *pathway enrichment significance (FDR)* ≤ 0.01, *overlap ratio* ≤ 80%) (He et al., 2026). Correlated gene expression was computed across three conditions: Normal (using only normal samples), Cancer (using only cancer samples), and Preserved (calculated as directionally consistent expression correlation observed across both normal and cancer contexts, thereby isolating interactions that robustly resist oncogenic state transitions). For each LS lncRNA (as the leading regulator), a correlation submatrix was extracted from the context-specific global gene–gene correlation matrix to identify a regulator set consisting of this LS lncRNA and its putative functional partners. A regulator set was identified by finding all maximal cliques in a correlation graph using the NetworkX implementation of the Bron–Kerbosch algorithm and collapsing maximal cliques into a single regulator by maximizing the mean absolute correlation.

To identify biologically rational modules, co-expression relationships were intersected with LS lncRNA-DBS targeting relationships, and final modules were defined as the intersection of expression-supported target correlations and physical DBS-supported targeting. Modules were iteratively merged until no two modules shared >80% overlapping genes. Three sets of modules were identified in the three conditions. Functional enrichment for genes in each module was calculated against KEGG and WikiPathway databases using a hypergeometric test. P-values were adjusted using the Benjamini–Hochberg method, and pathways with FDR < 0.01 were considered significantly enriched. Thus, each module has eight attributes: (a) ModuleID, (b) LncRNA, (c) Regulator set, (d) Target gene, (e) KEGG pathways, (f) Hit genes of enriched KEGG pathways, (g) WikiPathways, (h) Hit genes of enriched Wikipathways. Modules under the three conditions, along with enriched pathways, indicate reshaped or reconfigured (including gained, lost, and preserved) transcription and signaling.

### 4.7 Hallmark landscape analysis based on cancer hallmark mapping

To characterize the hallmark landscape of transcriptional regulatory modules across species, tumors, and conditions, we developed an algorithm to project modules into a unified UMAP manifold.

#### 4.7.1 Classification of KEGG pathways into cancer hallmarks

Enriched KEGG pathways (n = 313) in all modules identified by *eGRAM* were manually curated and classified into 11 cancer hallmarks (Hanahan, 2022, 2026). These hallmarks include: (a) *Sustaining proliferative signaling*, (b) *Resisting programmed cell death*, (c) *Inducing or accessing vasculature*, (d) *Unlocking phenotypic plasticity*, (e) *Activating invasion and metastasis*, (f) *Evading immune destruction*, (g) *Deregulating cellular metabolism*, (h) *Establishing replicative immortality*, (i) *Inactivating growth suppressors*, (j) *Non-mutational epigenetic reprogramming*, (k) *Tumor-promoting inflammation*.

#### 4.7.2 Determination of dominant hallmarks

We developed a quantitative scoring method based on pathway enrichment significance to assign a dominant hallmark to each module. For each pathway P in a module, an intensity score Score_P_ was computed as: Score_P_ = −log_10_(FDR_P_). Because pathways in a module may map multiple hallmarks, we aggregated pathway-level evidence for each hallmark H using the L2 norm (Euclidean magnitude): 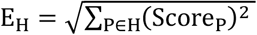. Applying the L2 norm ensures robust biological representation: it heavily weights highly significant pathways, but also accumulates weight from multiple moderate pathways supporting the same functional axis. Crucially, this mathematical structure rewards modules demonstrating broad, consistent pathway activation within a single hallmark, while preventing a single, statistically extreme outlier pathway from dominating the assignment in a way that pure linear summation might allow.

For each module, the hallmark scoring the highest E_H_ was designated as its hallmark, provided it passed two criteria: (a) *Absolute Significance*: The maximum E_H_ must exceed 1.3 (equivalent to a baseline FDR of 0.05), ensuring the assigned hallmark is supported by at least one definitively significant pathway or by the cumulative weight of multiple moderately significant pathways. (2) *Distinctiveness*: The difference between the top-scoring hallmark and the second-highest hallmark must exceed 0.1, ensuring unambiguous functional classification. Modules failing either threshold were labeled as *Unclassified* (functionally serving as a 12th baseline state in spatial mapping). There are three kinds of *Unclassified* modules: lacking enriched KEGG pathways, pathways could not be confidently assigned to these hallmarks, and no hallmark is significant. In all UMAP manifolds, the first two kinds are tightly clustered together.

#### 4.7.3 Manifold learning and visualization

To visualize functional relationships among modules, we constructed a weighted feature matrix in which rows represented transcriptional modules, columns represented KEGG pathways, and individual entries comprised the calculated ScoreP. Non-enriched pathways defaulted to 0. Dimensionality reduction was performed using Uniform Manifold Approximation and Projection (UMAP). To capture proportional similarities in pathway activation profiles regardless of sheer network size, we employed the Cosine distance metric (*n_neighbors* = 15, *min_dist* = 0.3, *metric* = cosine, *random_state* = 42). The embedding was computed across the integrated dataset encompassing all human and mouse modules across Normal, Cancer, and Preserved conditions. Crucially, while individual module nodes are colored by their mathematically assigned dominant hallmark, the topological projection is determined by the entire, unbiased vector of enriched pathways, enabling the visualization of polyfunctional and transitional regulatory states. If two modules are both Red (Proliferation), but one is located near the Brown (Inflammation) cluster, and the other is near the Purple (Metabolism) cluster, it means their *secondary* pathways are pulling them in different directions.

### 4.8 Module landscape analysis based on target gene mapping

Relying exclusively on predefined pathway databases to map landscapes introduces intrinsic knowledge bias. Specifically, highly novel, lineage-specific, or ultra-specialized modules often lack curated KEGG mapping, artificially swelling the *Unclassified* category and obscuring functional relationships. To map a high-resolution module landscape unconstrained by pre-curated definitions, we performed a second, purely data-driven manifold embedding based directly on the raw downstream target genes. We deployed UMAP using the Jaccard distance metric to quantify absolute compositional differences across the target gene sets of each module (*Parameters: n_neighbors = 7, min_dist = 0.25, metric = jaccard, random_state = 42*). Analogous to the hallmark manifold, modules across both species and all three biological conditions were projected into a single unified space where direct spatial proximity reflects raw regulatory homology. Projecting an unbiased gene-level landscape systematically addresses questions unanswerable by pathway-level heuristics, including how biologically related these modules are, regardless of species labels and condition, and how a module could be transformed from one condition to another or translated from one species to another.

### 4.9 Identify cross-species LS lncRNA and TDG modules

#### 4.9.1 Identifying cross-species TDG modules

To systematically investigate cross-species transcriptional divergence using a bottom-up statistical framework, we constructed integrated co-expression modules. First, to capture globally coordinated gene sets before explicitly assessing how lineage-specific regulators act upon them, we merged human and mouse TDG expression profiles into a unified, cross-species matrix for each tumor. Pairwise Spearman correlation coefficients (ρ) among these TDGs were computed using the *spearmanr* function in the *SciPy* package. Second, these correlation coefficients were converted into a standard distance metric (d = 1 − |ρ|), which served as the basis for constructing a binary dendrogram using *AgglomerativeClustering* in the *scikit-learn* package (with the *linkage* parameter set to “complete” to penalize loose connections and highly favor the identification of compact, highly cohesive functional modules. Third, modules were delineated by dynamically cutting the dendrogram at a stringent distance threshold corresponding to the 75th percentile of all pairwise distances. Candidate modules containing fewer than five genes were excluded to filter noise (notably, fewer than ten computationally derived modules were excluded across all tumors). Fourth, to rigorously validate the mathematical robustness of the remaining modules, permutation testing was executed over 10,000 iterations. The average pairwise correlation within each *N*-gene module was compared against the average correlation of *N* genes randomly sampled from all other modules. The empirical probability of a random background set exhibiting a higher average correlation than the average in any single module was <10^−4^, statistically confirming that these identified structures represent genuine, non-random co-expression programs. Across the 13 tumors, this robust procedure yielded 299 divergent TDG modules (ranging from 4 to 64 distinct modules per tumor).

#### 4.9.2 Adding LS lncRNAs to modules

We subsequently evaluated whether LS lncRNAs directionally regulate these integrated TDG modules. First, for each LS lncRNA and all TDGs within a specific species and tumor, we computed a standard intra-species Spearman correlation (ρ). Second, to accurately evaluate whether this intra-species correlated expression is associated with inter-species expression divergence, we defined an extended correlation metric, *ρ*′. Specifically, *ρ*′ = *ρ* if the TDG exhibited a higher relative expression in the primary species examined compared to its counterpart outgroup species; conversely, *ρ*′ = −*ρ* if the TDG exhibited lower relative expression. Consequently, a positive extended correlation (*ρ*′ > 0) implies a concerted, directional regulatory alignment. For example, if an LS lncRNA is positively correlated with a TDG (*ρ* > 0) and that specific TDG is concurrently upregulated in the same species relative to the other, the positive *ρ*′ metric mathematically indicates that the intra-species regulation directly parallels the cross-species regulation. Conversely, a *ρ*′ < 0 indicates that the intra-species regulation contradicts the cross-species divergence. Third, to formally and statistically assign an LS lncRNA to a defined TDG module, we compared the distribution of its *ρ*′ values with TDGs internal to the module against its *ρ*′ values with all TDGs external to the module using a one-sided Kolmogorov-Smirnov (KS) test (*ks_2samp* from *SciPy*). P-values were adjusted for multiple hypothesis testing using the Benjamini-Hochberg method. An LS lncRNA was assigned to a module only if it met a stringent FDR threshold of <10^−5^. While highly conservative, applying this harsh threshold forcefully minimizes computational false positives and ensures high-confidence regulatory mapping. Several LS lncRNAs satisfied these criteria across multiple discrete modules, indicating pleiotropic regulatory roles spanning distinct transcriptional programs and tumor contexts.

### 4.10 Analyze immune cell infiltration

To objectively quantify TIME and avoid algorithm-specific biases, we used both the *CIBERSORTx* and *quanTIseq* programs (implemented via the TIMER 2.0 platform) to estimate the fractional proportions of 10 major immune cell types by deconvolving bulk gene expression data (Finotello et al., 2019; Newman et al., 2019). Baseline comparisons of specific immune cell infiltration proportions between human and mouse tumors were conducted using the two-sided Mann–Whitney test, with P-values adjusted for multiple testing using the Benjamini-Hochberg method.

After defining the baseline TIME profiles, we investigated the specific contribution of each TDG and LS lncRNA within each module to overall immune cell infiltration using Spearman correlation testing. Specifically, this tested the correlation between the distinct expression of a TDG (and LS lncRNA) and the resulting deconvoluted proportion of each infiltrated immune cell type. Because TDGs are defined by 1:1 orthology, they were evaluated jointly using merged human and mouse data to establish overarching cross-species infiltration trends. Conversely, because LS lncRNAs are inherently lineage-specific, their correlation to immune infiltration was tested exclusively within their native human or mouse datasets. All corresponding P-values were adjusted to FDR using the Benjamini-Hochberg method.

Finally, to validate the functional identity of the constructed modules, the overarching enrichment of established immune cell markers was computed utilizing a hypergeometric overlap test. This evaluated the intersection between all assigned TDGs within a module and known expression markers for each respective immune cell type. Annotated reference marker genes were sourced directly from the CellMarker database (Zhang et al., 2019).

### 4.11 Identify ID-associated modules and ID-associated LS lncRNAs

Because many TDGs relate to immune biology (section 2.1), we established a systematic statistical pipeline to rigorously define specifically immune divergence-associated (ID-associated) modules based on our initial ρ′-based TDG clusters. First, recognizing that the tumor immune microenvironment is recruited by coordinated genetic programs rather than isolated transcripts, we constructed a multiple linear regression model linking all TDGs within a cross-species module to the estimated proportions of infiltrated immune cells (utilizing the *linear_model.LinearRegression* function within the *scikit-learn* package).

Second, we computed a multiple correlation coefficient (R), defined as the square root of the model’s determination coefficient. This continuous *R* coefficient ranges from 0 to 1 and quantitatively measures how effectively fluctuations in defined cell proportions can be predicted by the linear combination of module-specific TDG expression across species. To ensure stability, a mean *R* was finalized by averaging the two independent *R* values generated from the separate *CIBERSORTx* and *quanTIseq* data streams. Third, we computationally verified whether a module was fundamentally rooted in immune biology using the hypergeometric overlap test, measuring the intersection between module TDGs and genes annotated to the “*immune response”* GO term (GO:0006955). A module was designated as an ID-associated module only if it fulfilled three stringent, joint criteria: (a) the multiple correlation coefficient R relative to at least one immune cell type was >0.5, demonstrating high predictive power; (b) at least one immune cell type demonstrated a significantly different proportion of baseline infiltration across species; and (c) the TDGs comprising the module demonstrated significant enrichment for the aforementioned “*immune response”* GO term (FDR < 0.05).

Fourth, to prioritize the specific LS lncRNAs driving these immunomodulatory programs, we formulated an immune regulation (IR) score. This metric systematically calculates the likelihood that a specific LS lncRNA heavily regulates an ID-associated module by multiplying the statistical confidence of the module assignment by the maximum phenotypic effect size of that module on the TIME. For each LS lncRNA *m* mapped to an ID-associated module *j*, the *IR* score was derived as: *IR*_*m,j*_ = − *log*_10_ (*FDR*_*m,j*_) ^*^ *max*_*i*_(*R*_*ij*_). Here, the FDR_m,j_ value reflects the statistical significance from the Kolmogorov-Smirnov test for the correlation between the expression of the LS lncRNA and its downstream TDGs within module *j*. The metric *max*_*i*_(*R*_*ij*_)selects the maximum multiple correlation coefficient recorded between module j and the available array of ten defined immune cell type *i*. Fifth, under the assumption that central regulatory lncRNAs might execute immunomodulation pleiotropically across disparate cancer types, we calculated a comprehensive Summary IR (SIR) score for each LS lncRNA *m* aggregated across the 13 tumors: 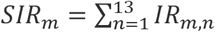. Inn this summation, if the LS lncRNA *m* was absent from all ID-associated modules within tumor *n*, the *IR*_*m,n*_ naturally defaulted to 0. Conversely, if LS lncRNA *m* populated multiple ID-associated modules within the same tumor *n*, the maximal local IR value was selectively adopted to represent that tumor space. Ultimately, LS lncRNAs that generated a finalized SIR score within the top 10% of the dataset were formally classified as broad ID-associated lncRNAs capable of potent cross-tumor immunological influence (Supplementary Table 8).

### 4.12 Gene set enrichment analysis

To fundamentally characterize the distinct biological processes governed by cross-species divergence versus cross-species conservation, we first performed baseline functional over-representation analysis on the broad sets of TDGs and TCGs utilizing the *g:Profiler* program queried against the Gene Ontology (GO) database (Fig. 1E) (Gene Ontology Consortium, 2021; Raudvere et al., 2019).

To examine whether target genes of the 28 HS lncRNAs and 39 MS lncRNAs are more enriched for immune-related genes than genome-wide PCGs, we used the *Gorilla* program to perform gene set enrichment analysis (Eden et al., 2009). Because a functionally important regulation is often reflected by its prevalence across multiple tumors in the pan-cancer setting, we structured this as a ranked-list analysis. For each PCG in the 54 ID-associated modules, we calculated its cross-tumor prevalence by counting the total number of tumors in which that gene emerged as differentially expressed. Genes were then ranked in descending order based on this tumor count. By taking these explicitly ranked lists of human and mouse genes as inputs, *GOrilla* identified significantly enriched GO terms (FDR < 0.05) weighted toward the top of the distribution, thereby selectively highlighting core immune pathways that are not only lineage-specific but act robustly as pleiotropic hubs across diverse tumor microenvironments.

### 4.13 Drug resistance analysis

To critically assess the clinical relevance of our comparative framework, we sought to determine whether the lineage-specific regulatory divergence we mapped actually dictates human therapeutic failure *in vivo*. To achieve this, we accessed patient-derived clinical transcriptomic profiles and their corresponding therapeutic response metadata from the CTR-DB database (Liu et al., 2022). This comprehensive dataset encompasses 28 distinct cancer types (including 10 of the 13 tumor types directly evaluated in our dual-landscape analysis) and spans patient responses to 123 distinct therapeutic agents, including chemotherapy, targeted therapy, and modern immunotherapy.

Within each specific drug treatment cohort, patients were rigorously stratified into clinical “response” and “non-response” groups. Transcripts putatively associated with drug resistance were identified by calculating DEGs between these clinical response categories. Given the characteristically high baseline expression variance inherent to real-world patient cohorts, a practical statistical threshold of |log_2_FC| ≥ 1 and FDR < 0.1 was applied to isolate resistance markers. Finally, we systematically intersected these independently verified clinical drug resistance genes with our defined catalog of clade-specific LS lncRNAs and TDGs. This intersection enabled us to objectively verify whether the transcriptionally divergent networks identified by our computational platform inherently harbor or directly modulate clinically relevant resistance pathways.

## Authors’ contributions

H.Z. and J.L. designed the study. H.Z. and J.L. developed the *eGRAM* program, H.Z. performed *eGRAM* and UMAP analyses, J.L. performed all other analyses, and X.L. performed drug efficacy analyses. H.Z. drafted the manuscript. All authors have read the manuscript and consent to its publication.

### Additional information

Supplementary file 1: Supplementary figures.

Supplementary file 2: Supplementary tables.

## References

Abrams, Z.B., Johnson, T.S., Huang, K., Payne, P.R.O., and Coombes, K. (2019). A protocol to evaluate RNA sequencing normalization methods. BMC Bioinformatics 20, 679.

Abu Almakarem, A.S., Petrov, A.I., Stombaugh, J., Zirbel, C.L., and Leontis, N.B. (2012). Comprehensive survey and geometric classification of base triples in RNA structures. Nucleic Acids Res 40, 1407–1423.

Arrowsmith, J., and Miller, P. (2013). Trial watch: phase II and phase III attrition rates 2011-2012. Nat Rev Drug Discov 12, 569.

Banerji, C.R., Miranda-Saavedra, D., Severini, S., Widschwendter, M., Enver, T., Zhou, J.X., and Teschendorff, A.E. (2013). Cellular network entropy as the energy potential in Waddington’s differentiation landscape. Sci Rep 3, 3039.

Bar-Joseph, Z., Gerber, G.K., Lee, T.I., Rinaldi, N.J., Yoo, J.Y., Robert, F., Gordon, D.B., Fraenkel, E., Jaakkola, T.S., Young, R.A., et al. (2003). Computational discovery of gene modules and regulatory networks. Nat Biotechnol 21, 1337–1342.

Becker, M., Nassar, H., Espinosa, C., Stelzer, I.A., Feyaerts, D., Berson, E., Bidoki, N.H., Chang, A.L., Saarunya, G., Culos, A., et al. (2023). Large-scale correlation network construction for unraveling the coordination of complex biological systems. Nat Comput Sci 3, 346–359.

Bradner, J.E., Hnisz, D., and Young, R.A. (2017). Transcriptional Addiction in Cancer. Cell 168, 629–643.

Breschi, A., Gingeras, T.R., and Guigo, R. (2017). Comparative transcriptomics in human and mouse. Nat Rev Genet 18, 425–440.

Bult, C.J., Blake, J.A., Smith, C.L., Kadin, J.A., Richardson, J.E., and Mouse Genome Database, G. (2019). Mouse Genome Database (MGD) 2019. Nucleic Acids Res 47, D801–D806.

Buske, F.A., Bauer, D.C., Mattick, J.S., and Bailey, T.L. (2012). Triplexator: detecting nucleic acid triple helices in genomic and transcriptomic data. Genome Res 22, 1372–1381.

Cancer Genome Atlas Research, N. (2011). Integrated genomic analyses of ovarian carcinoma. Nature 474, 609–615.

Cancer Genome Atlas Research, N., Weinstein, J.N., Collisson, E.A., Mills, G.B., Shaw, K.R., Ozenberger, B.A., Ellrott, K., Shmulevich, I., Sander, C., and Stuart, J.M. (2013). The Cancer Genome Atlas Pan-Cancer analysis project. Nat Genet 45, 1113–1120.

Chan, J.E., Pan, C.H., Rub, J., Guzman, G., Krause, K., Brown, E., Zhang, Z., Styers, H., Hartmann, G., Li, Z., et al. (2026). Critical role for a high-plasticity cell state in lung cancer. Nature 651, 231–241.

Cheng, Y., Ma, Z., Kim, B.H., Wu, W., Cayting, P., Boyle, A.P., Sundaram, V., Xing, X., Dogan, N., Li, J., et al. (2014). Principles of regulatory information conservation between mouse and human. Nature 515, 371–375.

Chiu, H.S., Somvanshi, S., Patel, E., Chen, T.W., Singh, V.P., Zorman, B., Patil, S.L., Pan, Y., Chatterjee, S.S., Cancer Genome Atlas Research, N., et al. (2018). Pan-Cancer Analysis of lncRNA Regulation Supports Their Targeting of Cancer Genes in Each Tumor Context. Cell Rep 23, 297–312 e212.

Chu, C., Qu, K., Zhong, F.L., Artandi, S.E., and Chang, H.Y. (2011). Genomic maps of long noncoding RNA occupancy reveal principles of RNA-chromatin interactions. Mol Cell 44, 667–678.

Clarke, K.R. (1993). Non-parametric multivariate analyses of changes in community structure. Australian Journal of Ecology 18, 117–143.

Day, C.P., Merlino, G., and Van Dyke, T. (2015). Preclinical mouse cancer models: a maze of opportunities and challenges. Cell 163, 39–53.

Degalez, F., Allain, C., Lagoutte, L., Lecerf, P., and Lagarrigue, P. (2024). Cross-species orthology detection of long non-coding RNAs (lncRNA) through 13 species using genomic and functional annotations. bioRxiv.

Derrien, T., Johnson, R., Bussotti, G., Tanzer, A., Djebali, S., Tilgner, H., Guernec, G., Martin, D., Merkel, A., Knowles, D.G., et al. (2012). The GENCODE v7 catalog of human long noncoding RNAs: analysis of their gene structure, evolution, and expression. Genome Res 22, 1775–1789.

Dreos, R., Ambrosini, G., Groux, R., Cavin Perier, R., and Bucher, P. (2017). The eukaryotic promoter database in its 30th year: focus on non-vertebrate organisms. Nucleic Acids Res 45, D51–D55.

Easwaran, H., Tsai, H.C., and Baylin, S.B. (2014). Cancer epigenetics: tumor heterogeneity, plasticity of stem-like states, and drug resistance. Mol Cell 54, 716–727.

Eden, E., Navon, R., Steinfeld, I., Lipson, D., and Yakhini, Z. (2009). GOrilla: a tool for discovery and visualization of enriched GO terms in ranked gene lists. BMC Bioinformatics 10, 48.

Eichhorn, J.H., and Scully, R.E. (1995). “Adenoid cystic” and basaloid carcinomas of the ovary: evidence for a surface epithelial lineage. A report of 12 cases. Mod Pathol 8, 731–740.

Encode Project Consortium, Moore, J.E., Purcaro, M.J., Pratt, H.E., Epstein, C.B., Shoresh, N., Adrian, J., Kawli, T., Davis, C.A., Dobin, A., et al. (2020). Expanded encyclopaedias of DNA elements in the human and mouse genomes. Nature 583, 699–710.

Engreitz, J.M., Haines, J.E., Perez, E.M., Munson, G., Chen, J., Kane, M., McDonel, P.E., Guttman, M., and Lander, E.S. (2016). Local regulation of gene expression by lncRNA promoters, transcription and splicing. Nature 539, 452–455.

Ferrando, A.A., and Lopez-Otin, C. (2017). Clonal evolution in leukemia. Nat Med 23, 1135–1145.

Finotello, F., Mayer, C., Plattner, C., Laschober, G., Rieder, D., Hackl, H., Krogsdam, A., Loncova, Z., Posch, W., Wilflingseder, D., et al. (2019). Molecular and pharmacological modulators of the tumor immune contexture revealed by deconvolution of RNA-seq data. Genome Med 11, 34.

Flavahan, W.A., Gaskell, E., and Bernstein, B.E. (2017). Epigenetic plasticity and the hallmarks of cancer. Science 357.

Frew, I.J., and Moch, H. (2015). A clearer view of the molecular complexity of clear cell renal cell carcinoma. Annu Rev Pathol 10, 263–289.

Fridman, W.H., Zitvogel, L., Sautes-Fridman, C., and Kroemer, G. (2017). The immune contexture in cancer prognosis and treatment. Nat Rev Clin Oncol 14, 717–734.

Gene Ontology Consortium (2021). The Gene Ontology resource: enriching a GOld mine. Nucleic Acids Res 49, D325–D334.

Goldman, M.J., Craft, B., Hastie, M., Repecka, K., McDade, F., Kamath, A., Banerjee, A., Luo, Y., Rogers, D., Brooks, A.N., et al. (2020). Visualizing and interpreting cancer genomics data via the Xena platform. Nat Biotechnol 38, 675–678.

Gower, A., Win, S., Ganguly, R., Johnson, M., Velez, M.A., Cummings, A.L., Lisberg, A., Garon, E.B., and Di Carlo, B. (2025). Identification of Targetable EGFR Mutations in Ovarian Cancer. JCO Precis Oncol 9, e2500390.

GTEx Consortium (2015). Human genomics. The Genotype-Tissue Expression (GTEx) pilot analysis: multitissue gene regulation in humans. Science 348, 648–660.

Guo, W., Wang, Y., Yang, M., Wang, Z., Wang, Y., Chaurasia, S., Wu, Z., Zhang, M., Yadav, G.S., Rathod, S., et al. (2021). LincRNA-immunity landscape analysis identifies EPIC1 as a regulator of tumor immune evasion and immunotherapy resistance. Sci Adv 7.

Gutschner, T., and Diederichs, S. (2012). The hallmarks of cancer: a long non-coding RNA point of view. RNA Biol 9, 703–719.

Haemmerle, M., and Gutschner, T. (2015). Long non-coding RNAs in cancer and development: where do we go from here? Int J Mol Sci 16, 1395–1405.

Hanahan, D. (2022). Hallmarks of Cancer: New Dimensions. Cancer Discov 12, 31–46.

Hanahan, D. (2026). Hallmarks of cancer - Then and now, and behond. Cell 189.

He, S., Xiong, W., Huo, J., Lin, J., Li, J., and Zhu, H. (2026). Substantial unannotated noncoding transcripts in tumors may transcriptionally regulate cancer-related genes. BMC Biol 24.

He, S., Zhang, H., Liu, H., and Zhu, H. (2015). LongTarget: a tool to predict lncRNA DNA-binding motifs and binding sites via Hoogsteen base-pairing analysis. Bioinformatics 31, 178–186.

Hegde, P.S., and Chen, D.S. (2020). Top 10 Challenges in Cancer Immunotherapy. Immunity 52, 17–35.

Ho, A.S., Kannan, K., Roy, D.M., Morris, L.G., Ganly, I., Katabi, N., Ramaswami, D., Walsh, L.A., Eng, S., Huse, J.T., et al. (2013). The mutational landscape of adenoid cystic carcinoma. Nat Genet 45, 791–798.

Hoadley, K.A., Yau, C., Hinoue, T., Wolf, D.M., Lazar, A.J., Drill, E., Shen, R., Taylor, A.M., Cherniack, A.D., Thorsson, V., et al. (2018). Cell-of-Origin Patterns Dominate the Molecular Classification of 10,000 Tumors from 33 Types of Cancer. Cell 173, 291–304 e296.

Hodge, R.D., Bakken, T.E., Miller, J.A., Smith, K.A., Barkan, E.R., Graybuck, L.T., Close, J.L., Long, B., Johansen, N., Penn, O., et al. (2019). Conserved cell types with divergent features in human versus mouse cortex. Nature 573, 61–68.

Hsieh, J.J., Purdue, M.P., Signoretti, S., Swanton, C., Albiges, L., Schmidinger, M., Heng, D.Y., Larkin, J., and Ficarra, V. (2017). Renal cell carcinoma. Nat Rev Dis Primers 3, 17009.

Huarte, M. (2015). The emerging role of lncRNAs in cancer. Nat Med 21, 1253–1261.

Iaccarino, I., and Klapper, W. (2021). LncRNA as Cancer Biomarkers. Methods Mol Biol 2348, 27–41.

Jacob, S., Jurinovic, V., Lampert, C., Pretzsch, E., Kumbrink, J., Neumann, J., Haoyu, R., Renz, B.W., Kirchner, T., Guba, M.O., et al. (2021). The association of immunosurveillance and distant metastases in colorectal cancer. J Cancer Res Clin Oncol 147, 3333–3341.

Jonasch, E., Gao, J., and Rathmell, W.K. (2014). Renal cell carcinoma. BMJ 349, g4797.

Kadian, L.K., Verma, D., Lohani, N., Yadav, R., Ranga, S., Gulshan, G., Pal, S., Kumari, K., and Chauhan, S.S. (2024). Long non-coding RNAs in cancer: multifaceted roles and potential targets for immunotherapy. Mol Cell Biochem 479, 3229–3254.

Khokhlova, S., Alnaqqash, M.A., Bahaj, W., Bujassoum, S., Lee, J., Mokhtar, M., Tyulyandina, A., Vargas Malaga, C.L., and Wu, C.H. (2025). Prevalence of homologous recombination deficiency among women with newly diagnosed ovarian, primary peritoneal, and/or fallopian tube cancer: the international HALO study. Int J Gynecol Cancer 35, 101645.

Konstantinopoulos, P.A., Ceccaldi, R., Shapiro, G.I., and D’Andrea, A.D. (2015). Homologous Recombination Deficiency: Exploiting the Fundamental Vulnerability of Ovarian Cancer. Cancer Discov 5, 1137–1154.

Kremer, K.N., Buser, A., Thumkeo, D., Narumiya, S., Jacobelli, J., Pelanda, R., and Torres, R.M. (2022). LPA suppresses T cell function by altering the cytoskeleton and disrupting immune synapse formation. Proc Natl Acad Sci U S A 119, e2118816119.

Krivtsov, A.V., Twomey, D., Feng, Z., Stubbs, M.C., Wang, Y., Faber, J., Levine, J.E., Wang, J., Hahn, W.C., Gilliland, D.G., et al. (2006). Transformation from committed progenitor to leukaemia stem cell initiated by MLL-AF9. Nature 442, 818–822.

Kuo, C.C., Hanzelmann, S., Senturk Cetin, N., Frank, S., Zajzon, B., Derks, J.P., Akhade, V.S., Ahuja, G., Kanduri, C., Grummt, I., et al. (2019). Detection of RNA-DNA binding sites in long noncoding RNAs. Nucleic Acids Res 47, e32.

Laws, M.J., Kannan, A., Pawar, S., Haschek, W.M., Bagchi, M.K., and Bagchi, I.C. (2014). Dysregulated estrogen receptor signaling in the hypothalamic-pituitary-ovarian axis leads to ovarian epithelial tumorigenesis in mice. PLoS Genet 10, e1004230.

Leek, J.T., Johnson, W.E., Parker, H.S., Jaffe, A.E., and Storey, J.D. (2012). The sva package for removing batch effects and other unwanted variation in high-throughput experiments. Bioinformatics 28, 882–883.

Li, Y., Ge, X., Peng, F., Li, W., and Li, J.J. (2022). Exaggerated false positives by popular differential expression methods when analyzing human population samples. Genome Biol 23, 79.

Lin, J., Wen, Y., He, S., Yang, X., Zhang, H., and Zhu, H. (2019). Pipelines for cross-species and genome-wide prediction of long noncoding RNA binding. Nat Protoc 14, 795–818.

Lin, J., Wen, Y., Tang, J., Zhang, X., Zhang, H., and Zhu, H. (2026). Human-specific lncRNAs contributed critically to human evolution by distinctly regulating gene expression. Elife 12.

Liu, D., and Hofman, P. (2022). Expression of NOTCH1, NOTCH4, HLA-DMA and HLA-DRA is synergistically associated with T cell exclusion, immune checkpoint blockade efficacy and recurrence risk in ER-negative breast cancer. Cell Oncol (Dordr).

Liu, H., Shang, X., and Zhu, H. (2017). LncRNA/DNA binding analysis reveals losses and gains and lineage specificity of genomic imprinting in mammals. Bioinformatics 33, 1431–1436.

Liu, S.J., Dang, H.X., Lim, D.A., Feng, F.Y., and Maher, C.A. (2021). Long noncoding RNAs in cancer metastasis. Nat Rev Cancer 21, 446–460.

Liu, Z., Liu, J., Liu, X., Wang, X., Xie, Q., Zhang, X., Kong, X., He, M., Yang, Y., Deng, X., et al. (2022). CTR-DB, an omnibus for patient-derived gene expression signatures correlated with cancer drug response. Nucleic Acids Res 50, D1184–D1199.

Mattick, J.S., Amaral, P.P., Carninci, P., Carpenter, S., Chang, H.Y., Chen, L.L., Chen, R., Dean, C., Dinger, M.E., Fitzgerald, K.A., et al. (2023). Long non-coding RNAs: definitions, functions, challenges and recommendations. Nat Rev Mol Cell Biol 24, 430–447.

Mestas, J., and Hughes, C.C. (2004). Of mice and not men: differences between mouse and human immunology. J Immunol 172, 2731–2738.

Miras, I., Vazquez-Gutierrez, I., Estevez-Garcia, P., and Munoz-Galvan, S. (2025). DNA repair pathways in ovarian cancer: Implications for therapy and resistance. Biomed Pharmacother 193, 118719.

Nawrocki, E.P., and Eddy, S.R. (2013). Computational identification of functional RNA homologs in metagenomic data. RNA Biol 10, 1170–1179.

Nawrocki, E.P., Kolbe, D.L., and Eddy, S.R. (2009). Infernal 1.0: inference of RNA alignments. Bioinformatics 25, 1335–1337.

Newman, A.M., Steen, C.B., Liu, C.L., Gentles, A.J., Chaudhuri, A.A., Scherer, F., Khodadoust, M.S., Esfahani, M.S., Luca, B.A., Steiner, D., et al. (2019). Determining cell type abundance and expression from bulk tissues with digital cytometry. Nat Biotechnol 37, 773–782.

Olson, B., Li, Y., Lin, Y., Liu, E.T., and Patnaik, A. (2018). Mouse Models for Cancer Immunotherapy Research. Cancer Discov 8, 1358–1365.

Ota, Y., Gupta, V., Fashemi, B.V., Akande, M., Babu, P., Thuthika, P., Elizagaray, M., Sun, L., Sanders, B., Kuroki, L.M., et al. (2025). Targeting RAD52 overcomes PARP inhibitor resistance in preclinical Brca2-deficient ovarian cancer model. bioRxiv.

Petryszak, R., Fonseca, N.A., Fullgrabe, A., Huerta, L., Keays, M., Tang, Y.A., and Brazma, A. (2017). The RNASeq-er API-a gateway to systematically updated analysis of public RNA-seq data. Bioinformatics 33, 2218–2220.

Pignatelli, M., Vilella, A.J., Muffato, M., Gordon, L., White, S., Flicek, P., and Herrero, J. (2016). ncRNA orthologies in the vertebrate lineage. Database (Oxford) 2016.

Quesada, S., Penault-Llorca, F., Matias-Guiu, X., Banerjee, S., Barberis, M., Coleman, R.L., Colombo, N., DeFazio, A., McNeish, I.A., Nogueira-Rodrigues, A., et al. (2025). Homologous recombination deficiency in ovarian cancer: Global expert consensus on testing and a comparison of companion diagnostics. Eur J Cancer 215, 115169.

Raimondo, T.M., Reed, K., Shi, D., Langer, R., and Anderson, D.G. (2023). Delivering the next generation of cancer immunotherapies with RNA. Cell 186, 1535–1540.

Rangarajan, A., and Weinberg, R.A. (2003). Opinion: Comparative biology of mouse versus human cells: modelling human cancer in mice. Nat Rev Cancer 3, 952–959.

Raudvere, U., Kolberg, L., Kuzmin, I., Arak, T., Adler, P., Peterson, H., and Vilo, J. (2019). g:Profiler: a web server for functional enrichment analysis and conversions of gene lists (2019 update). Nucleic Acids Res 47, W191–W198.

Robinson, M.D., and Oshlack, A. (2010). A scaling normalization method for differential expression analysis of RNA-seq data. Genome Biol 11, R25.

Rossi, M.J., DiDomenico, S.F., Patel, M., and Mazin, A.V. (2021). RAD52: Paradigm of Synthetic Lethality and New Developments. Front Genet 12, 780293.

Saelens, W., Cannoodt, R., and Saeys, Y. (2018). A comprehensive evaluation of module detection methods for gene expression data. Nat Commun 9, 1090.

Sansom, O.J., Meniel, V., Wilkins, J.A., Cole, A.M., Oien, K.A., Marsh, V., Jamieson, T.J., Guerra, C., Ashton, G.H., Barbacid, M., et al. (2006). Loss of Apc allows phenotypic manifestation of the transforming properties of an endogenous K-ras oncogene in vivo. Proc Natl Acad Sci U S A 103, 14122–14127.

Sarropoulos, I., Marin, R., Cardoso-Moreira, M., and Kaessmann, H. (2019). Developmental dynamics of lncRNAs across mammalian organs and species. Nature 571, 510–514.

Schmitt, A.M., and Chang, H.Y. (2016). Long Noncoding RNAs in Cancer Pathways. Cancer Cell 29, 452–463.

Seok, J., Warren, H.S., Cuenca, A.G., Mindrinos, M.N., Baker, H.V., Xu, W., Richards, D.R., McDonald-Smith, G.P., Gao, H., Hennessy, L., et al. (2013). Genomic responses in mouse models poorly mimic human inflammatory diseases. Proc Natl Acad Sci U S A 110, 3507–3512.

Shao, C., Indeglia, A., Foster, M., Casey, K., Leung, J., Modarai, S.R., Leu, J.I., Duong, B., Mes-Masson, A.M., Sims-Mourtada, J., et al. (2026). Mutant p53 binds and controls estrogen receptor activity to drive endocrine resistance in ovarian cancer. Genes Dev 40, 199–214.

Shih, A.H., Jiang, Y., Meydan, C., Shank, K., Pandey, S., Barreyro, L., Antony-Debre, I., Viale, A., Socci, N., Sun, Y., et al. (2015). Mutational cooperativity linked to combinatorial epigenetic gain of function in acute myeloid leukemia. Cancer Cell 27, 502–515.

Somerfield, P.J., Clarke, K.R., and Gorley, R.N. (2021). Analysis of similarities (ANOSIM) for 2-way layouts using a generalised ANOSIM statistic, with comparative notes on Permutational Multivariate Analysis of Variance (PERMANOVA). Austral Ecology.

Somervaille, T.C., Matheny, C.J., Spencer, G.J., Iwasaki, M., Rinn, J.L., Witten, D.M., Chang, H.Y., Shurtleff, S.A., Downing, J.R., and Cleary, M.L. (2009). Hierarchical maintenance of MLL myeloid leukemia stem cells employs a transcriptional program shared with embryonic rather than adult stem cells. Cell Stem Cell 4, 129–140.

Teschendorff, A.E., and Enver, T. (2017). Single-cell entropy for accurate estimation of differentiation potency from a cell’s transcriptome. Nat Commun 8, 15599.

Thorsson, V., Gibbs, D.L., Brown, S.D., Wolf, D., Bortone, D.S., Ou Yang, T.H., Porta-Pardo, E., Gao, G.F., Plaisier, C.L., Eddy, J.A., et al. (2019). The Immune Landscape of Cancer. Immunity 51, 411–412.

Ulitsky, I. (2016). Evolution to the rescue: using comparative genomics to understand long non-coding RNAs. Nat Rev Genet 17, 601–614.

van den Berg, R.A., Hoefsloot, H.C., Westerhuis, J.A., Smilde, A.K., and van der Werf, M.J. (2006). Centering, scaling, and transformations: improving the biological information content of metabolomics data. BMC Genomics 7, 142.

Vergote, I.B., Jimeno, A., Joly, F., Katsaros, D., Coens, C., Despierre, E., Marth, C., Hall, M., Steer, C.B., Colombo, N., et al. (2014). Randomized phase III study of erlotinib versus observation in patients with no evidence of disease progression after first-line platin-based chemotherapy for ovarian carcinoma: a European Organisation for Research and Treatment of Cancer-Gynaecological Cancer Group, and Gynecologic Cancer Intergroup study. J Clin Oncol 32, 320–326.

Vilella, A.J., Severin, J., Ureta-Vidal, A., Heng, L., Durbin, R., and Birney, E. (2009). EnsemblCompara GeneTrees: Complete, duplication-aware phylogenetic trees in vertebrates. Genome Res 19, 327–335.

Wainwright, E.N., and Scaffidi, P. (2017). Epigenetics and Cancer Stem Cells: Unleashing, Hijacking, and Restricting Cellular Plasticity. Trends Cancer 3, 372–386.

Wen, Y., Wu, Y., Xu, B., Lin, J., and Zhu, H. (2022). Fasim-LongTarget enables fast and accurate genome-wide lncRNA/DNA binding prediction. Comput Struct Biotechnol J 20, 3347–3350.

Winkle, M., El-Daly, S.M., Fabbri, M., and Calin, G.A. (2021). Noncoding RNA therapeutics - challenges and potential solutions. Nat Rev Drug Discov 20, 629–651.

Wu, Y., Wu, B.Z., Ellenbogen, Y., Kant, J.B.Y., Yu, P., Li, X., Caloren, L., Sotov, V., Tran, C., Restrepo, M., et al. (2025). Neurodevelopmental hijacking of oligodendrocyte lineage programs drives glioblastoma infiltration. Dev Cell 60, 2420–2433 e2412.

Yap, K.L., Li, S., Munoz-Cabello, A.M., Raguz, S., Zeng, L., Mujtaba, S., Gil, J., Walsh, M.J., and Zhou, M.M. (2010). Molecular interplay of the noncoding RNA ANRIL and methylated histone H3 lysine 27 by polycomb CBX7 in transcriptional silencing of INK4a. Mol Cell 38, 662–674.

Yin, M., Li, X., Tan, S., Zhou, H.J., Ji, W., Bellone, S., Xu, X., Zhang, H., Santin, A.D., Lou, G., et al. (2016). Tumor-associated macrophages drive spheroid formation during early transcoelomic metastasis of ovarian cancer. J Clin Invest 126, 4157–4173.

Yue, F., Cheng, Y., Breschi, A., Vierstra, J., Wu, W., Ryba, T., Sandstrom, R., Ma, Z., Davis, C., Pope, B.D., et al. (2014). A comparative encyclopedia of DNA elements in the mouse genome. Nature 515, 355–364.

Zhang, X., Lan, Y., Xu, J., Quan, F., Zhao, E., Deng, C., Luo, T., Xu, L., Liao, G., Yan, M., et al. (2019). CellMarker: a manually curated resource of cell markers in human and mouse. Nucleic Acids Res 47, D721–D728.

Zhao, Y., Li, J., Ting, K.K., Chen, J., Coleman, P., Liu, K., Wan, L., Moller, T., Vadas, M.A., and Gamble, J.R. (2021). The VE-Cadherin/beta-catenin signalling axis regulates immune cell infiltration into tumours. Cancer Lett 496, 1–15.

