## Supplementary Figures for "Lineage-specific lncRNAs critically determine cross-species differences in tumors"

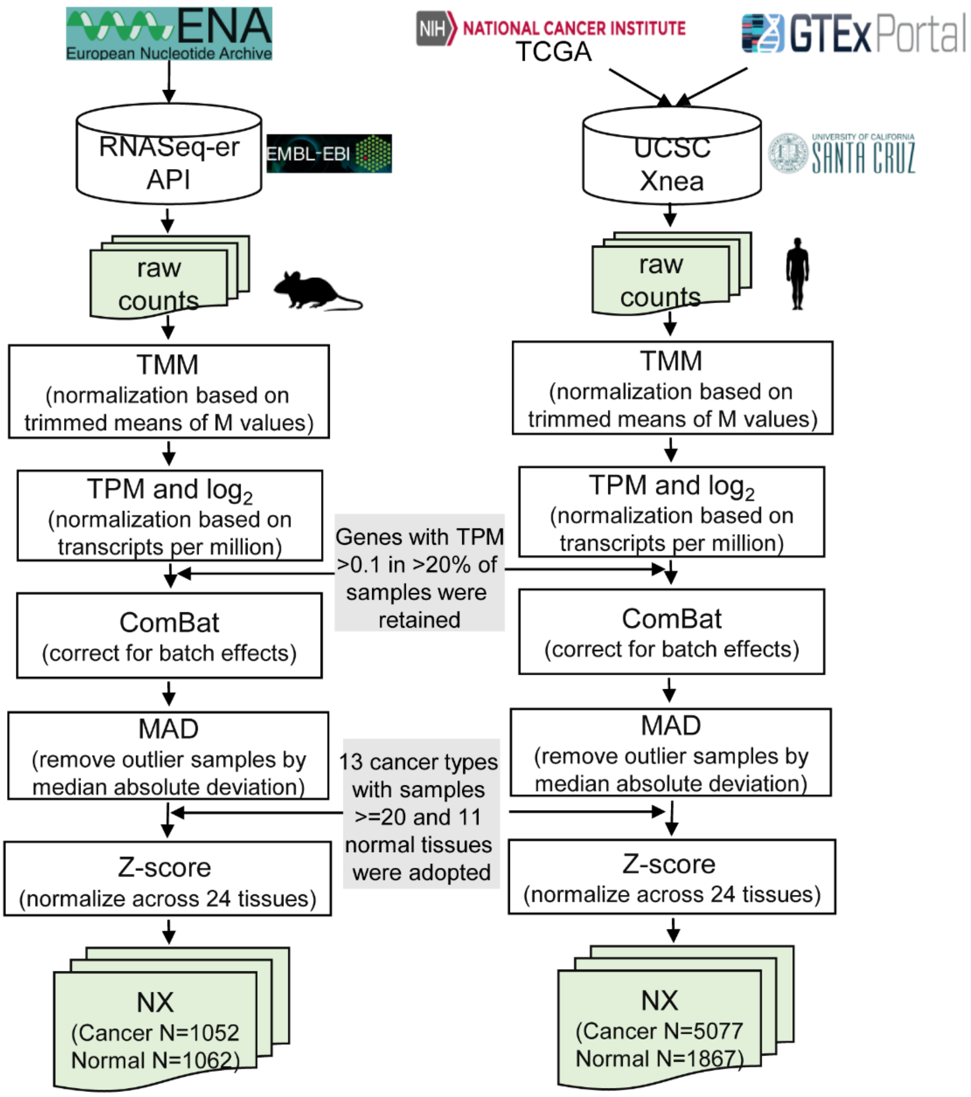


Supplementary Figure 1-1. Schematic overview of data normalization and integration. Raw count data were normalized using the trimmed mean of M values (TMM) and transcripts per million (TPM) methods, and batch correction was performed using the *ComBat* algorithm. Outlier samples were removed using a median absolute deviation (MAD)-based approach. Finally, TPM values were rescaled to z-scores across all cancer types and normal tissues to generate normalized expression (NX) values.

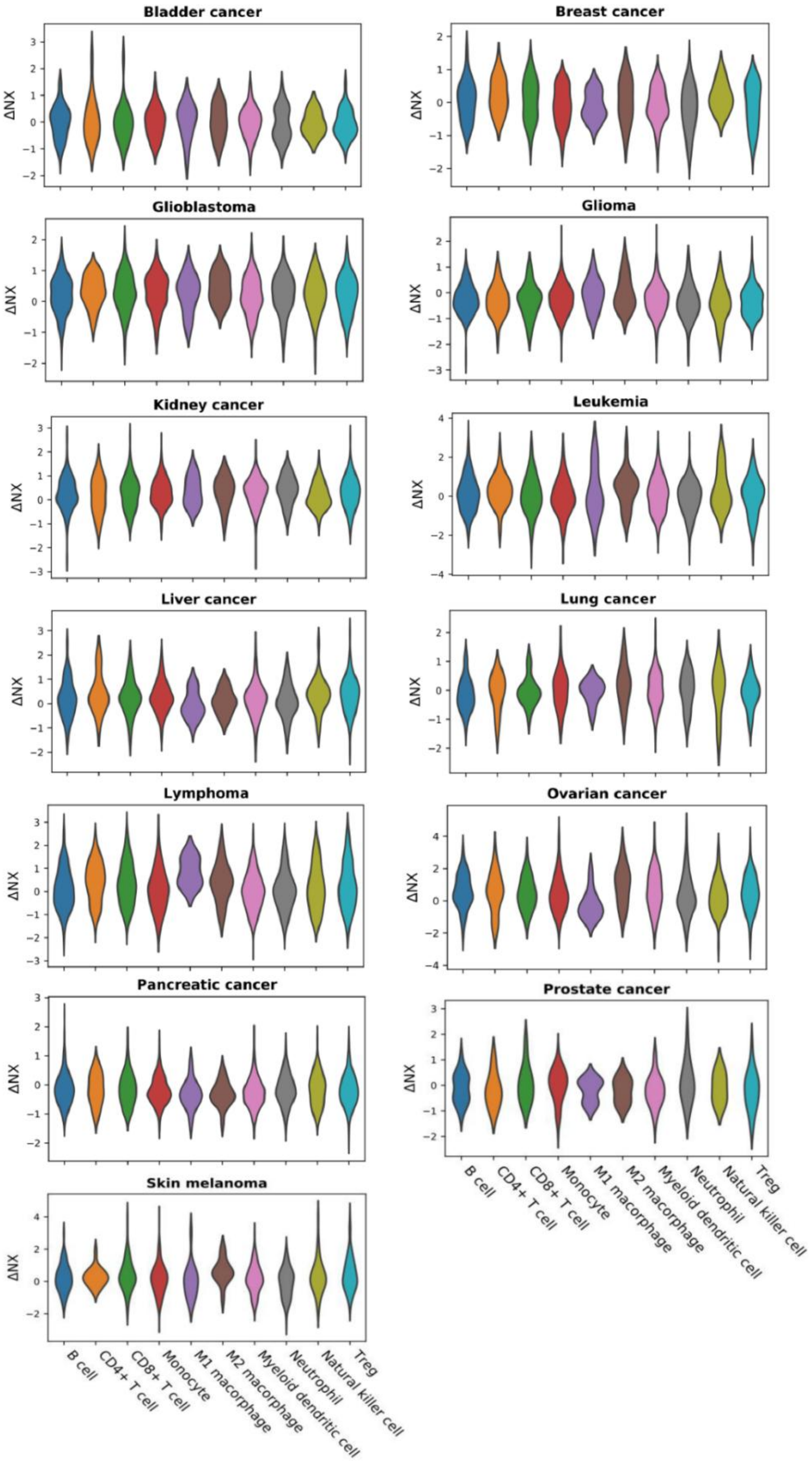


Supplementary Figure 1-2. The cross-species ΔNX values of marker genes for ten cancer-infiltrating immune cell types display approximately zero-centered distributions across tumor types, suggesting no systematic species bias in immune marker expression.

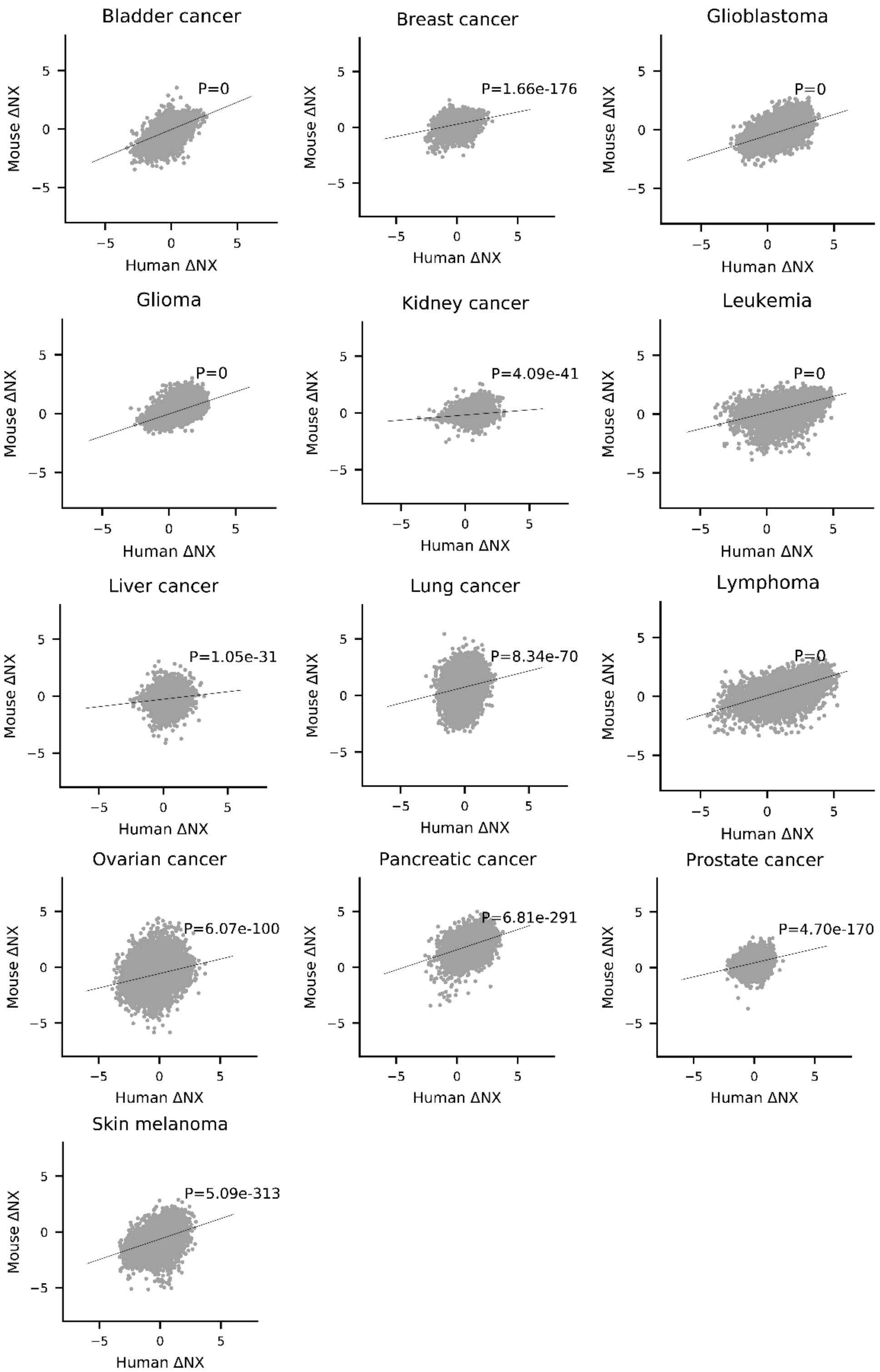


Supplementary Figure 1-3. Differential expression (ΔNX) of all one-to-one orthologous protein-coding genes (1:1 PCGs) across the 13 tumors shows significant cross-species Pearson correlations, indicating partial conservation of gene-level transcriptional changes.

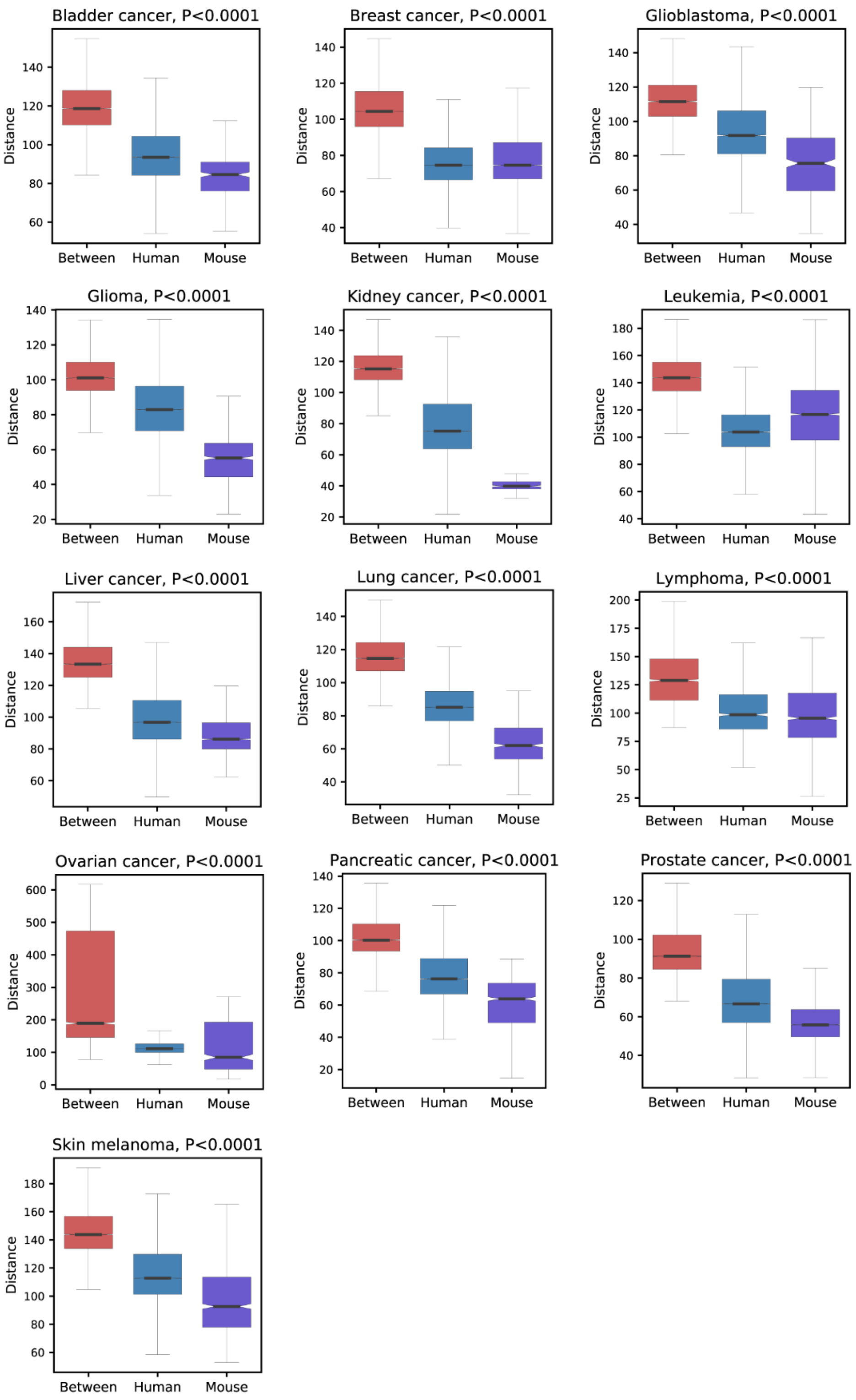


Supplementary Figure 1-4. ANOSIM analysis demonstrates that transcriptomic variation between species exceeds variation within species across tumor types (10,000 permutations; $P < 10^{-4}$ in all cancers). “Between” indicates cross-species comparisons; “Human” and “Mouse” indicate within-species comparisons.

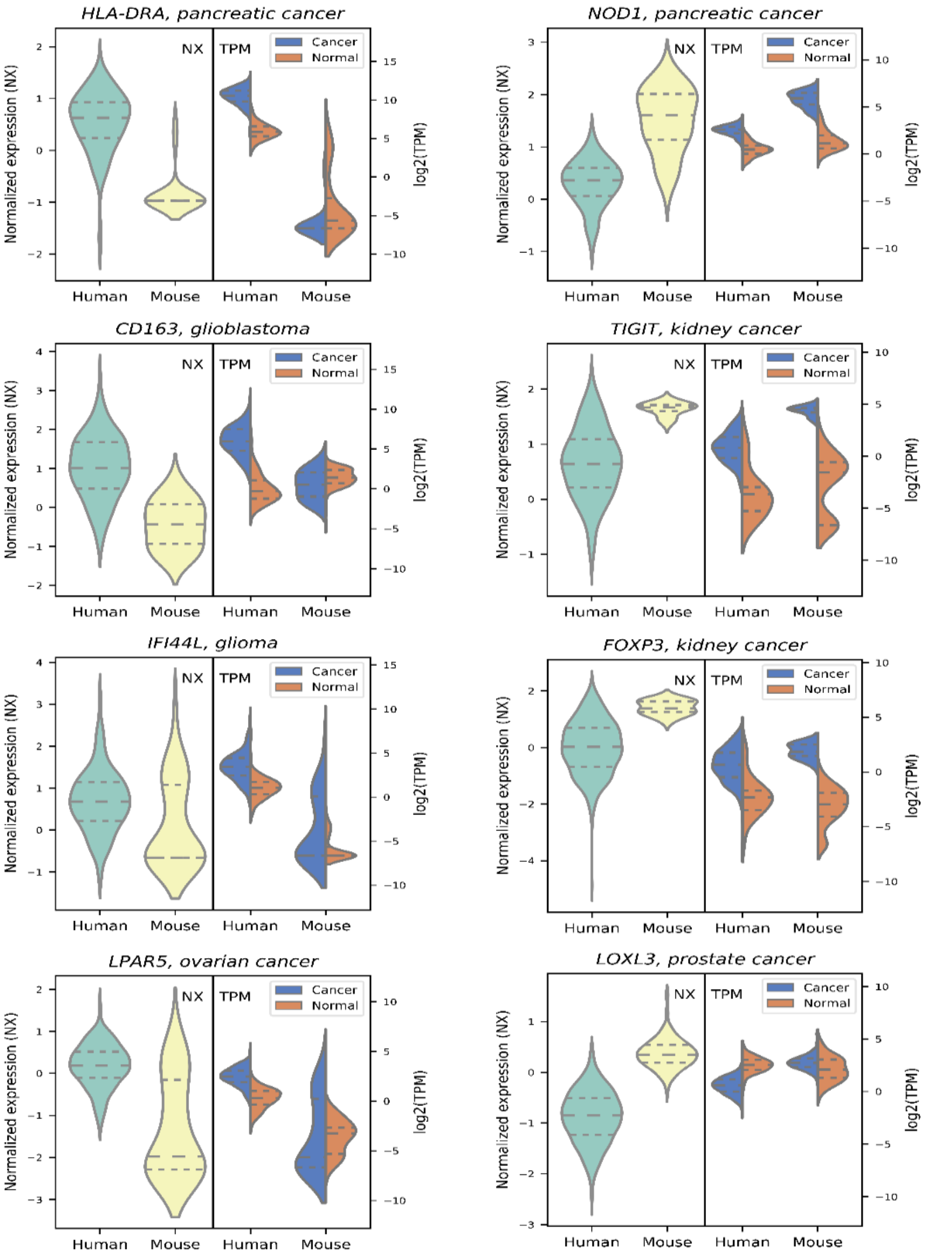


Supplementary Figure 1-5. Representative cancer-related genes showing discordance between intra-species differential expression and cross-species transcriptional divergence. Left panels show genes with higher expression in human tumors; right panels show genes with higher expression in mouse tumors.

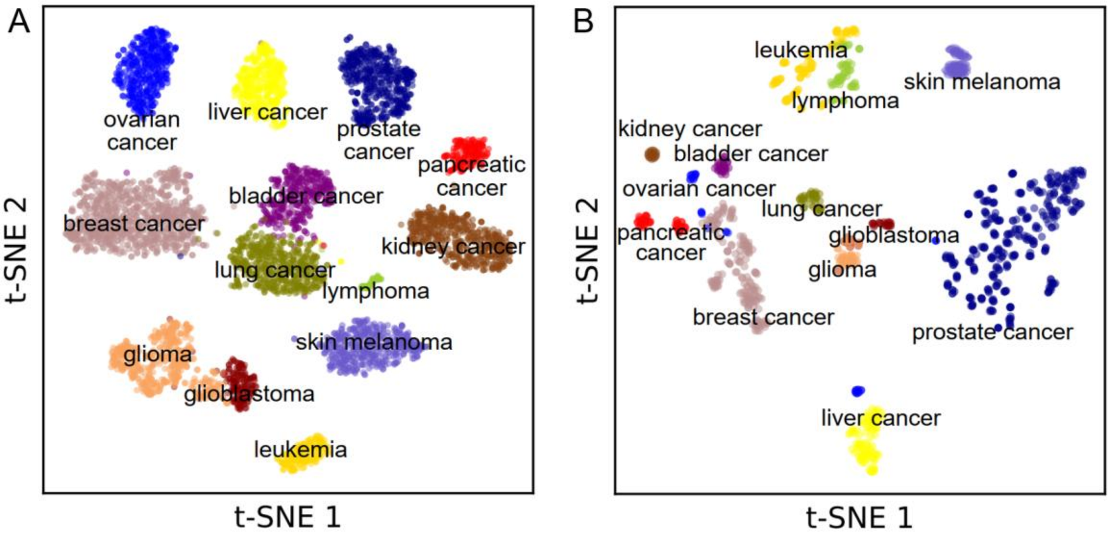


Supplementary Figure 1-6. Characterization of tumor identity by LS lncRNAs. (A) t-SNE embedding based on PS/SS lncRNA expression separates human tumor types. (B) t-SNE embedding based on RS lncRNA expression separates mouse tumor types.

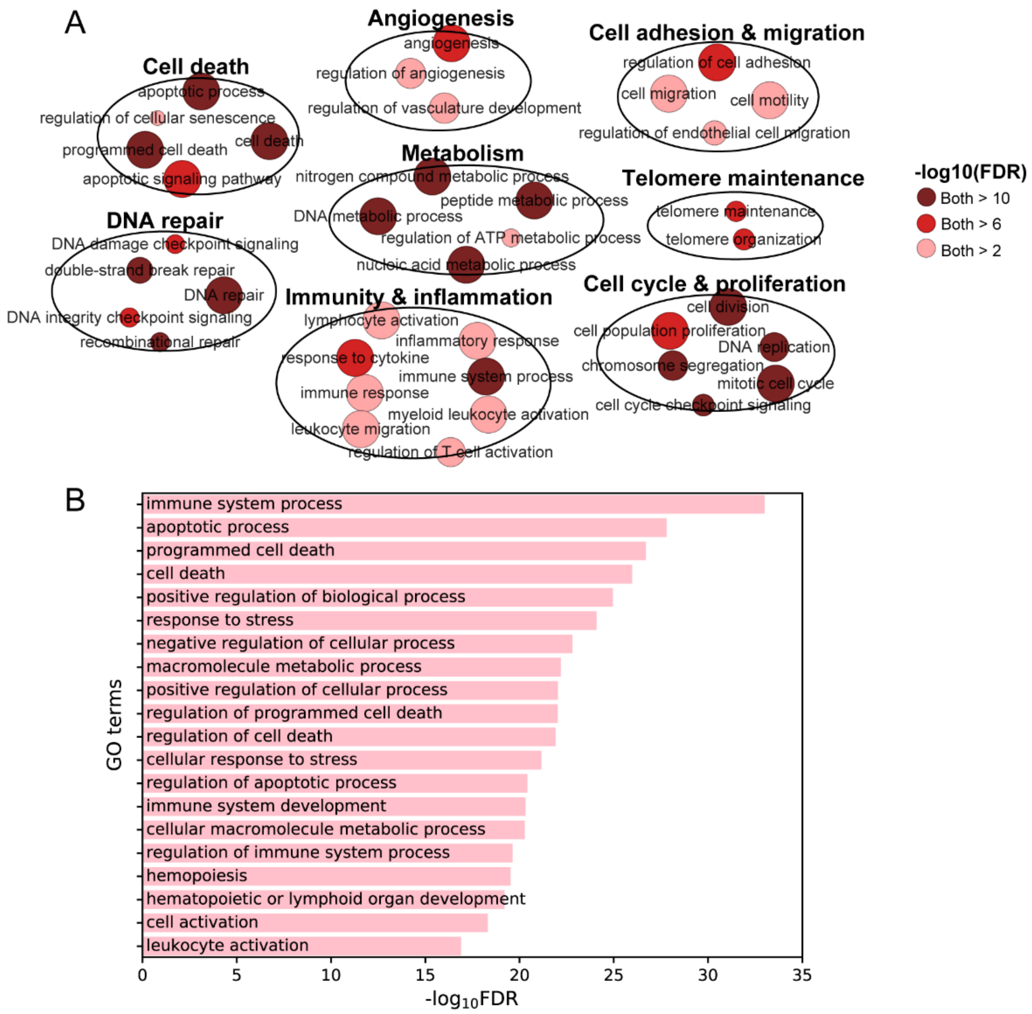


Supplementary Figure 1-7. TDGs are preferentially associated with immune-related functions. (A) Shared DEGs across multiple tumors are enriched in cell cycle, DNA repair, adhesion, and migration pathways, but show limited enrichment in immune-related GO terms. (B) In contrast, TDGs identified in ≥2 tumors are strongly enriched in immune-related GO terms (top 20 terms shown).

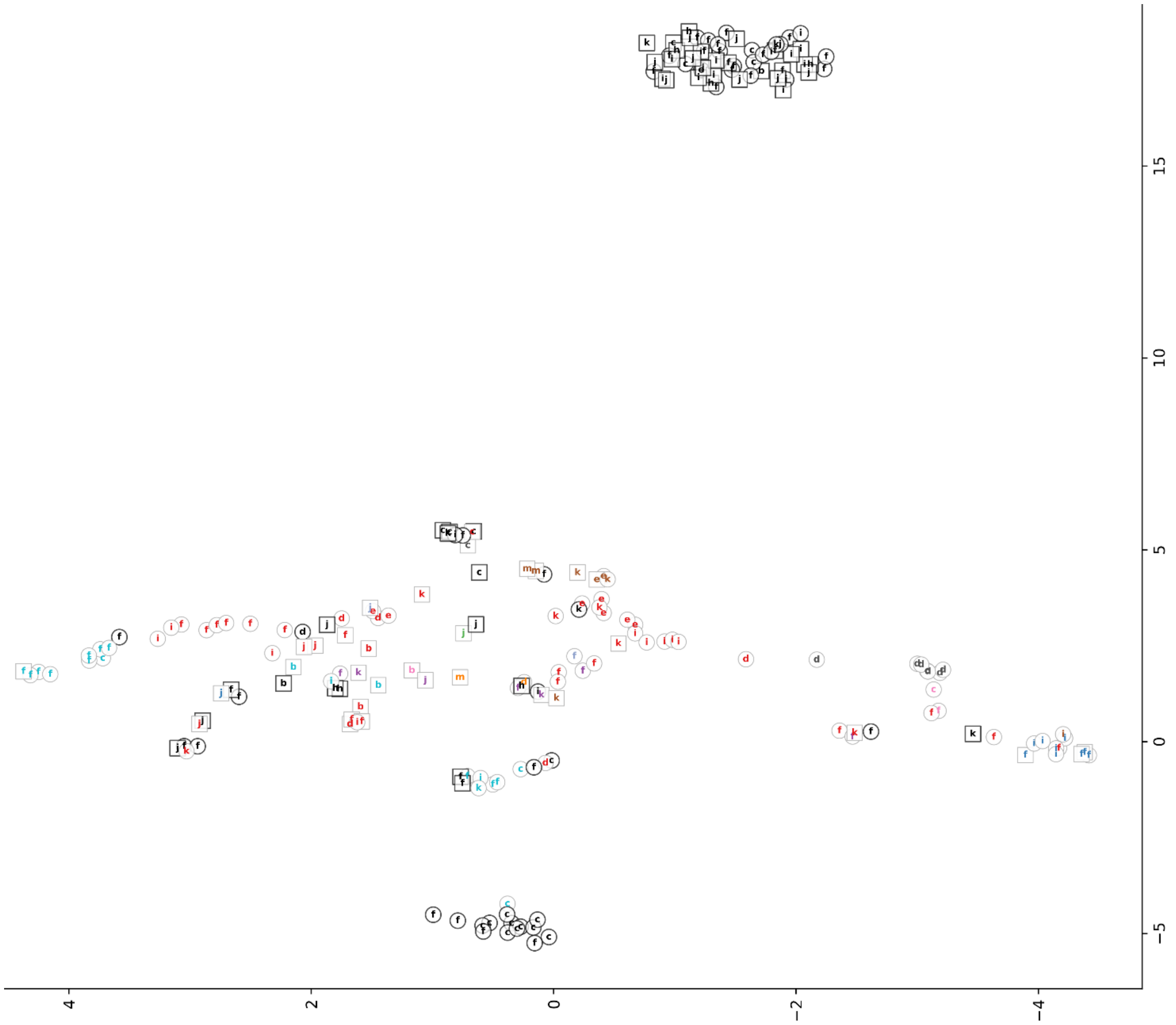


Supplementary Figure 2-1. UMAP projection of all modules based on hallmarks under the Preserved condition, highlighting inter-tumor variability in retained regulatory architecture.

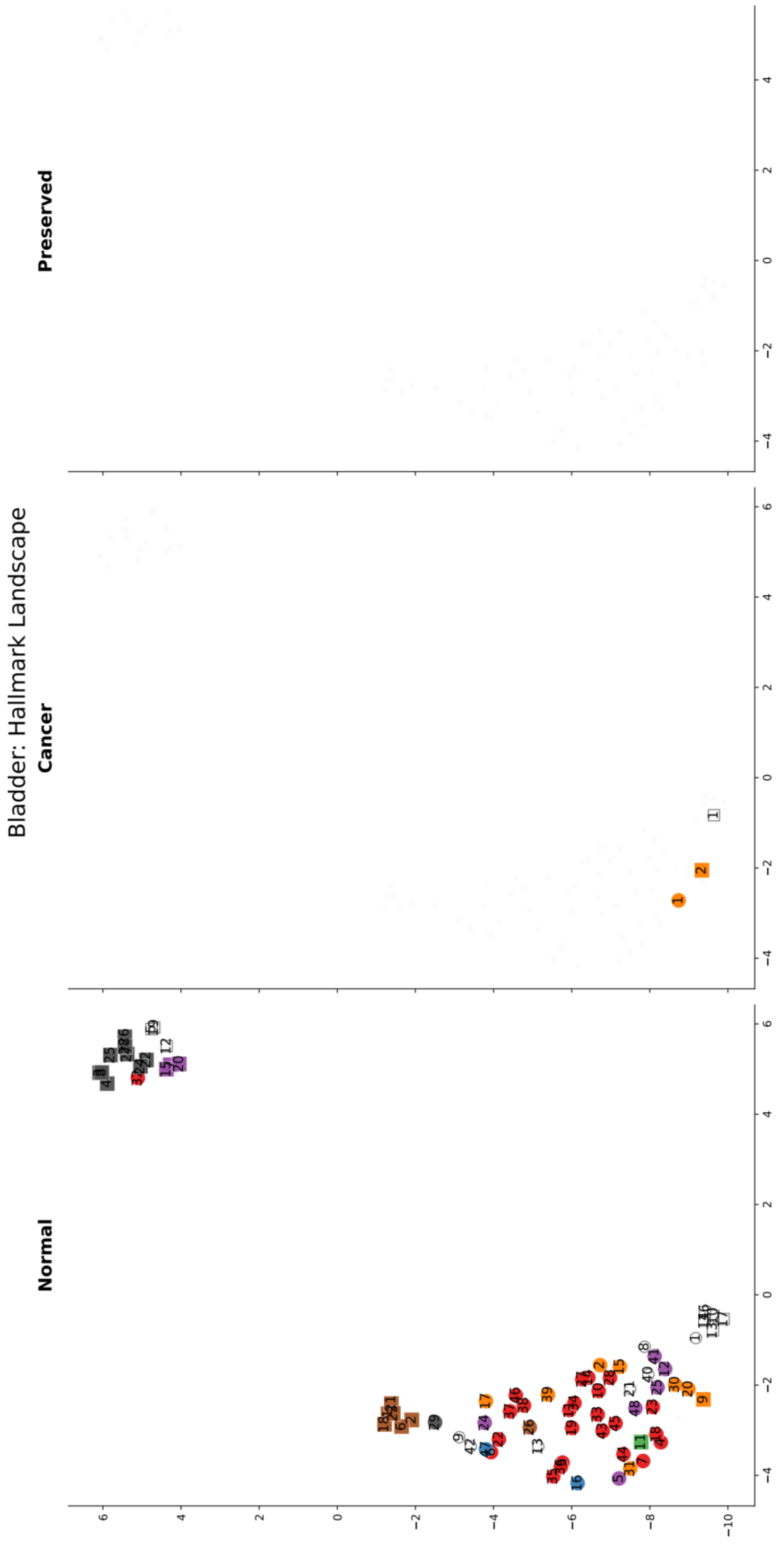


Supplementary Figure 3-1. UMAP projection of bladder cancer modules based on hallmarks. Human Normal modules form a large cluster, suggesting greater baseline regulatory complexity compared with mouse models. A reduction in module diversity from Normal to Cancer is observed. Few modules persist in the Preserved condition, consistent with extensive regulatory remodeling reported in urothelial carcinoma (Robertson et al., 2017).

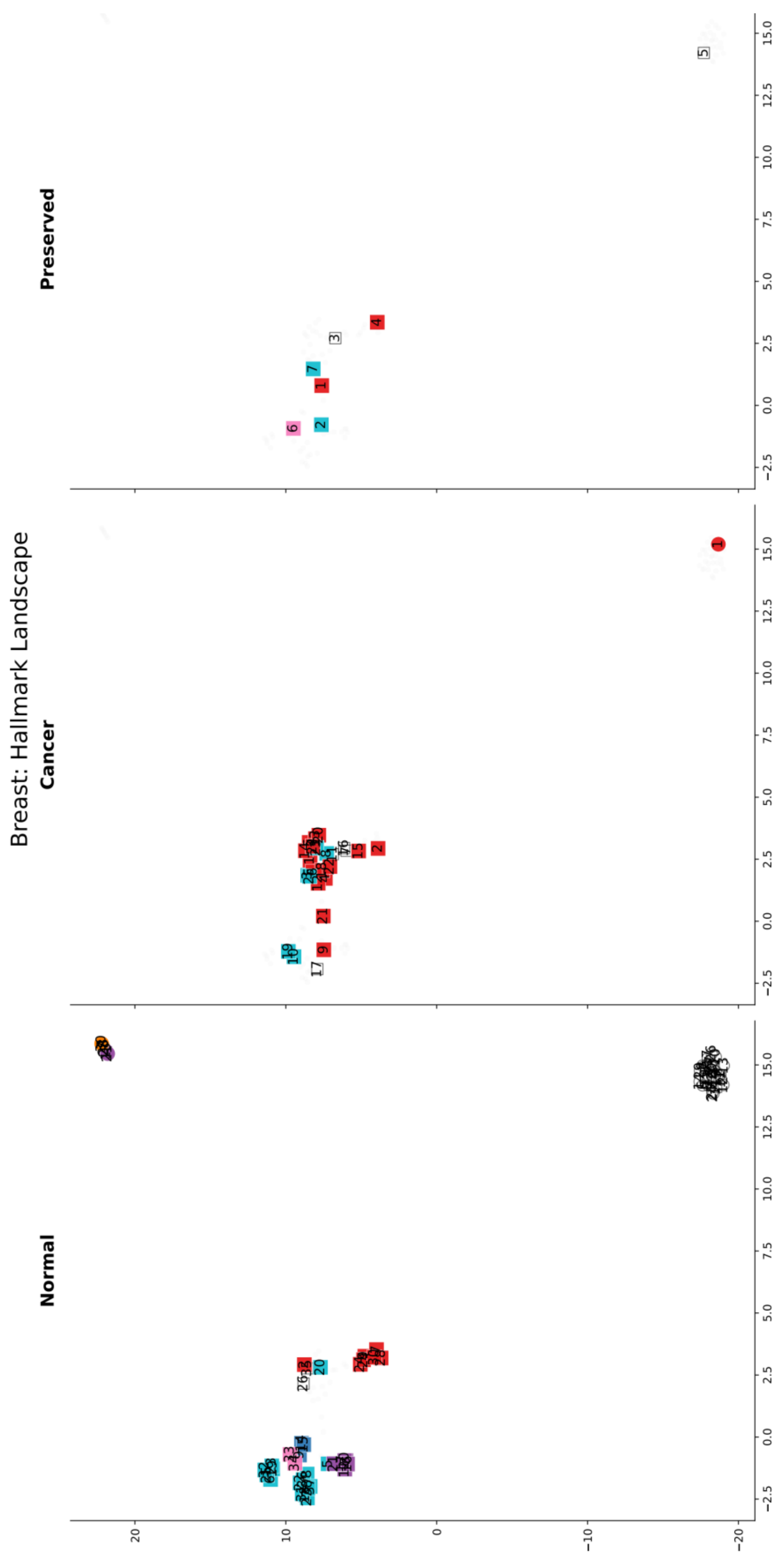


Supplementary Figure 3-2. UMAP projection of breast cancer modules based on hallmarks. Normal human breast tissue exhibits complex regulatory organization, consistent with cyclical hormone-driven proliferation regulated by PS lncRNAs (Lin et al., 2016). Multiple *Unclassified* human modules (forming a tight cluster) in the Normal condition are enriched for WikiPathways. GEMM mouse models are driven by early oncogene activation in the mammary tissue from birth (Pfefferle et al., 2013); this may explain the presence of multiple mouse modules in the Cancer and Preserved conditions, as the "normal" (pre-tumor) tissue is already transcriptionally hijacked and "hard-wired" for proliferation and epigenetic remodeling (Herschkowitz et al., 2007).

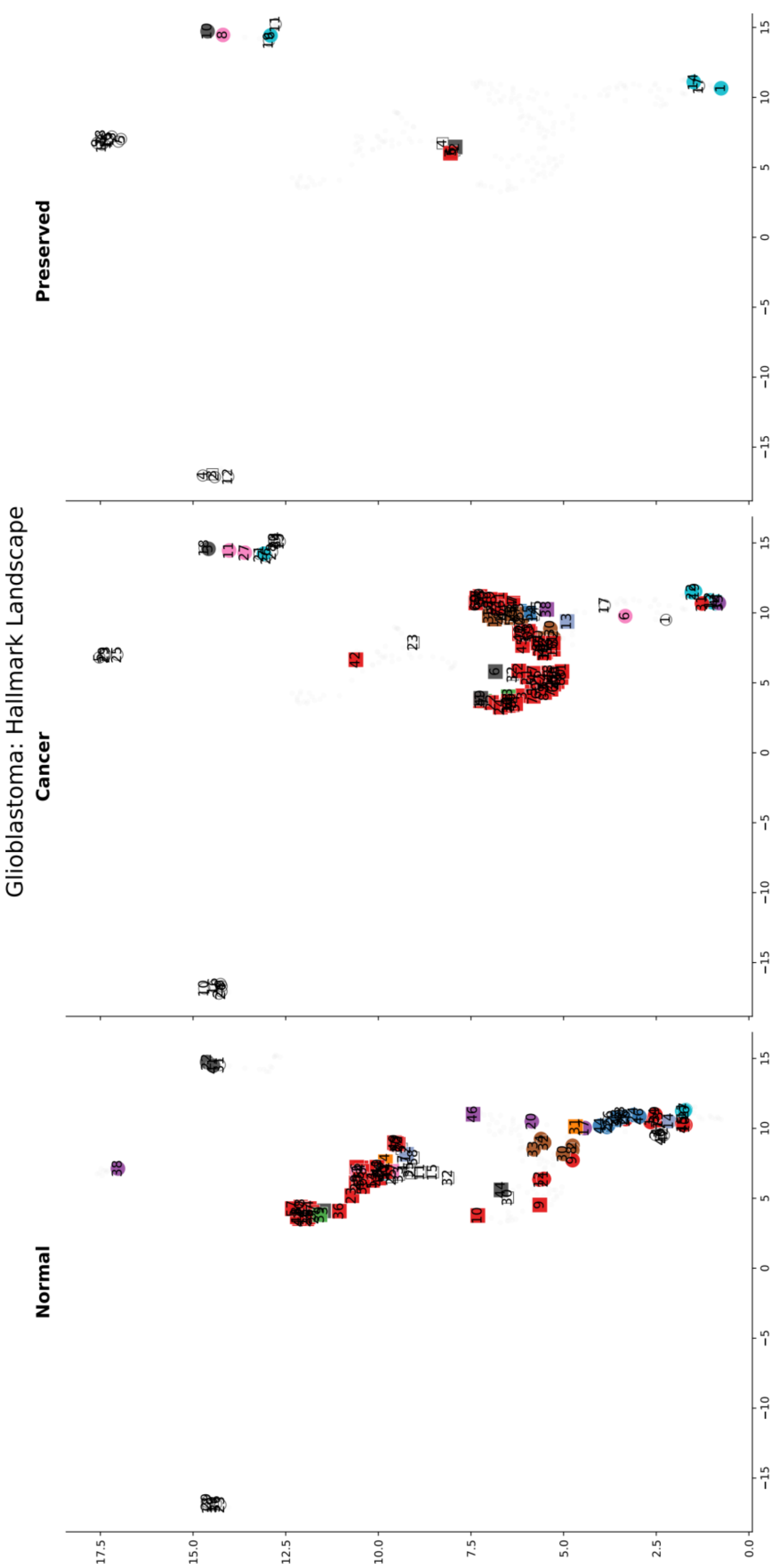


Supplementary Figure 3-3. UMAP projection of glioblastoma modules based on hallmarks. The extensive module retention in Cancer is consistent with high cellular plasticity and heterogeneity. Mouse Cancer modules are predominantly associated with proliferative signaling and inflammation, reflecting the highly proliferative and microglia-infiltrated nature of the standard mouse model (Miyai et al., 2017), whereas human modules are characterized by non-mutational epigenetic reprogramming and growth suppressor inactivation. Human modules in the Preserved condition suggest continued reliance on neurodevelopmental regulatory programs (Brennan et al., 2013).

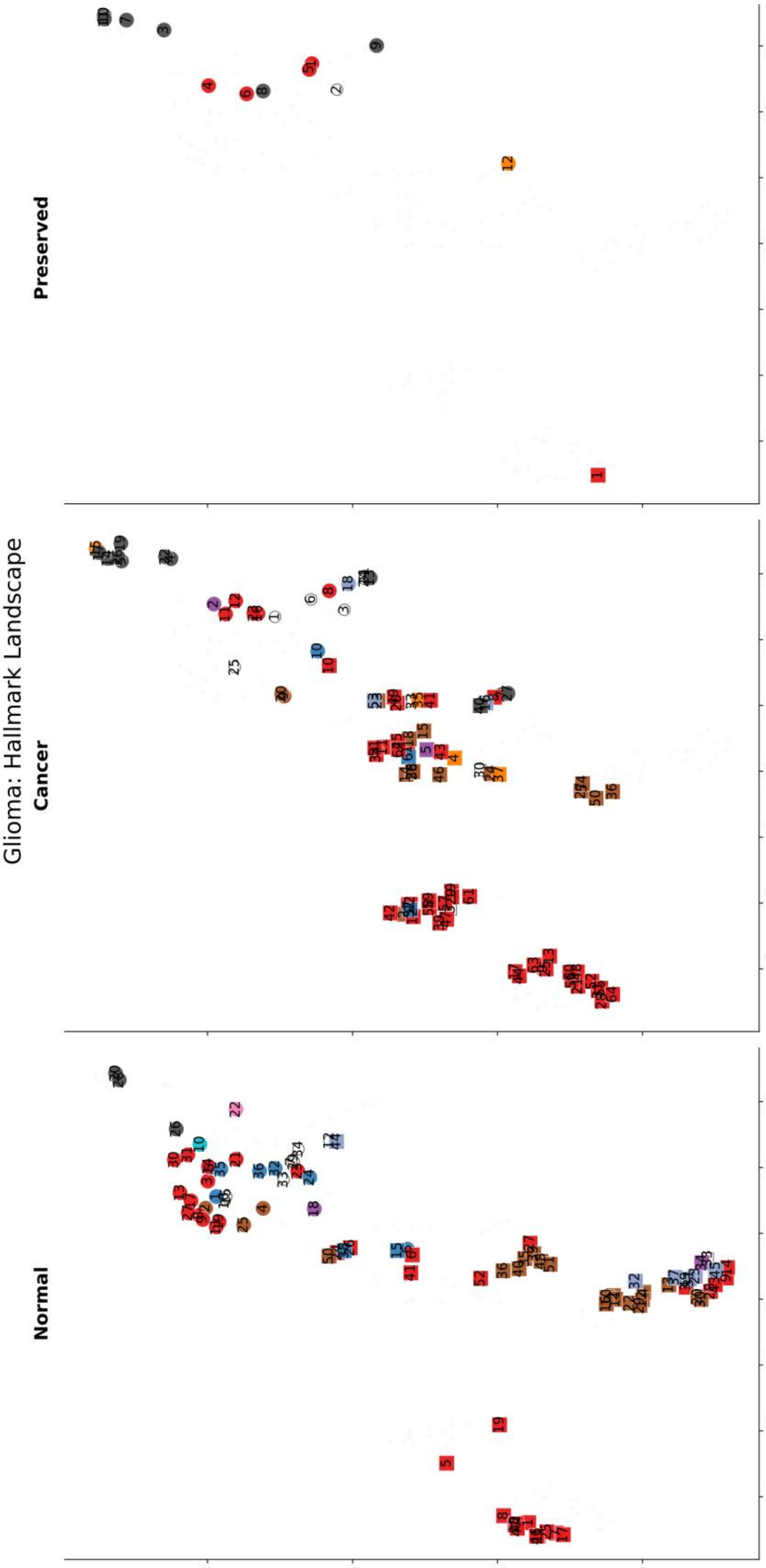


Supplementary Figure 3-4. UMAP projection of glioma modules based on hallmarks. Human glioma is enriched for *Unlocking phenotypic plasticity* modules across conditions, consistent with the reports that human glioma survives by locking or co-opting the normal neurodevelopmental plasticity pathways (Hanahan, 2022; Venteicher et al., 2017). In contrast, mouse glioma has only one proliferative module in the Preserved condition, potentially reflecting that mouse models usually rely on slamming the cell with overwhelming oncogenes, which instantly triggers massive hyper-proliferation and microglial infiltration. This topological divergence thus helps explain why standard mouse models fail to recapitulate the subtle lineage-plasticity mechanisms central to human glioma progression (Chen et al., 2012).

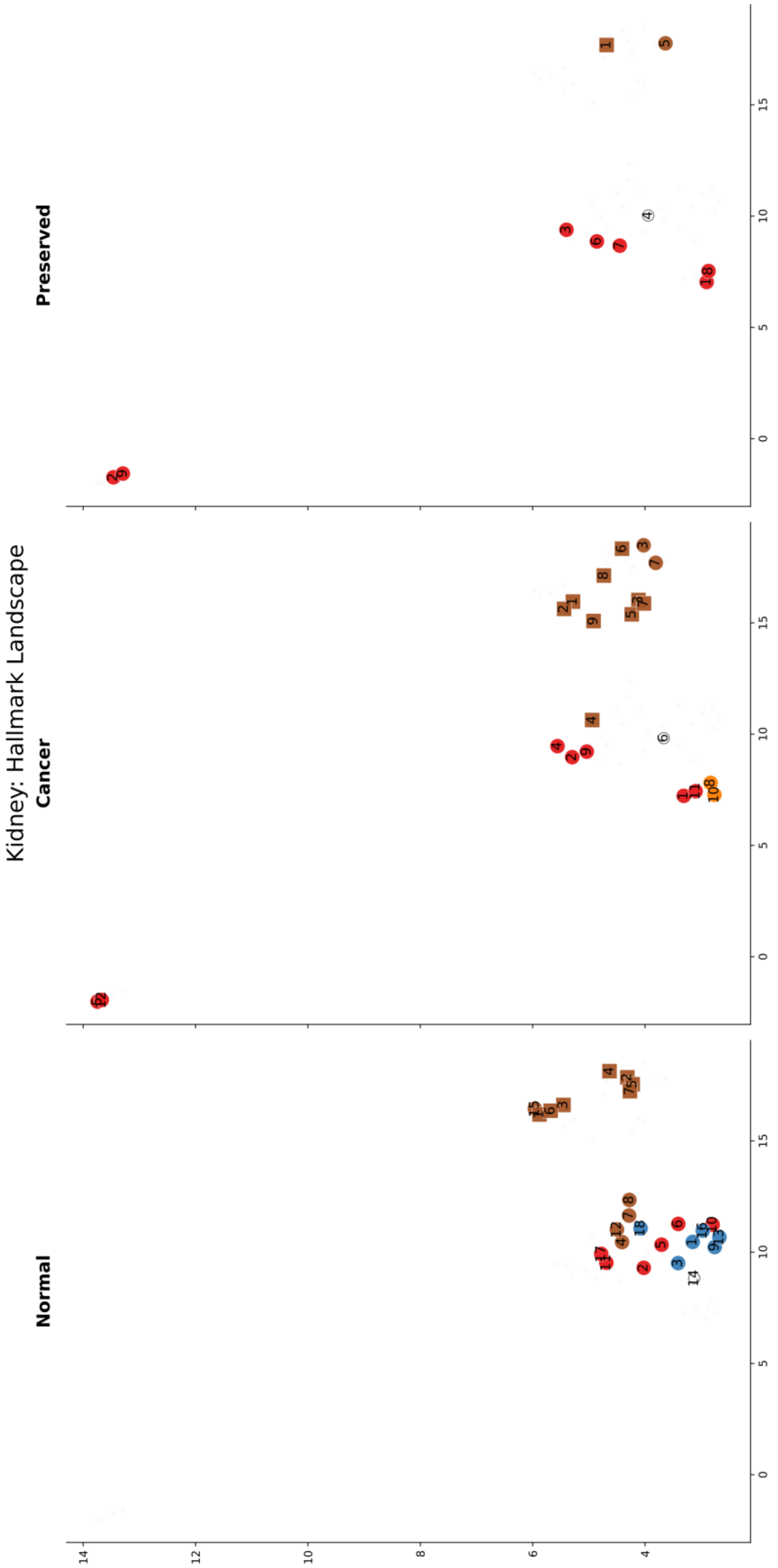


Supplementary Figure 3-5. UMAP projection of kidney cancer modules based on hallmarks (Figure 3B).

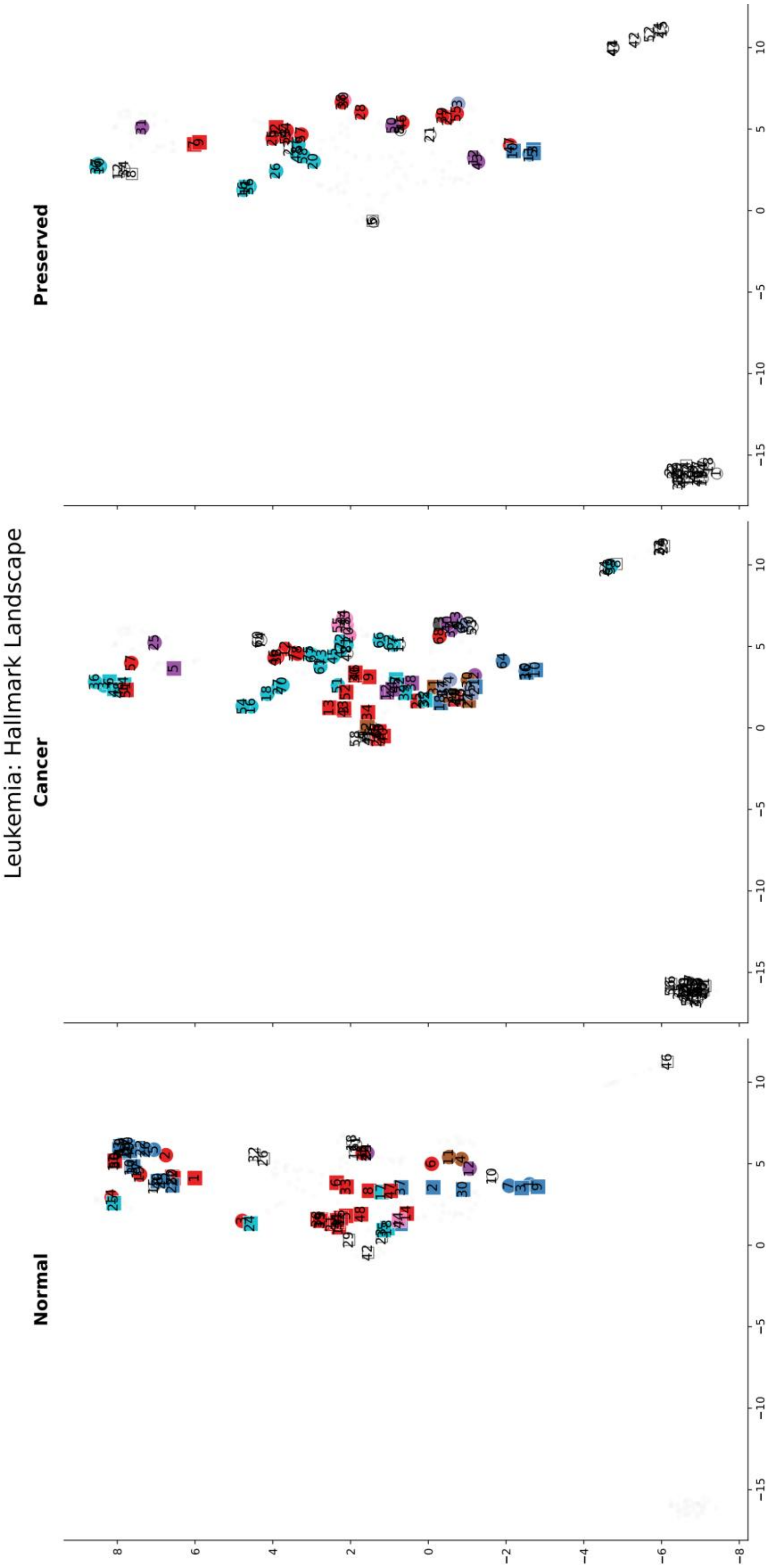


Supplementary Figure 3-6. UMAP projection of leukemia modules based on hallmarks(Figure 3A).

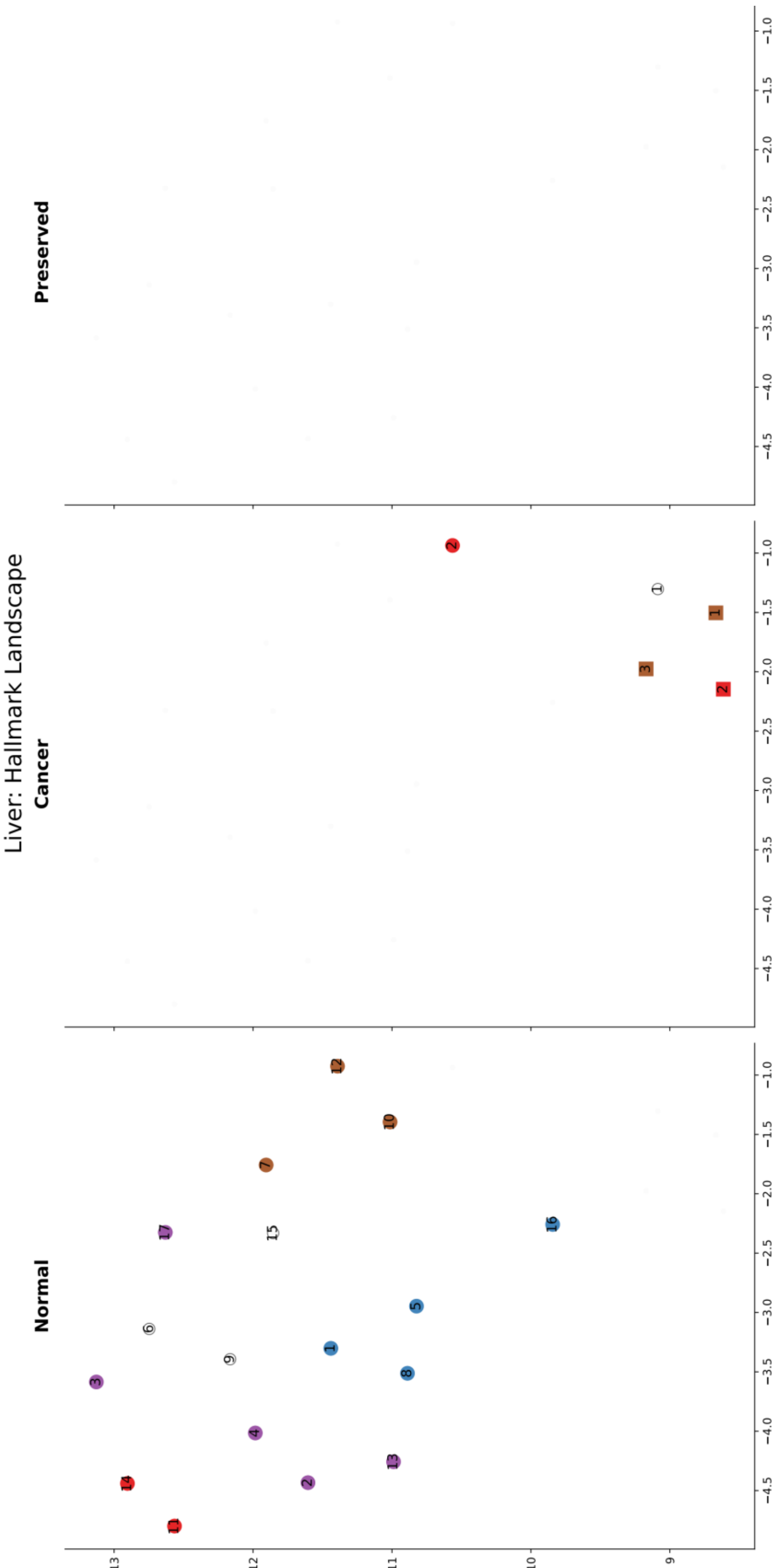


Supplementary Figure 3-7. UMAP projection of liver cancer modules based on hallmarks. In clinical liver cancer datasets, control tissues often exhibit chronic inflammatory and cirrhotic changes (Affo et al., 2017), which result in high transcriptional variability (Gao et al., 2019). The liver of laboratory mice living in sterile environments is transcriptionally comparatively homogeneous, and the limited variance in gene expression across control samples makes it hard to reach the threshold for correlation (r) between DEGs (>0.6). These differences may cause greater module diversity in the Normal condition. Indeed, ~2,200 DEGs were identified in humans, but only ~1,070 in mice. Also, chemically induced mouse models rely on massive acute inflammation, which differs mechanistically from the slow, spontaneous human disease and may explain the *Tumor-promoting inflammation* modules in the mouse Cancer condition.

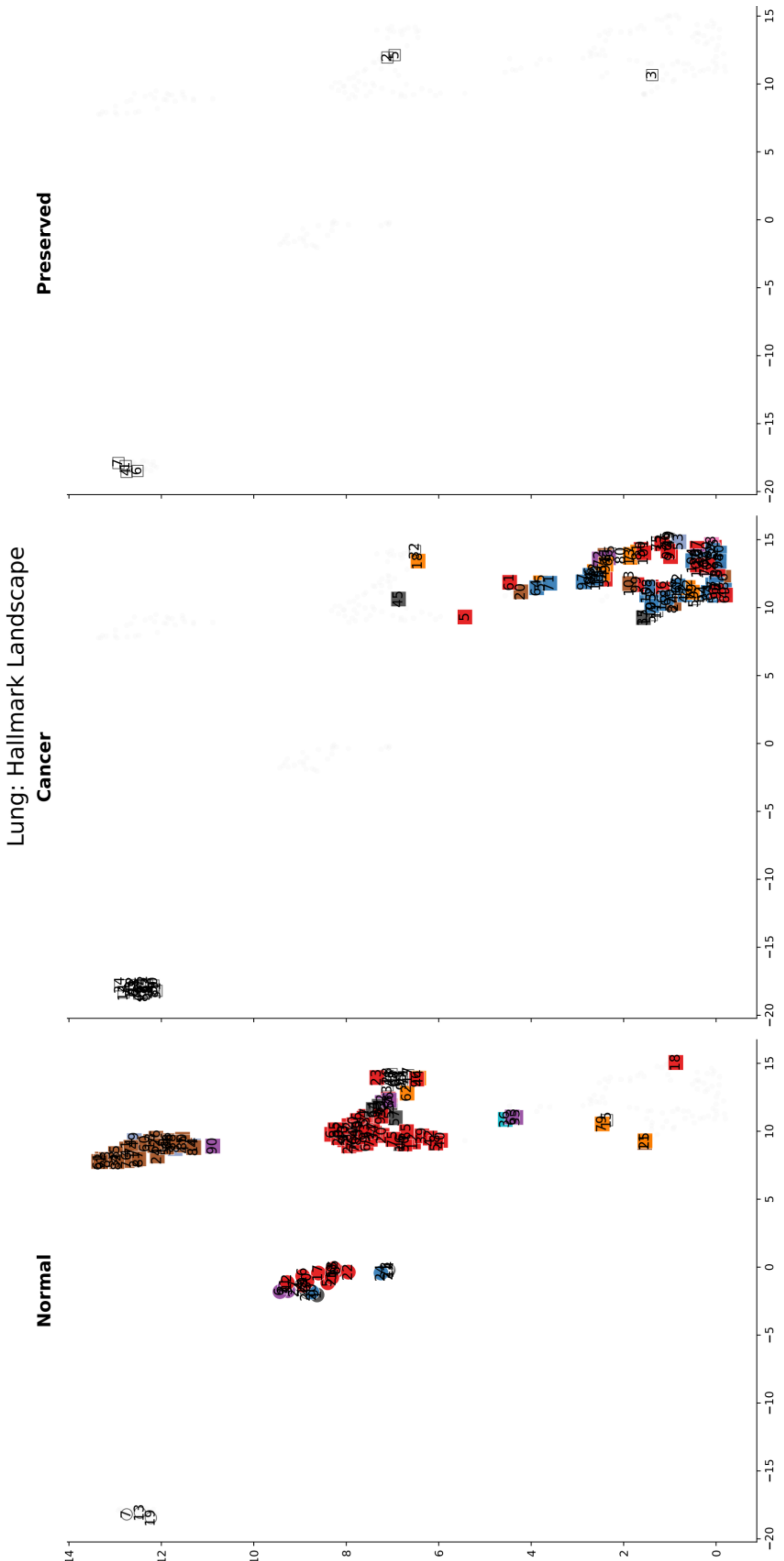


Supplementary Figure 3-8. UMAP projection of lung cancer modules based on hallmarks. Cross-species differences in lung cancer. In the Normal condition, one mouse cluster comprises dense *Tumor-promoting inflammation* modules, indicating massive, acute inflammatory responses. In the Cancer condition, there are many mouse but few human modules. The decrease in human modules from Normal to Cancer and Preserved reveals substantial cross-species differences, reflecting the key feature of human lung cancer, which is driven by decades of environmental assault (smoking, pollution) and the massive mutational burden shatters the genome (Alexandrov et al., 2013).

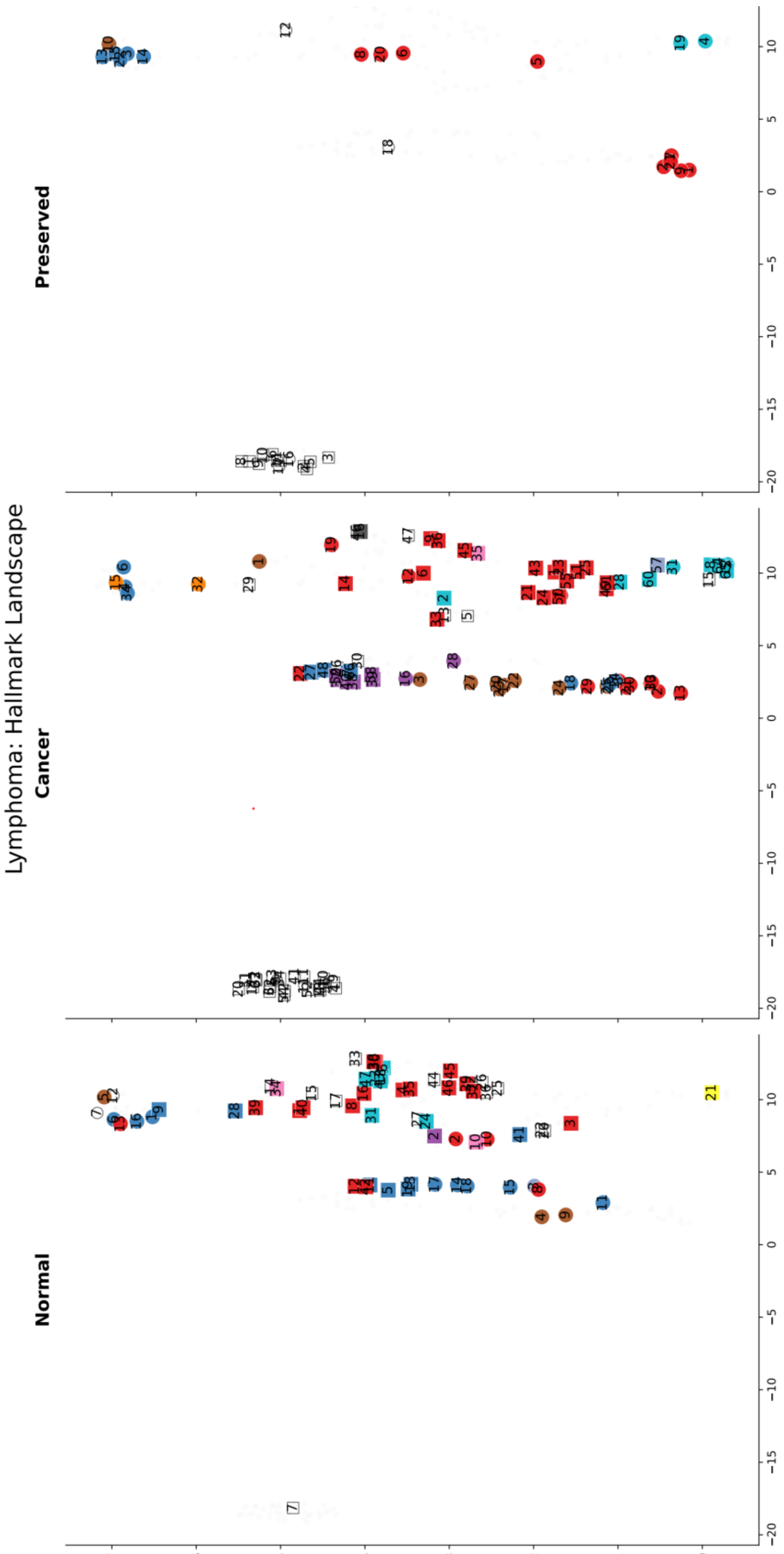


Supplementary Figure 3-9. UMAP projection of lymphoma modules based on hallmarks. Modules in the Normal and Cancer conditions are comparable to those of leukemia. However, in the Preserved condition, human lymphoma is characterized by *Sustaining proliferating signaling*, *Resisting programmed cell death*, and *Non-mutational epigenetic reprogramming* modules, but mouse models are characterized by exclusively *Unclassified* modules that do not even have any enriched WikiPathways. This may reflect the observation that the human lymphomas rely heavily on hijacking normal lymphocyte activation signals and coupling them with unrestrained proliferation (Basso and Dalla-Favera, 2015); in contrast, many mouse lymphomas are driven by endogenous mouse leukemia viruses (MuLV).

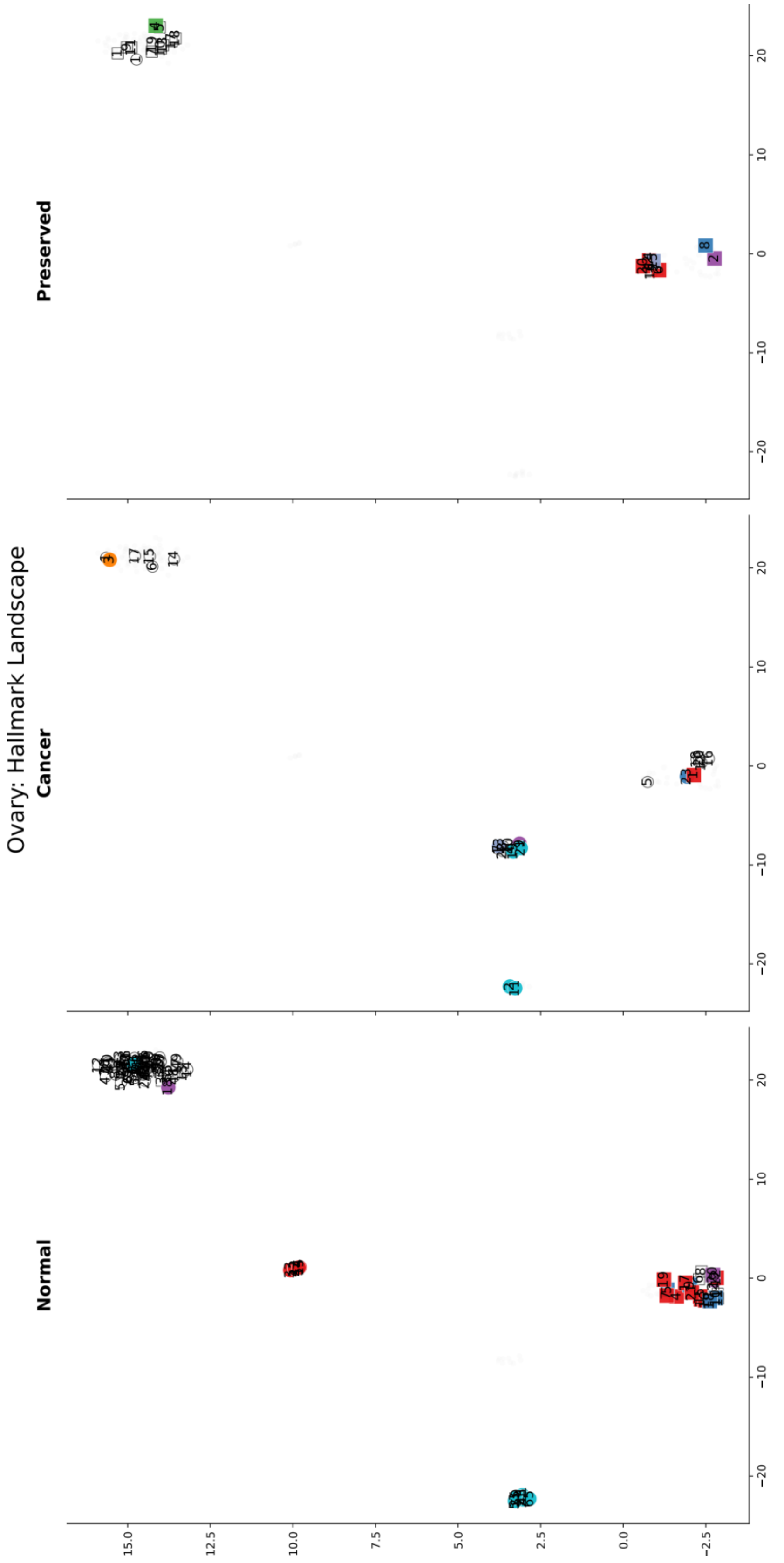


Supplementary Figure 3-10. UMAP projection of ovarian cancer modules based on hallmarks (Figure 3C).

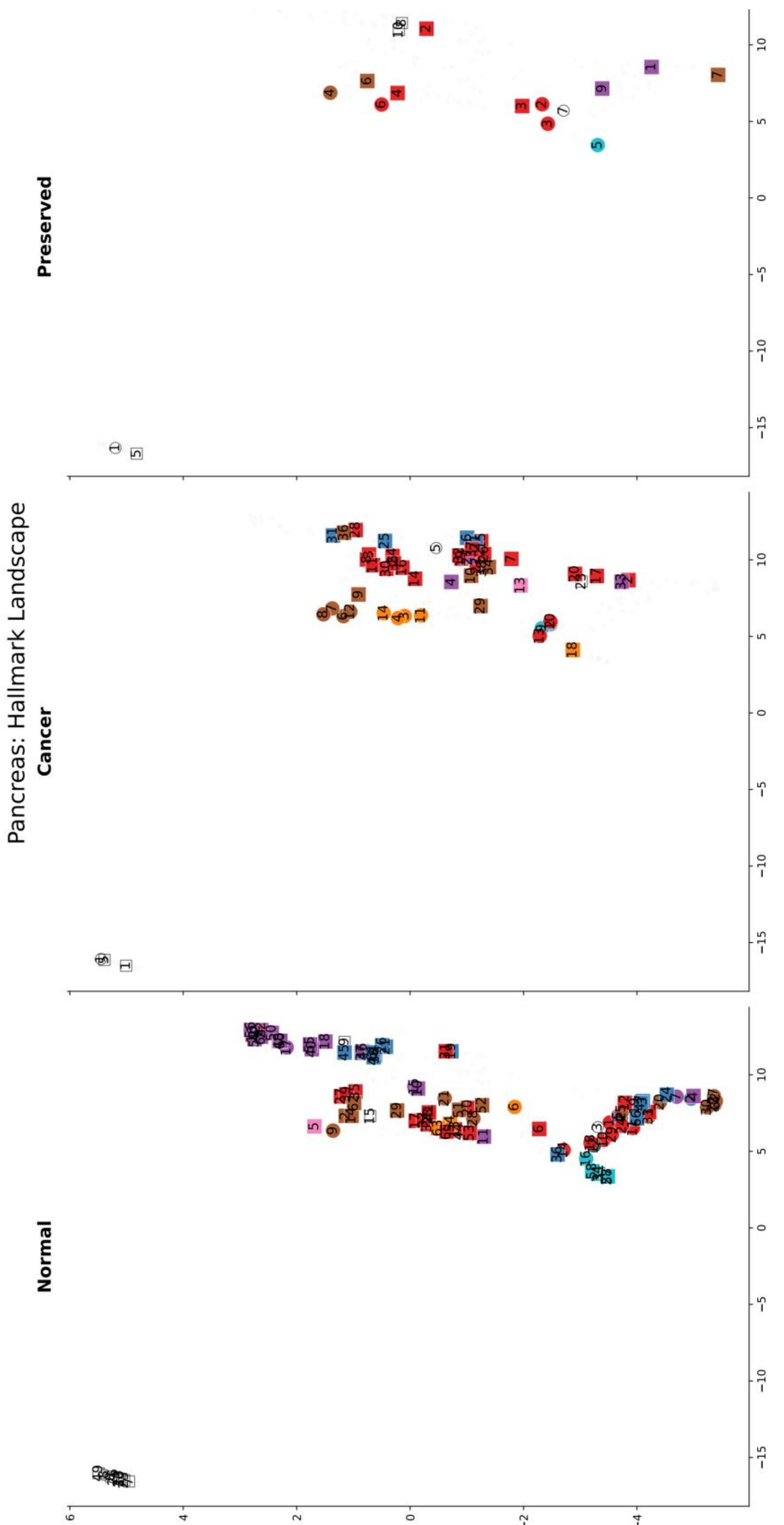


Supplementary Figure 3-11.UMAP projection of pancreatic cancer modules based on hallmarks. This topology exhibits substantial cross-species alignment across conditions. Overlapping human and mouse modules suggest conservation of regulatory organization, consistent with the observation that widely utilized mouse models (e.g., KPC models) faithfully recapitulate the complex regulatory syntax of human pancreatic ductal adenocarcinoma (Hingorani et al., 2005).

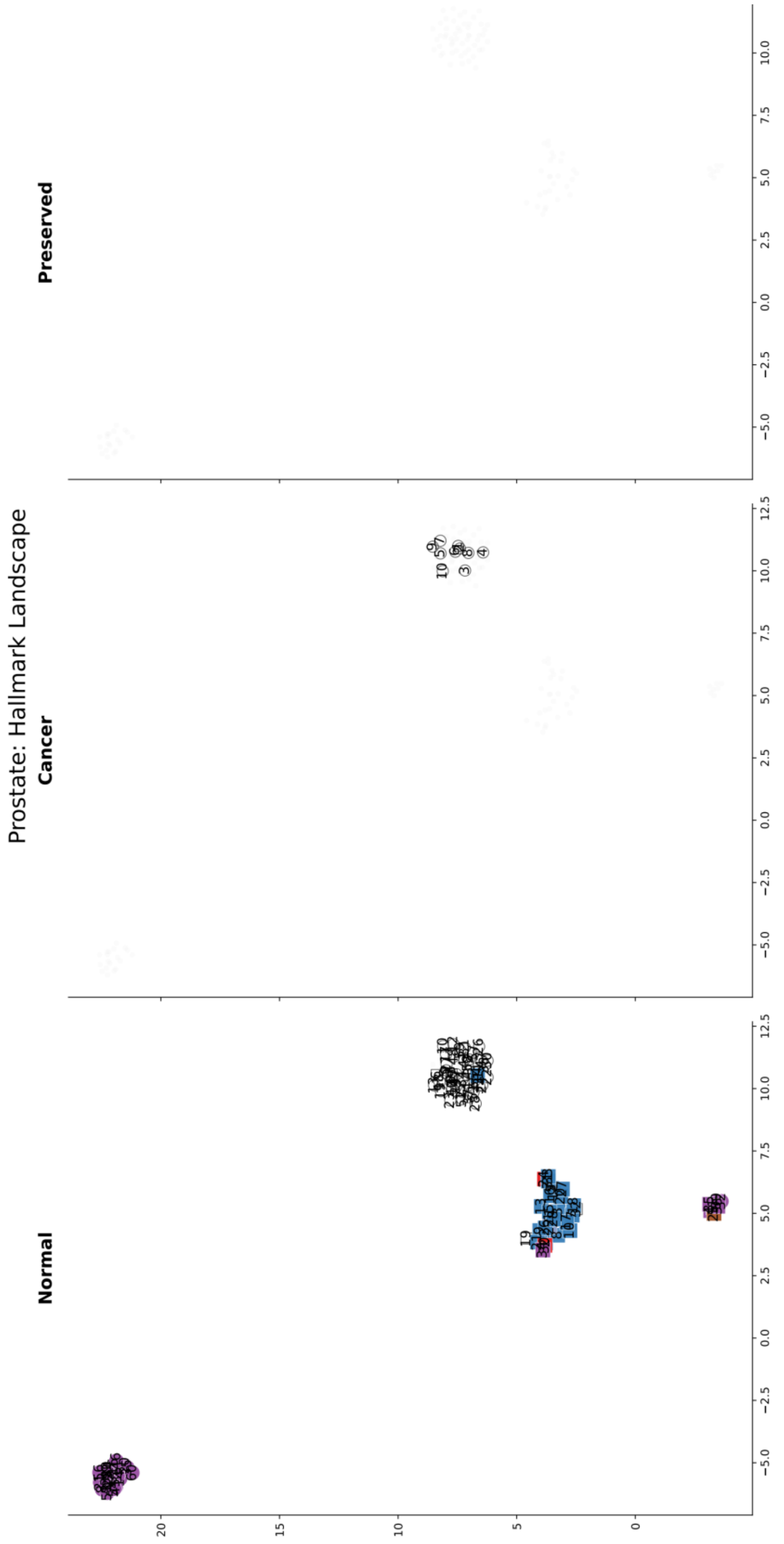


Supplementary Figure 3-12. UMAP projection of prostate cancer modules based on hallmarks. The highly segregated human and mouse modules in the Normal condition indicate divergent baseline regulatory networks. The loss of mouse *Deregulating cellular metabolism* modules may capture the "death" of the normal prostate metabolic factory: the normal prostate is a highly specialized metabolic factory designed to synthesize, accumulate, and secrete massive amounts of citrate (and zinc) into seminal fluid (by truncating the TCA cycle); when prostate cancer develops, it undergoes a fundamental "Metabolic Flip" to restore the TCA cycle to oxidize citrate for energy to fuel tumor growth. *Unclassified* human modules in Cancer condition are enriched for WikiPathways, including *mRNA processing* (WP411) and *Neovascularization processes* (WP4331), which reflects that prostate cancer mRNA processing is frequently dysregulated, particularly through alternative pre-mRNA splicing, and that the formation of new blood vessels is a critical process enabling prostate cancer growth and metastasis (Grizzi et al., 2023; Phillips et al., 2020).

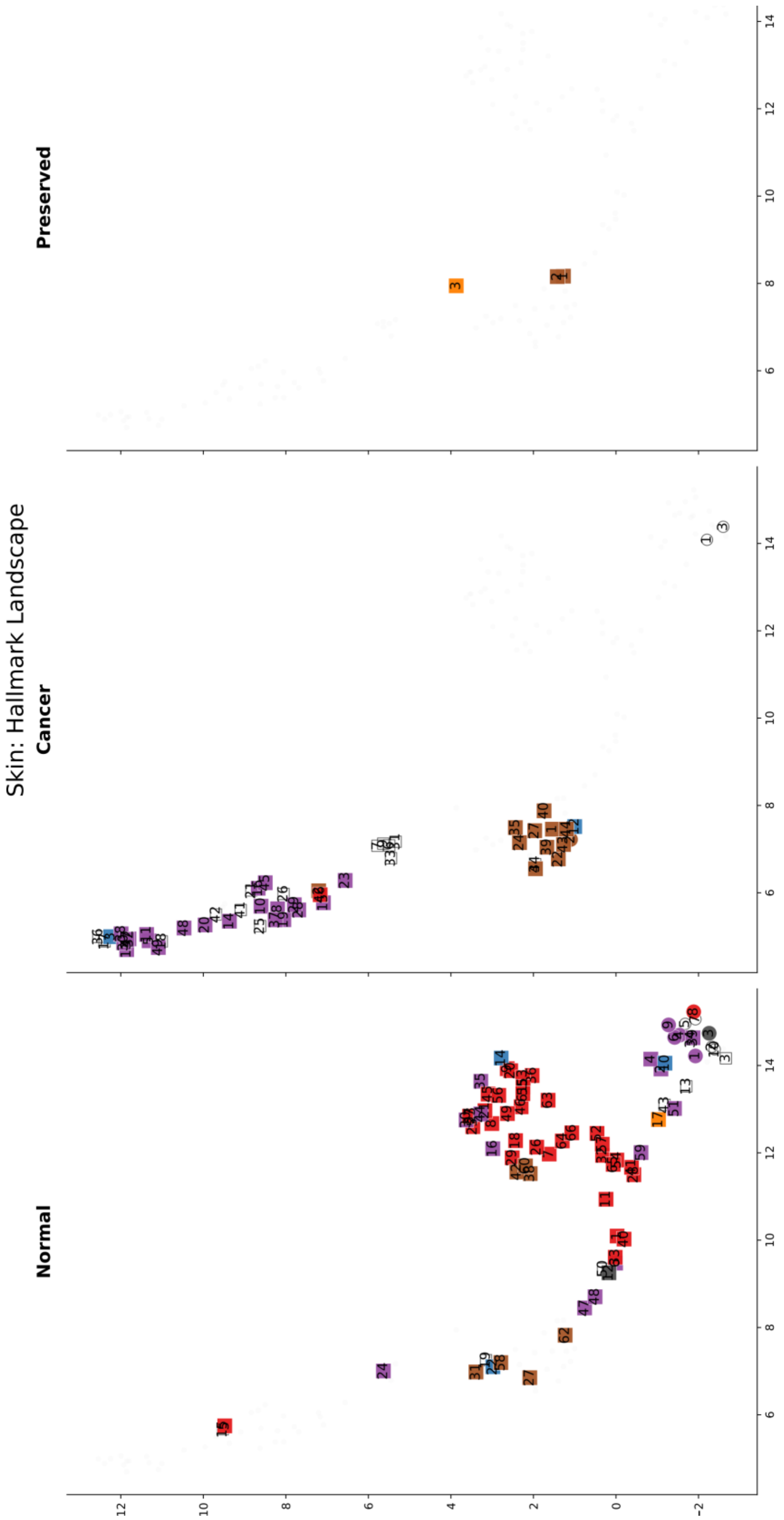


Supplementary Figure 3-13. UMAP projection of skin cancer modules based on hallmarks. This topology exhibits distinct species patterns. In the Normal condition, mouse modules exhibit regulatory complexity and likely reflect the highly synchronized, proliferative hair follicle cycling that is intrinsic to mouse skin but absent in human control tissues. In the Cancer condition, this normal mouse proliferative architecture is dismantled and replaced by *Deregulating cellular metabolism* and *Tumor-promoting inflammation* clusters, probably reflecting the metabolic reprogramming and immune-evasive dependencies characteristic of standard mouse melanoma models (e.g., BRAF/PTEN models) (Dankort et al., 2009). Conversely, human melanoma exhibits limited Cancer and Preserved modules, consistent with the UV-induced mutational burden and phenotypic plasticity (Rambow et al., 2019).

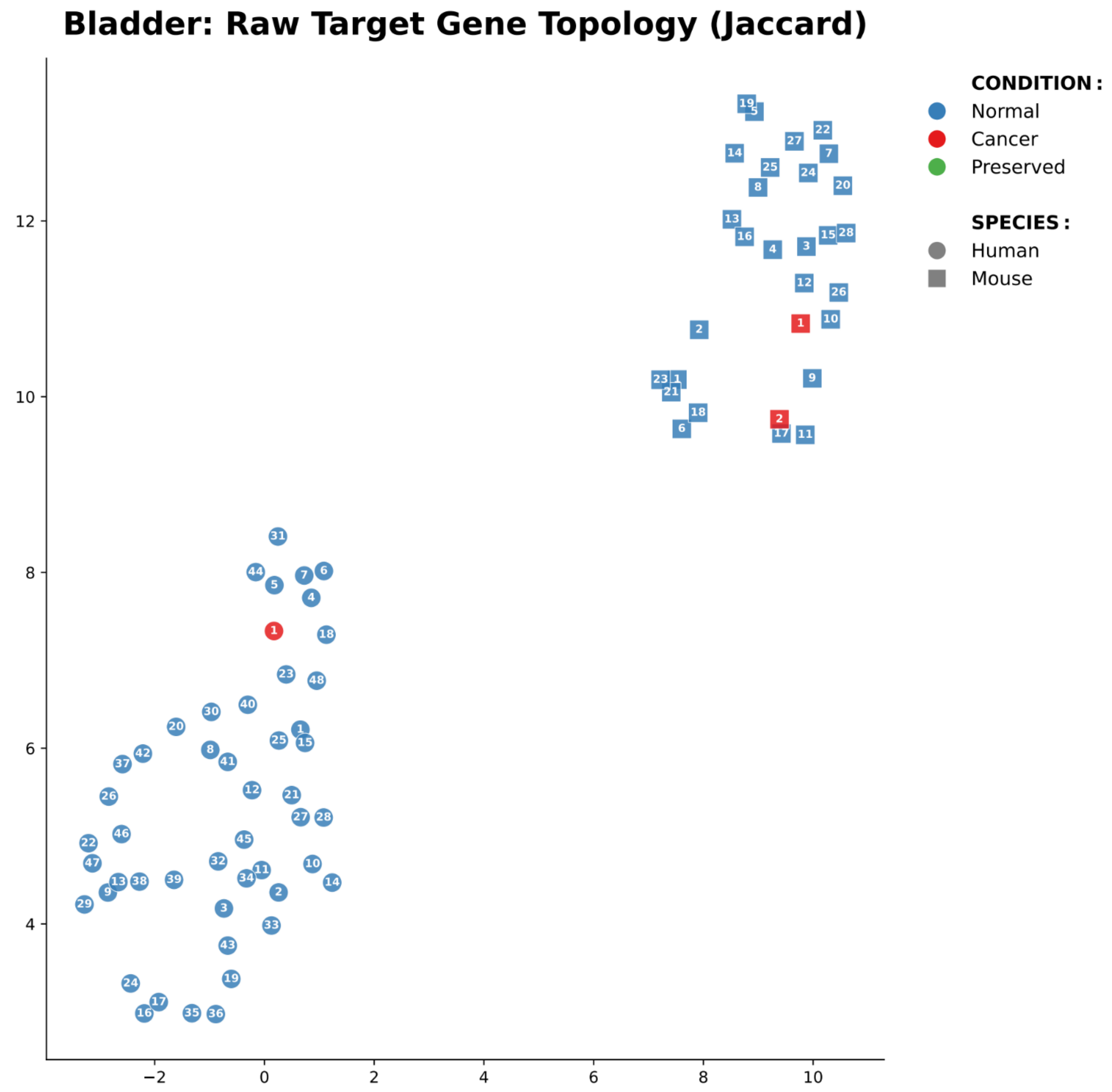


Supplementary Figure 4-1. UMAP projection of bladder cancer modules based on target gene Jaccard distances. This spatial landscape maps modules based entirely on the raw Jaccard overlap similarity between their downstream target gene sets, completely independent of predefined hallmarks or pathways. Circular and square nodes designate human and mouse modules; blue, red, and green coloring indicates Normal, Cancer, and Preserved conditions; and numbers indicate module identifiers. Within this topology, human and murine modules remain absolutely segregated, indicating fundamentally divergent non-coding regulatory architectures. The Preserved condition (green nodes) is absent in both species, and the diverse Normal module repertoire collapses into a sparse number of highly isolated Cancer modules, visually indicating catastrophic topological rewiring. This suggests that malignant transformation in the bladder dismantles normal homeostatic LS lncRNA programs and forces the invention of *de novo* regulatory networks for tumor survival.

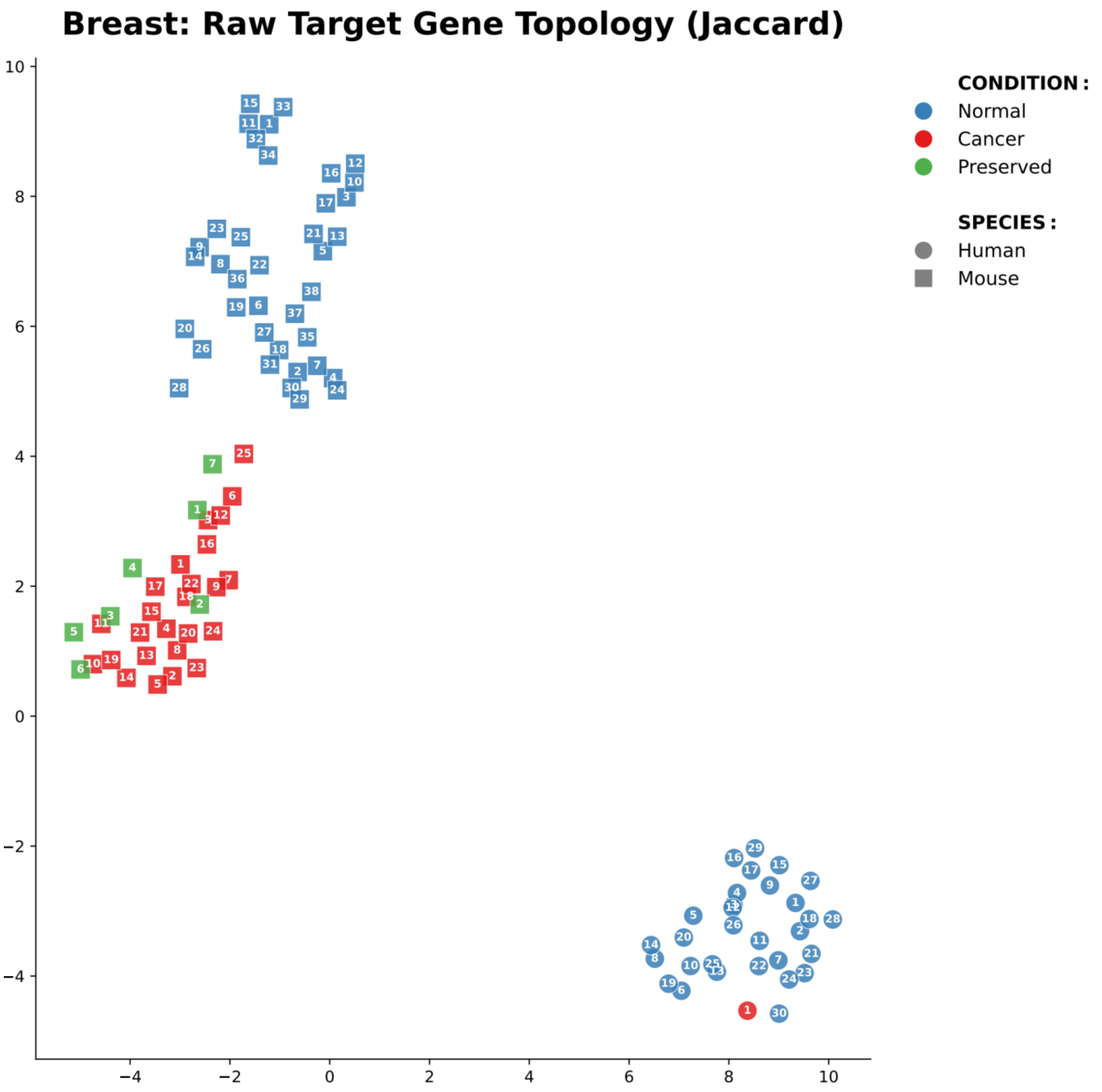


Supplementary Figure 4-2. UMAP projection of breast cancer modules based on target gene Jaccard distances. Within this topology, human and murine modules exhibit extreme spatial segregation, highlighting a fundamental divergence in their underlying regulatory architecture. This landscape visualizes two distinct etiologies of oncogenesis between the species. The murine landscape maintains a densely populated cluster of integrated Cancer and Preserved modules, indicating that standard preclinical breast cancer models survive through extensive "regulatory co-option" (maintaining and hijacking native pathways). In contrast, the human landscape completely lacks Preserved modules and has collapsed into a single identified Cancer module (node 1). This severe disparity suggests that human breast oncogenesis undergoes catastrophic topological rewiring, totally dismantling normal homeostatic programs and rendering murine regulatory dependencies highly non-representative of the human disease.

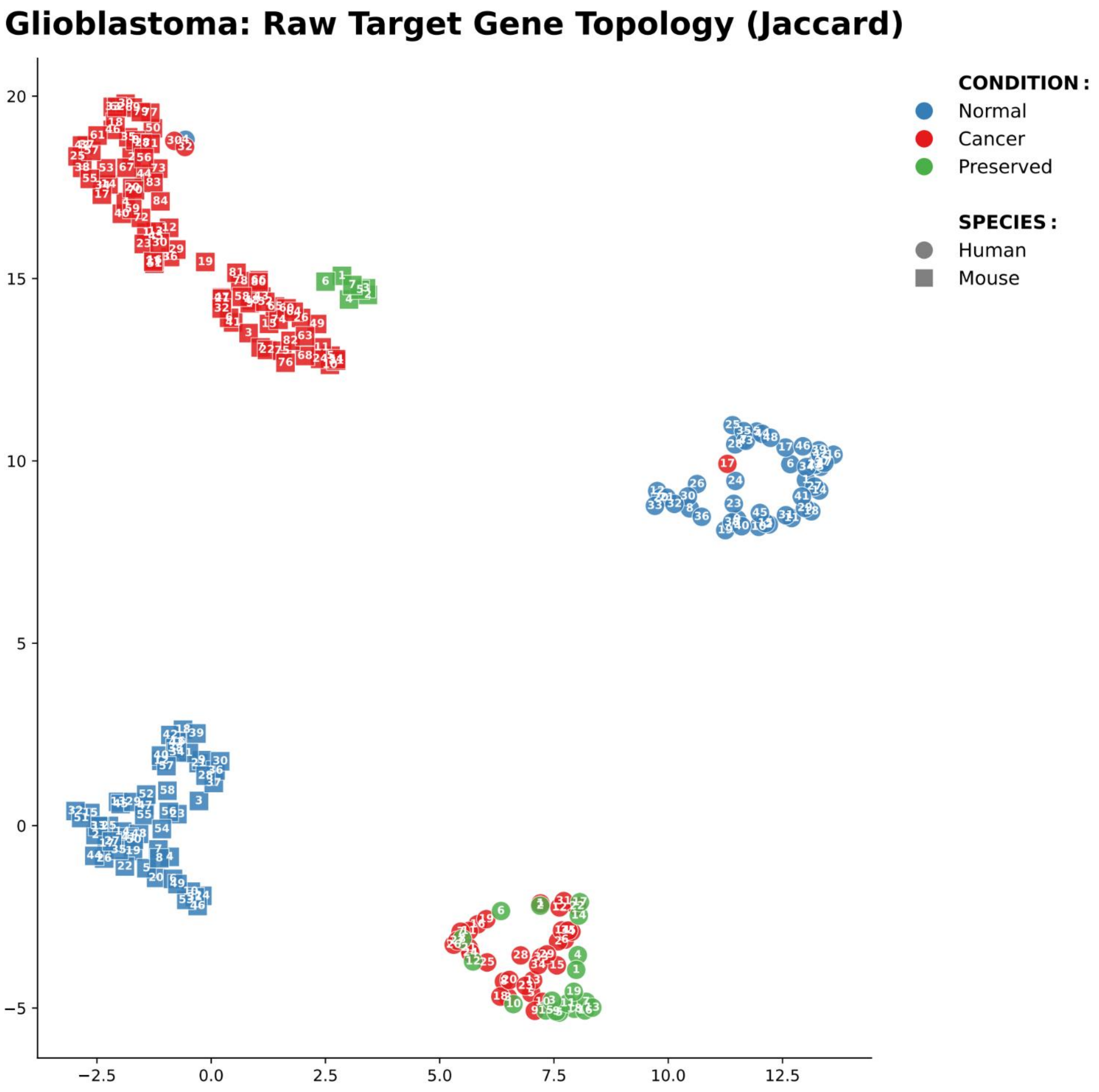


Supplementary Figure 4-3. UMAP projection of glioblastoma modules based on target gene Jaccard distances. Consistent with other evaluated solid tumors, human and murine modules remain completely spatially segregated. Within this structural framework, the two species exhibit highly divergent states of oncogenic transition. In the human landscape, Cancer and Preserved modules are densely intermingled to form a tight survival hub perfectly separated from the Normal state; this topology structurally reflects "regulatory co-option," wherein the human tumor survives by permanently hijacking tightly integrated developmental or epigenetic networks. Conversely, the murine landscape exhibits a massive spatial expansion of uniquely Cancer modules (top left) that have physically severed ties with both Normal and Preserved states, indicating extensive "topological rewiring" and the invention of sprawling *de novo* circuitry. Notably, distinct topographical anomalies—such as the massive over-proliferation of murine Cancer states, or the isolated presence of a single human Cancer module (node 17) that rigidly maintains a strictly Normal regulatory topology—highlight complex, multi-layered regulatory shifts. These unique topological features invite diverse biological and clinical interpretations regarding how different mammalian genomes uniquely navigate extreme neuro-oncogenic stress.

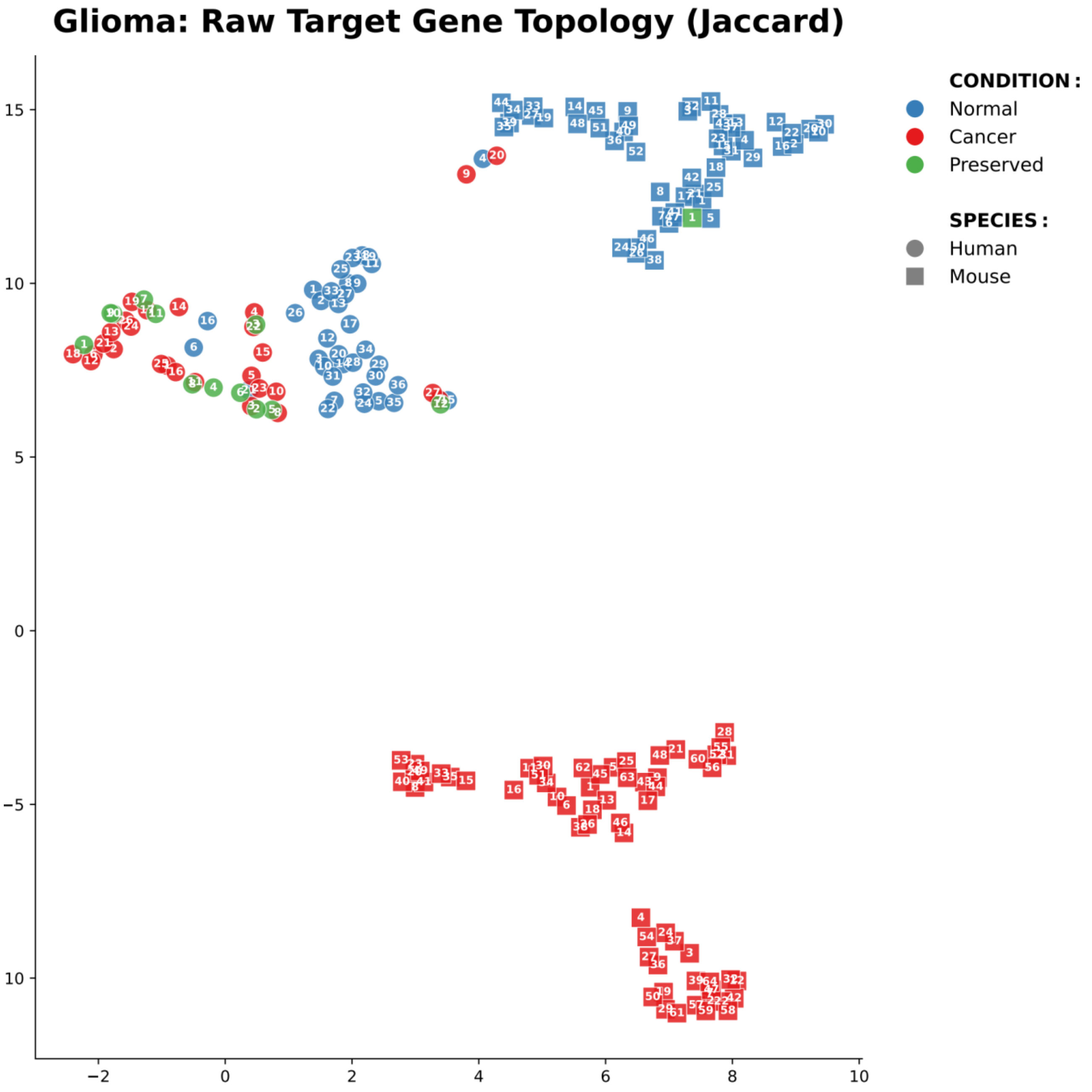


Supplementary Figure 4-4. UMAP projection of glioma modules based on target gene Jaccard distances. Within this topology, human and murine modules remain distinctly segregated. Notably, the spatial relationship between Normal and Cancer states differs markedly across the two species. The human landscape forms a highly continuous trajectory, where the Normal condition physically transitions into an intermixed, densely populated arc of Cancer and Preserved modules. This structural gradient suggests that human glioma progresses via graduated "regulatory co-option," retaining substantial native topologies throughout malignant transformation. Conversely, the murine landscape exhibits a stark discontinuous break: while murine Normal modules and the Preserved module (nodes 1) group tightly, the murine Cancer state has been structurally severed and fractured into highly isolated topological islands in the bottom area. This extreme separation visually captures severe "topological rewiring", accurately mirroring the etiology of standard murine glioma models generated via aggressive, artificial genetic alterations rather than continuous, natural tumor evolution. These distinct topologies invite further investigation into exactly which specific murine sub-clusters might best mimic the integrated human disease state.

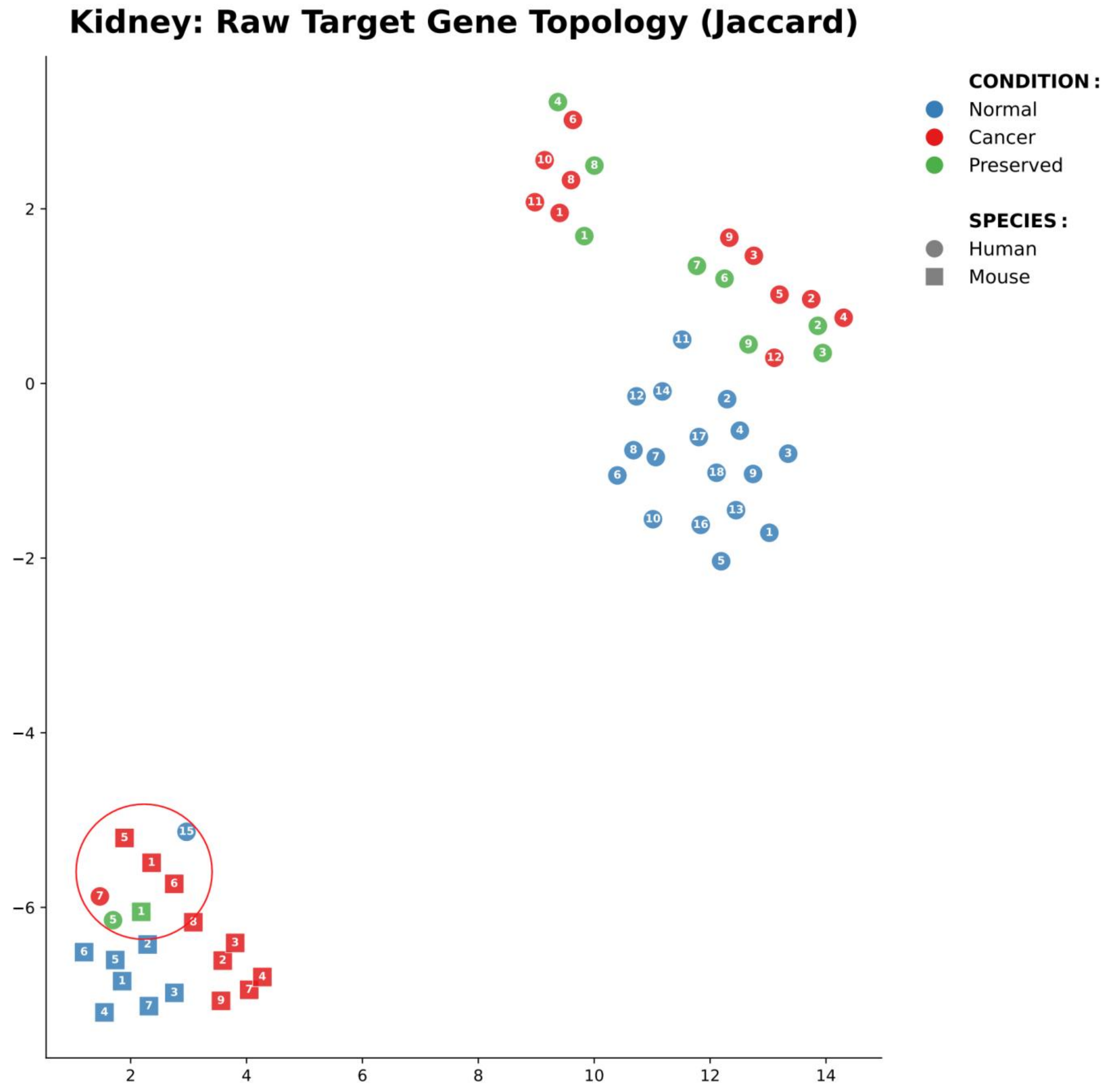


Supplementary Figure 4-5. UMAP projection of kidney cancer modules based on target gene Jaccard distances. Of note, three human modules (Normal ME15, Cancer ME7, and Preserved ME5) are close to the murine cluster. Notably, in the hallmark landscape (Figure 3B), human Cancer ME7 is with mouse Cancer modules and human Preserved ME5 is with mouse Preserved ME1, all having the hallmark of *Tumor-promoting inflammation*. This dual-landscape concordance of the exact same modules mathematically validates the entire analysis pipeline. They indicate that the mouse model uses the same orthologous genes to drive the same inflammatory phenotype. While the mouse fails at capturing the complex, multi-hit metabolic and proliferative machinery of human kidney cancer (the *Sustaining proliferative signaling* and *Resisting programmed cell death* modules in the hallmark landscape have no mouse equivalents), it perfectly captures the "*Tumor-promoting inflammation*" axis. This topology exhibits profound structural insights into the duality of animal models: what they fail at (the widely separated nodes in the two landscapes) and what they excel at (the tightly co-clustered nodes in the two landscapes).

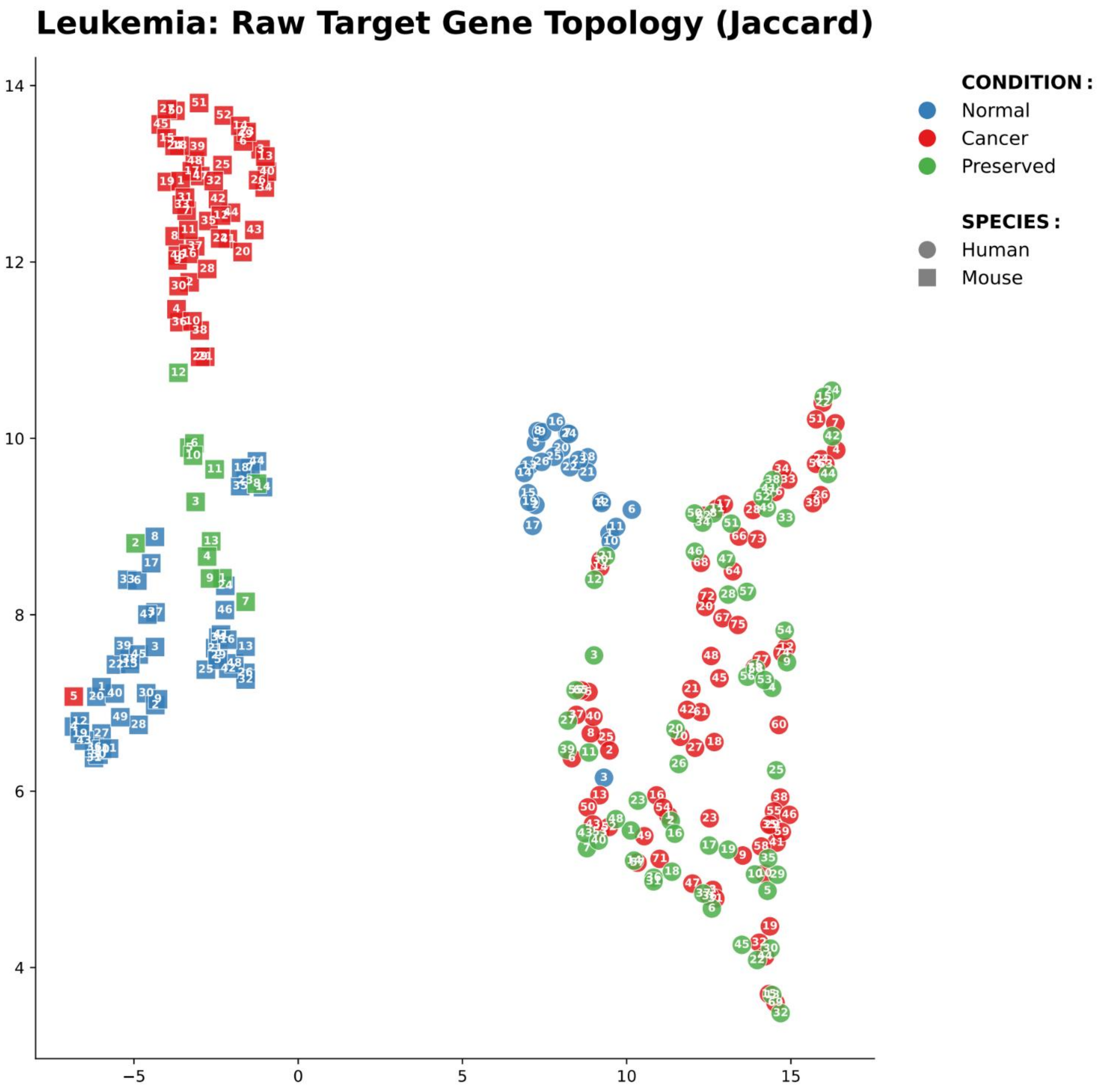


Supplementary Figure 4-6. UMAP projection of leukemia modules based on target gene Jaccard distances. This topology exhibits complete spatial segregation between the two species. This topology mathematically demonstrates that, while mammalian leukemogenesis converges on identically conserved physiological functions (Figure 3A), humans and mice orchestrate these hallmarks using strictly species-specific raw target gene networks downstream of LS lncRNAs. This supports a core thesis of the study: even when humans and mice have perfectly convergent macroscopic functions (Hallmarks), the actual raw gene syntax orchestrated by LS lncRNAs is entirely lineage-specific. Furthermore, the topography of both species is singularly defined by an extraordinarily high density of the Preserved modules (green nodes). This distinct topological preservation visually supports the etiology of "regulatory co-option," structurally confirming that leukemia survives not by inventing completely *de novo* signaling networks, but by heavily hijacking and permanently locking pre-existing hematopoietic and epigenetic programs.

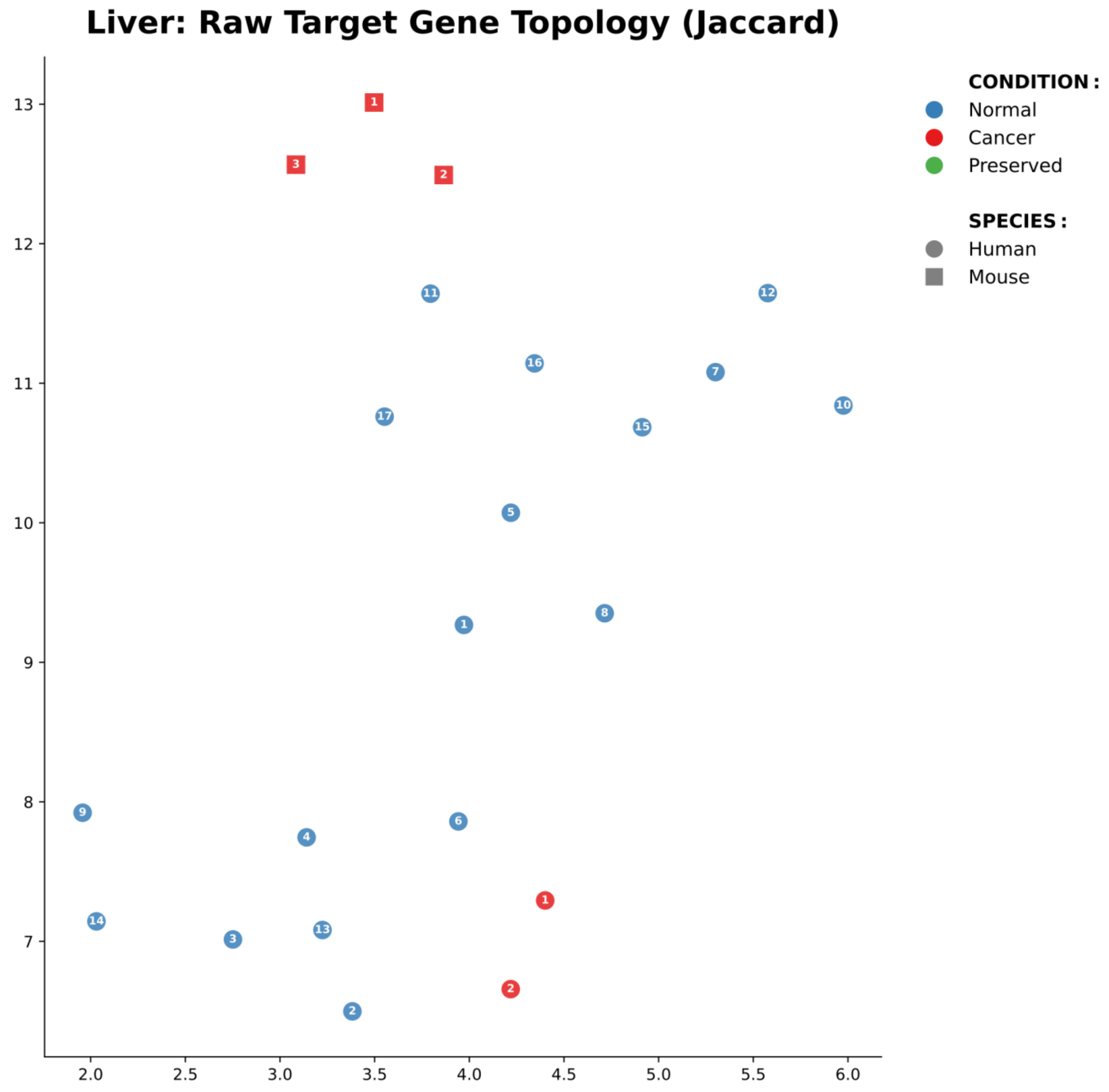


Supplementary Figure 4-7. UMAP projection of liver cancer modules based on target gene Jaccard distances. The liver cancer topology is characterized by severe regulatory collapse and complete species segregation. Both humans and mice display an entirely empty "Preserved" landscape (zero green nodes). This visually indicates that hepatocellular transformation involves catastrophic "topological rewiring," completely dismantling normal hepatic programs without retaining any baseline homeostatic circuitry—a dynamic consistent with clinical hepatocellular carcinoma, which typically arises by obliterating normal liver architecture through chronic cirrhosis. In the human landscape (circles), the diverse, dispersed regulatory architectures defining the Normal state undergo massive functional contraction, collapsing into merely two highly isolated Cancer modules. Furthermore, the murine landscape (squares) is extraordinarily sparse, with only three disconnected Cancer modules and no structurally coherent Normal regulatory modules. This extreme regulatory poverty mathematically underscores a major translational limitation: standard murine models of liver cancer operate using a drastically simplified transcriptional machinery that fundamentally fails to mirror the rich, complex regulatory topography of the human disease.

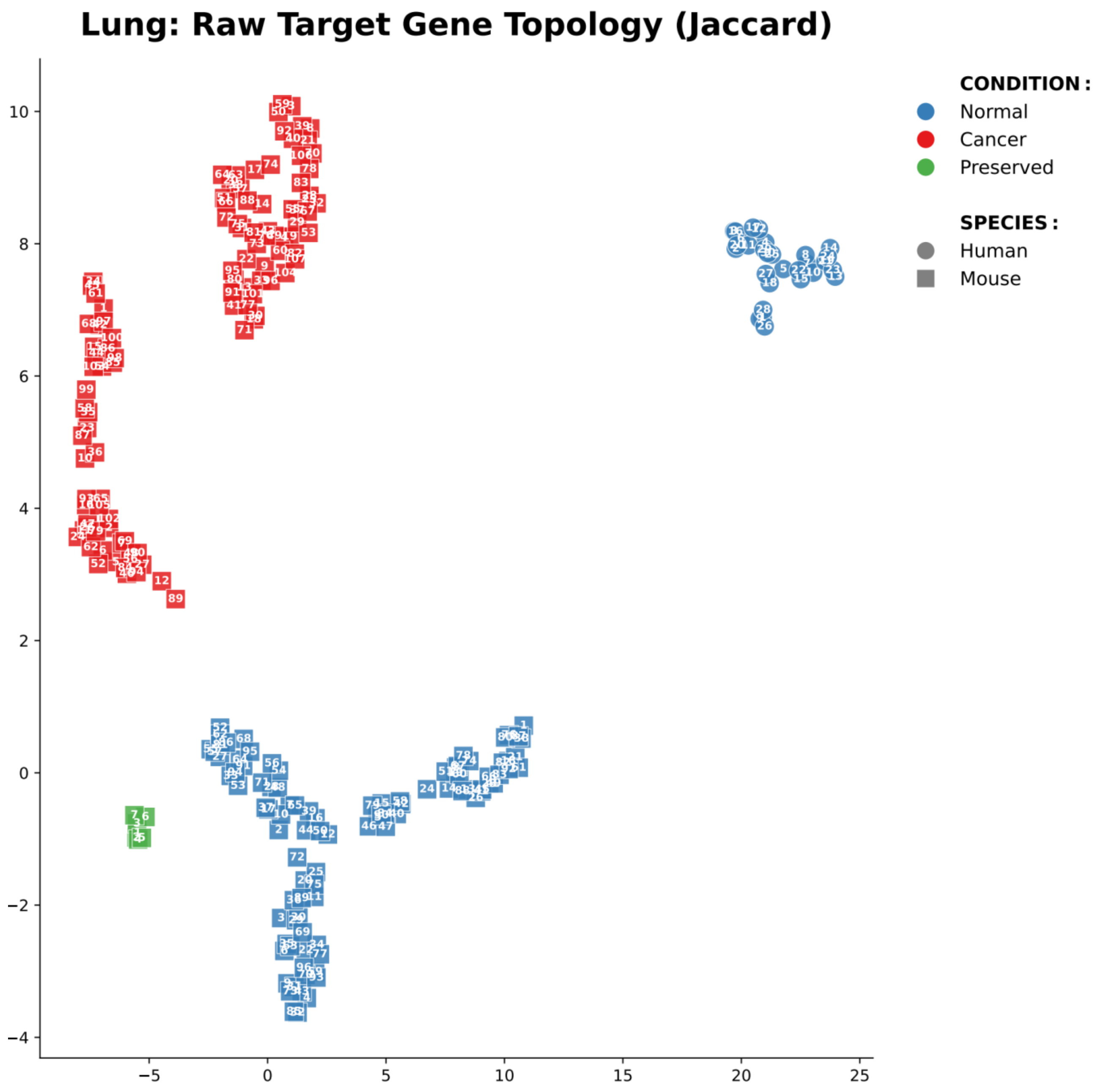


Supplementary Figure 4-8. UMAP projection of lung cancer modules based on target gene Jaccard distances. The lung cancer topology provides a striking visualization of species-specific regulatory plasticity. Extreme spatial segregation exists between humans and mice. The murine landscape is overwhelmingly populated, exhibiting vast, structurally complex domains of Normal and Cancer modules. This structural overabundance provides direct mathematical visualization of the uniquely high developmental plasticity often observed in engineered murine lung cancer models (Chan et al., 2026), equipping the mouse tumor with an expansive regulatory toolkit to drive therapy resistance. In dramatic contrast, the human landscape is confined strictly to Normal modules; cohesive Cancer and Preserved regulatory modules are entirely absent. This complete regulatory "dropout" implies that human lung oncogenesis—which typically evolves over decades of catastrophic, carcinogen-induced mutational stress—structurally shatters cohesive native LS lncRNA architectures, resulting in a fundamentally different, biologically barren non-coding topology compared to the hyperplastic murine disease.

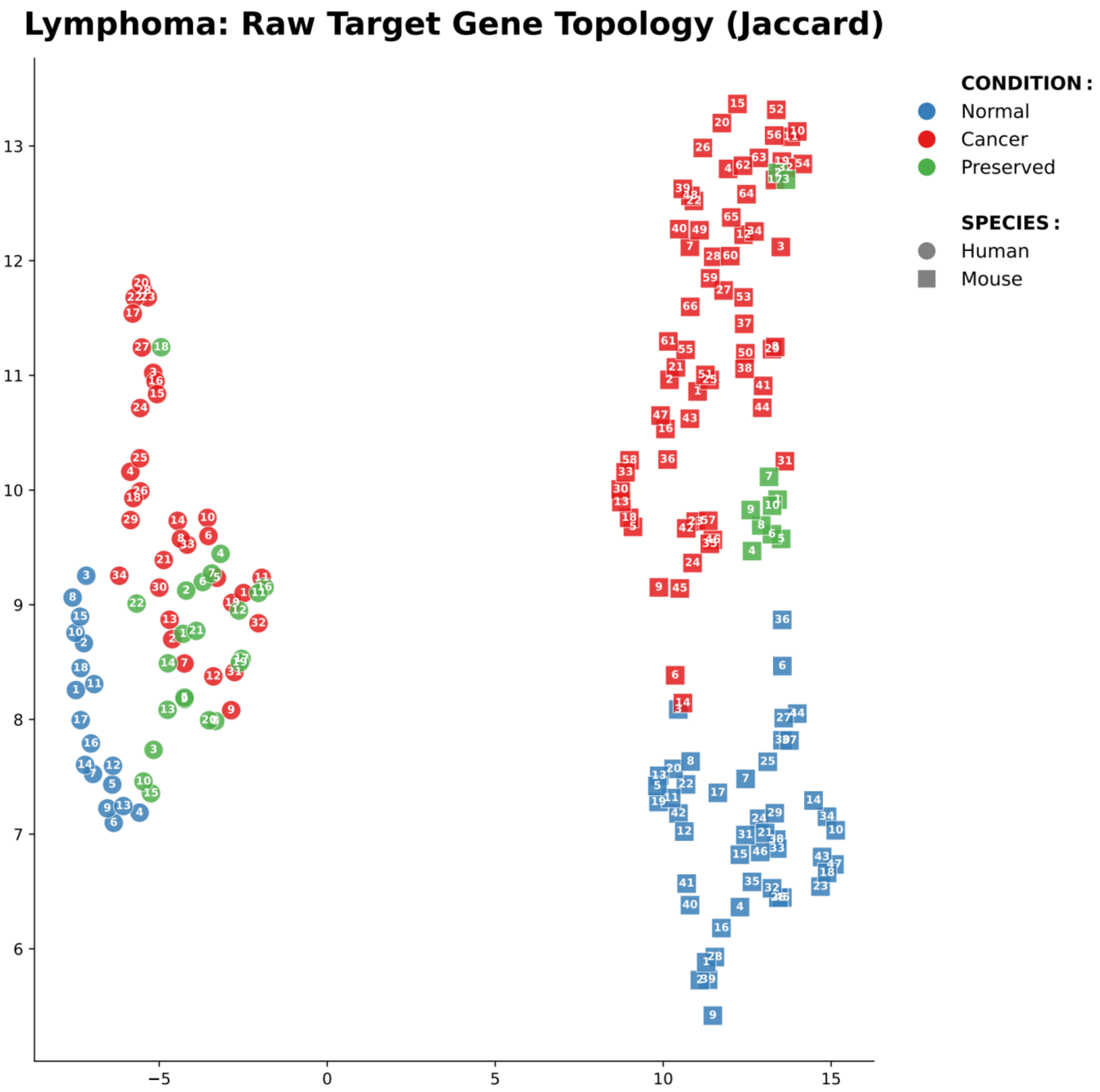


Supplementary Figure 4-9. UMAP projection of lymphoma modules based on target gene Jaccard distances. This spatial landscape maps modules based entirely on the raw Jaccard overlap similarity between their downstream target gene sets, independent of predefined hallmark annotations. Circular and square nodes designate human and mouse modules, respectively. Blue, red, and green indicate Normal, Cancer, and Preserved biological conditions, with numbered labels corresponding to specific module identifiers. Within the lymphoma topology, human and murine modules maintain strict spatial segregation, demonstrating that despite sharing broad hematologic cancer hallmarks, their foundational LS lncRNA target architectures remain highly lineage-specific. Notably, the two species map distinctly different trajectories of oncogenic transformation. The human landscape (circles) forms a highly continuous structural gradient; Normal modules integrate smoothly into a dense transitional bridge of Preserved modules, which are deeply intimately intermingled with Cancer modules. This cohesive topography visually represents classic "regulatory co-option," whereby human lymphoma naturally and progressively evolves by hijacking deeply entrenched lymphoid homeostatic networks. Conversely, the murine landscape (squares) shows a massive topological expansion of isolated Cancer networks, stretching far from the Normal baseline. This expansive murine Cancer state points toward extreme "topological rewiring"—a structural dynamic highly characteristic of standard preclinical models forced into rapid malignancy via aggressive, engineered alterations rather than the continuous, stepwise genomic evolution observed in human patients.

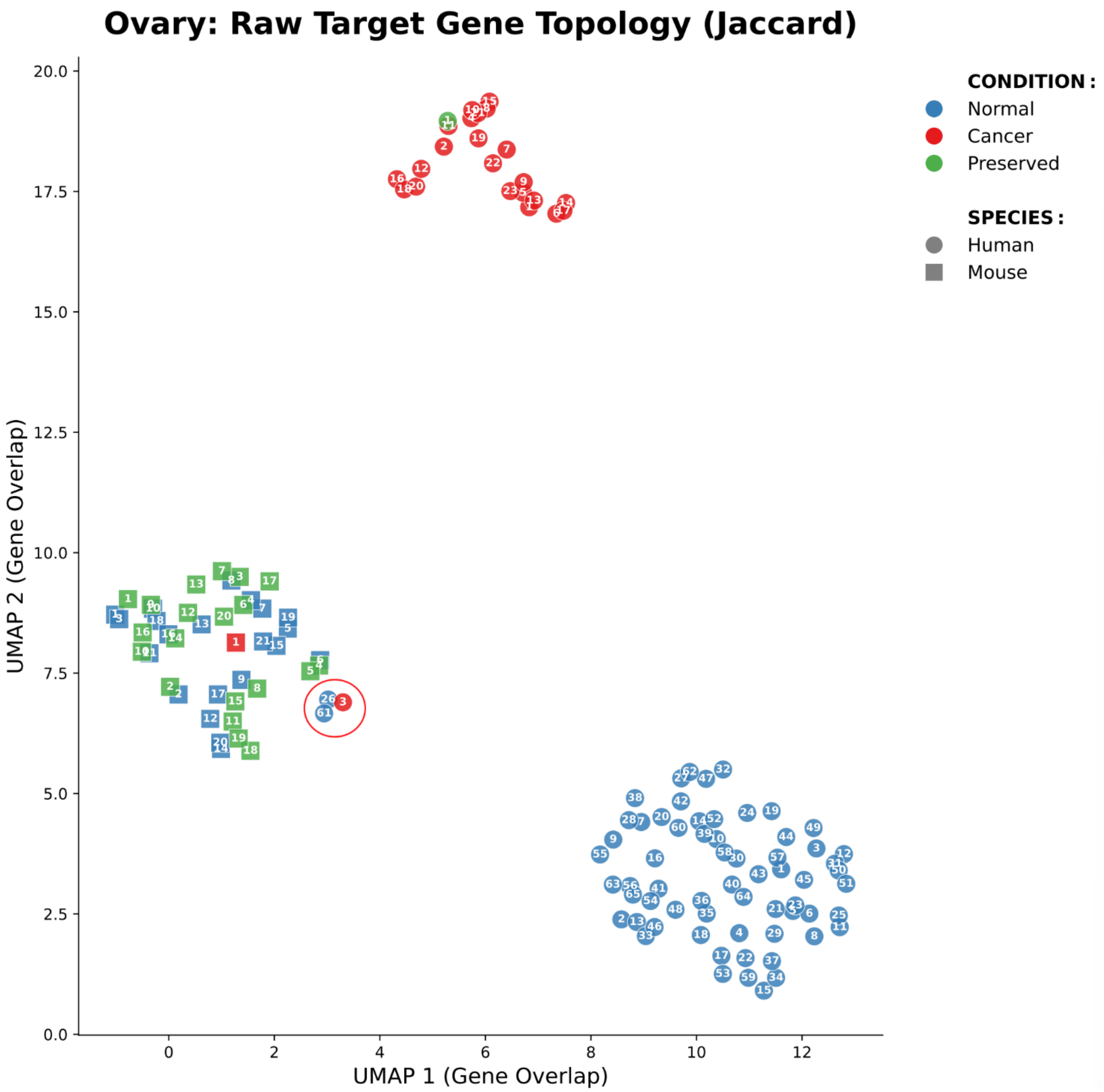


Supplementary Figure 4-10. UMAP projection of ovarian cancer modules based on target gene Jaccard distances. The topology of ovarian cancer mathematically visualizes profound cross-species divergence in epithelial tumor biology, contrasted against deeply conserved microenvironmental responses. The human landscape displays a topological fracture: the dense Normal baseline (bottom right) is structurally severed from the Cancer/Preserved cluster (top center), reflecting catastrophic epithelial topological rewiring—a dynamic consistent with the profound DNA repair dysfunction, *p53* loss, and extensive homologous recombination deficiency driving human high-grade serous ovarian carcinoma. Conversely, the murine landscape exhibits tight, cohesive topological integration across Normal, Preserved, and Cancer conditions, underscoring how standard murine models rely on intact, heavily co-opted classical growth factor pathways (e.g., EGFR and estrogen signaling) rather than genome-instability-driven survival. Strikingly, despite this extreme epithelial divergence, highly specific translatable networks co-cluster within the murine manifold (red circle). These physically embedding human modules (Cancer ME3, Normal ME26/61) are heavily defined by canonical stromal, EMT, and angiogenic effectors (e.g., *ACTA2*, *ZEB1*, *FLT4*). This precise co-localization proves that while standard murine models structurally fail to recapitulate the human epithelial genomic collapse, they remarkably successfully conserve the host tumor microenvironment and angiogenic response, thereby defining the optimal pre-clinical window for specific therapeutic testing.

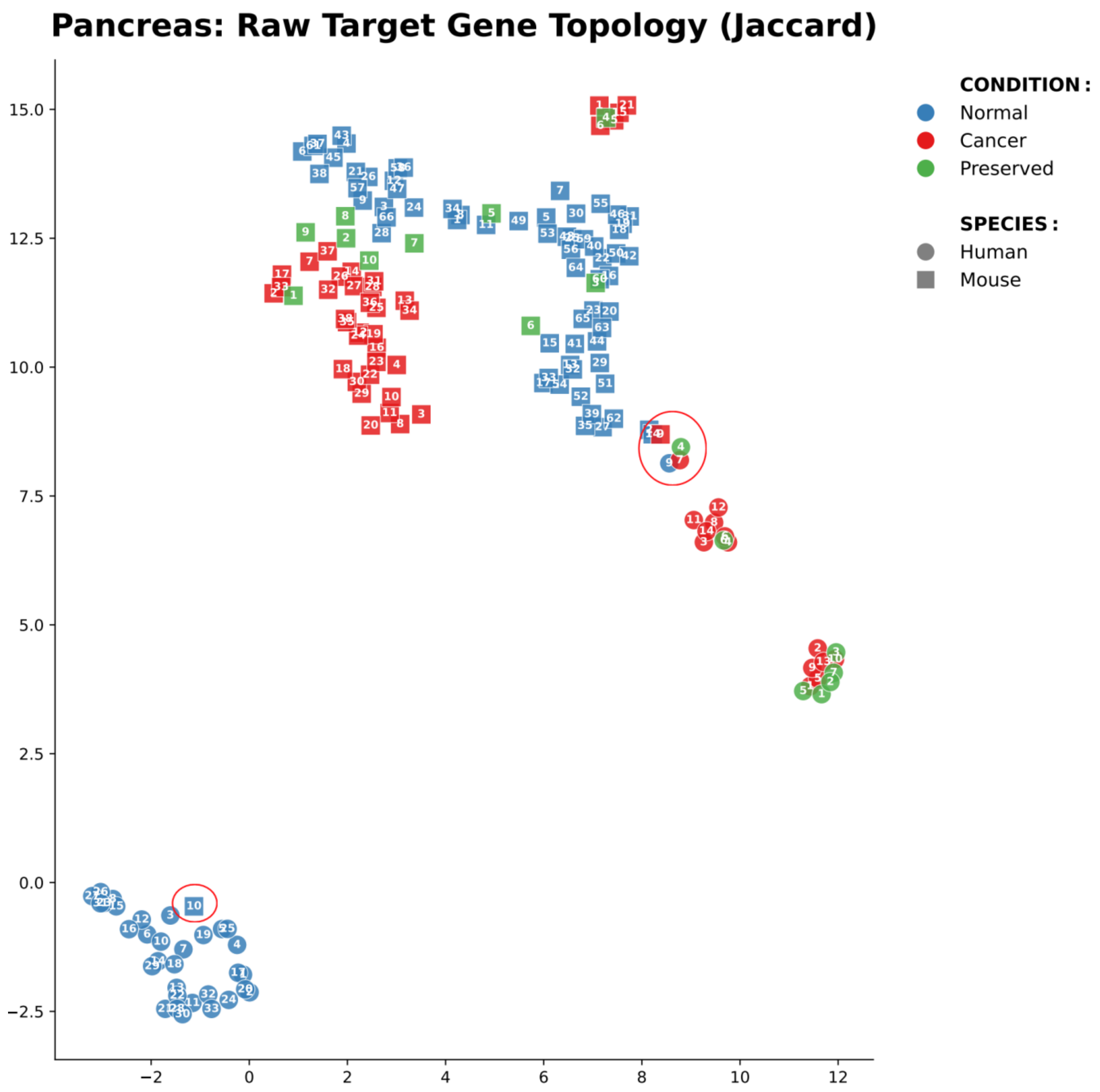


Supplementary Figure 4-11. UMAP projection of pancreatic cancer modules based on target gene Jaccard distances. This spatial landscape maps modules based entirely on the raw Jaccard overlap similarity between their downstream target gene sets, independent of predefined hallmark annotations. Circular and square nodes designate human and mouse modules, respectively. Blue, red, and green indicate Normal, Cancer, and Preserved biological conditions, with numbered labels corresponding to specific module identifiers. Red circles highlight specific spatial coordinates of acute cross-species convergence. The overall pancreatic architecture reveals a fascinating topological reversal compared to other solid tumors. The murine landscape (squares) displays an incredibly cohesive, massive structural continuum bridging Normal, Preserved, and Cancer states. This uniquely integrated murine topography visually corroborates the well-established experimental fidelity of advanced, genetically engineered PDAC models (e.g., the KPC model), which are explicitly designed to mirror the slow, continuous progression from pre-malignant lesions to invasive carcinoma via sequential "regulatory co-option." Conversely, the human landscape (circles) features a stark structural fracture totally severing the Normal baseline cluster from the emerging sequence of Cancer and Preserved modules, reflecting the catastrophic "topological rewiring" and extreme stromal/metabolic shifts that define natural human disease evolution. Despite this macro-level divergence, highly specific network nodes structurally converge (red circles). A murine Normal module precisely embeds within the dense human Normal cluster, denoting a deeply conserved baseline physiological identity. Crucially, a specific subset of murine Cancer and Preserved modules perfectly co-cluster within the fractured human Cancer trajectory. This strict mathematical convergence physically identifies the exact underlying regulatory pathways that successfully transcend species boundaries, highlighting the most robustly translatable axes for preclinical therapeutic intervention in pancreatic adenocarcinoma.

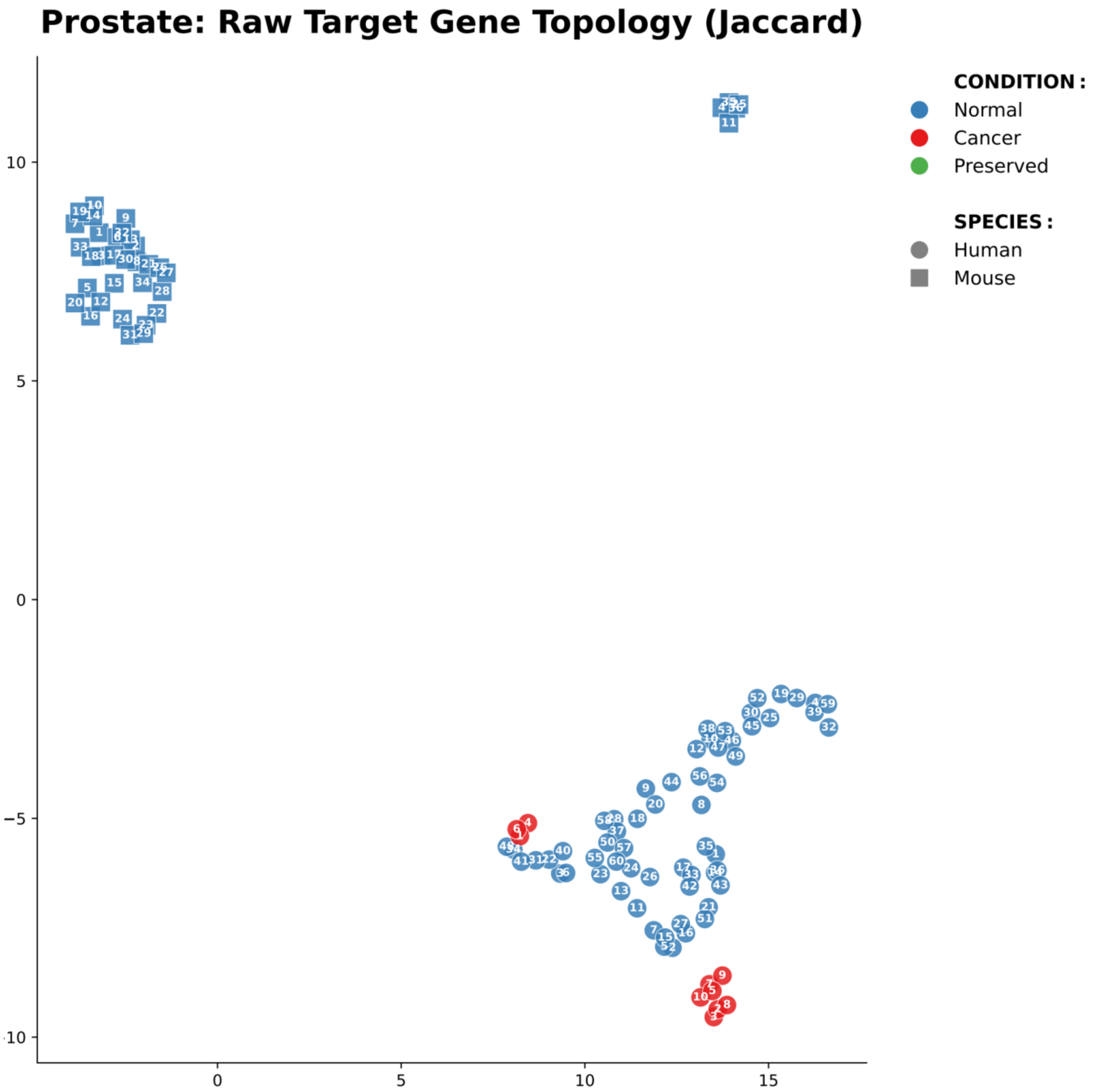


Supplementary Figure 4-12. UMAP projection of prostate cancer modules based on target gene Jaccard distances. This spatial landscape maps modules based entirely on the raw Jaccard overlap similarity between their downstream target gene sets, independent of predefined hallmark annotations. Circular and square nodes designate human and mouse modules, respectively. Blue, red, and green indicate Normal, Cancer, and Preserved biological conditions, with numbered labels corresponding to specific module identifiers. The prostate cancer topology reveals extreme structural disparity between species, underscored by a profound regulatory collapse during malignant transformation. The two species display complete spatial segregation, confirming deeply divergent baseline regulatory architectures. The most striking topological feature is the absolute absence of structurally cohesive murine Cancer (red) or Preserved (green) modules; the murine landscape is populated entirely by Normal regulatory networks. This catastrophic regulatory "dropout" mathematically visualizes a widely recognized biological translational barrier: because mice do not naturally develop prostate cancer, engineered murine models utilize highly artificial pathways that fundamentally fail to construct the complex, cohesive noncoding regulatory networks characterizing genuine tumorigenesis. Conversely, the human landscape features a massive, interconnected Normal topological web that undergoes severe structural contraction upon transformation, generating only two highly isolated, small Cancer clusters. The complete lack of Preserved modules across both species confirms that prostate oncogenesis requires severe "topological rewiring," completely dismantling native homeostatic pathways to assemble restricted, *de novo* oncogenic regulatory programs.

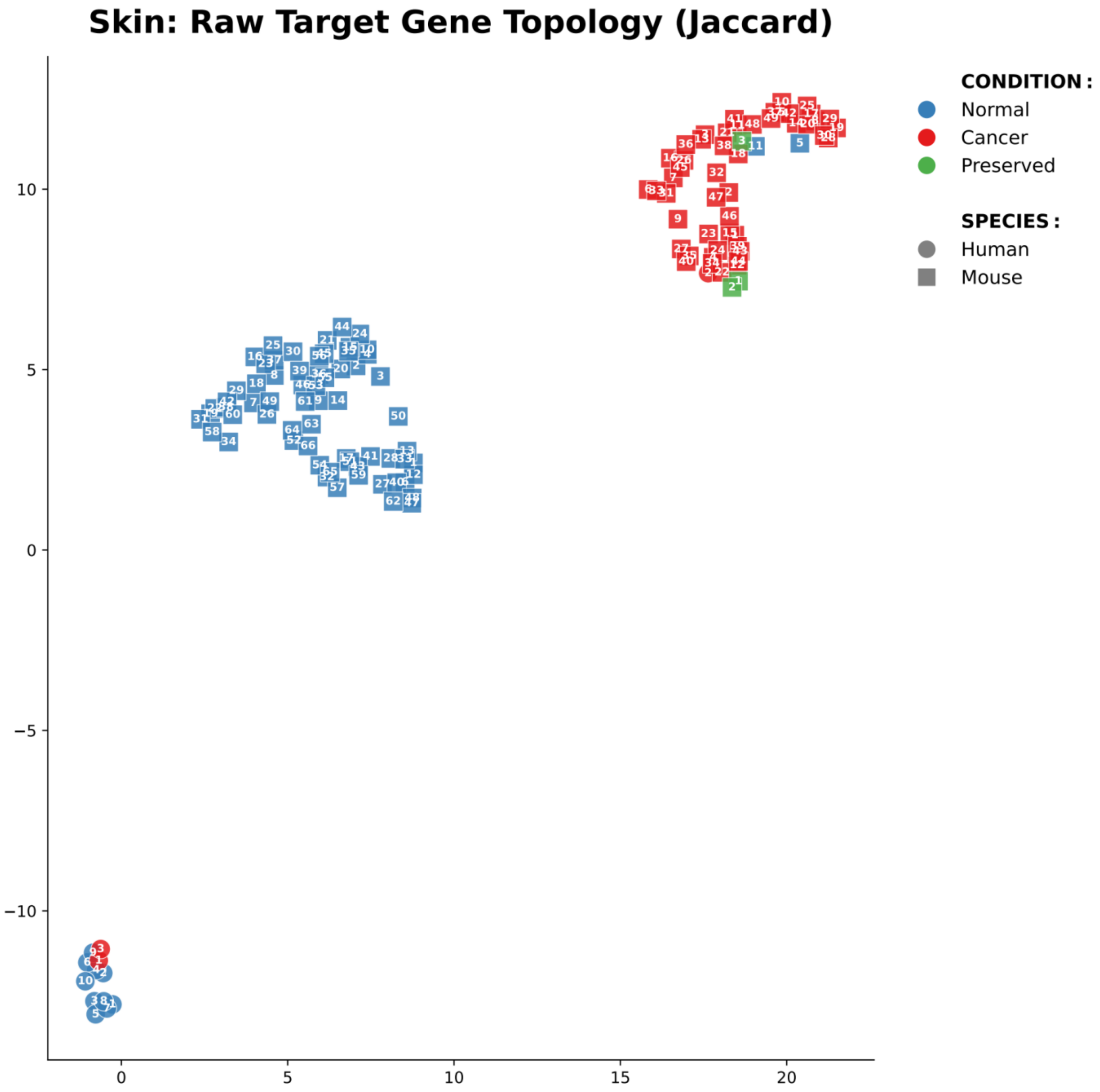


Supplementary Figure 4-13. UMAP projection of skin cancer (melanoma) modules based on target gene Jaccard distances. This spatial landscape maps modules based entirely on the raw Jaccard overlap similarity between their downstream target gene sets, independent of predefined hallmark annotations. Circular and square nodes designate human and mouse modules, respectively. Blue, red, and green indicate Normal, Cancer, and Preserved biological conditions, with numbered labels corresponding to specific module identifiers. The melanoma topology is characterized by absolute species segregation and starkly contrasts natural tumor evolution with engineered model genesis. In the human landscape (circles in the bottom left), regulatory networks exhibit severe structural contraction. A tightly clustered subset of Normal modules violently fragments into an extremely small repertoire of Cancer modules, with complete absence of topological preservation (zero green nodes). This structural obliteration mathematically mirrors the etiology of human melanoma, wherein catastrophic, UV-induced hypermutation extensively shatters normal homeostatic regulatory syntaxes, forcing extreme *de novo* topological rewiring. In extraordinary contrast, the murine landscape (squares) features immense structural capacity but a massive geometric divide. A sprawling "continent" of normal murine regulatory networks has completely physically severed from a similarly expansive continent of murine Cancer modules. This profound spatial discontinuity perfectly visualizes the mechanism of typical transgenic murine melanoma models (e.g., forced *BRAF*/*PTEN* alterations): the introduction of extreme, artificial oncogenic payloads forcefully snaps the biological system out of its native regulatory space, triggering an immediate and isolated transcriptional explosion completely unrepresentative of the heavily constricted, progressive structural degradation seen in human disease.

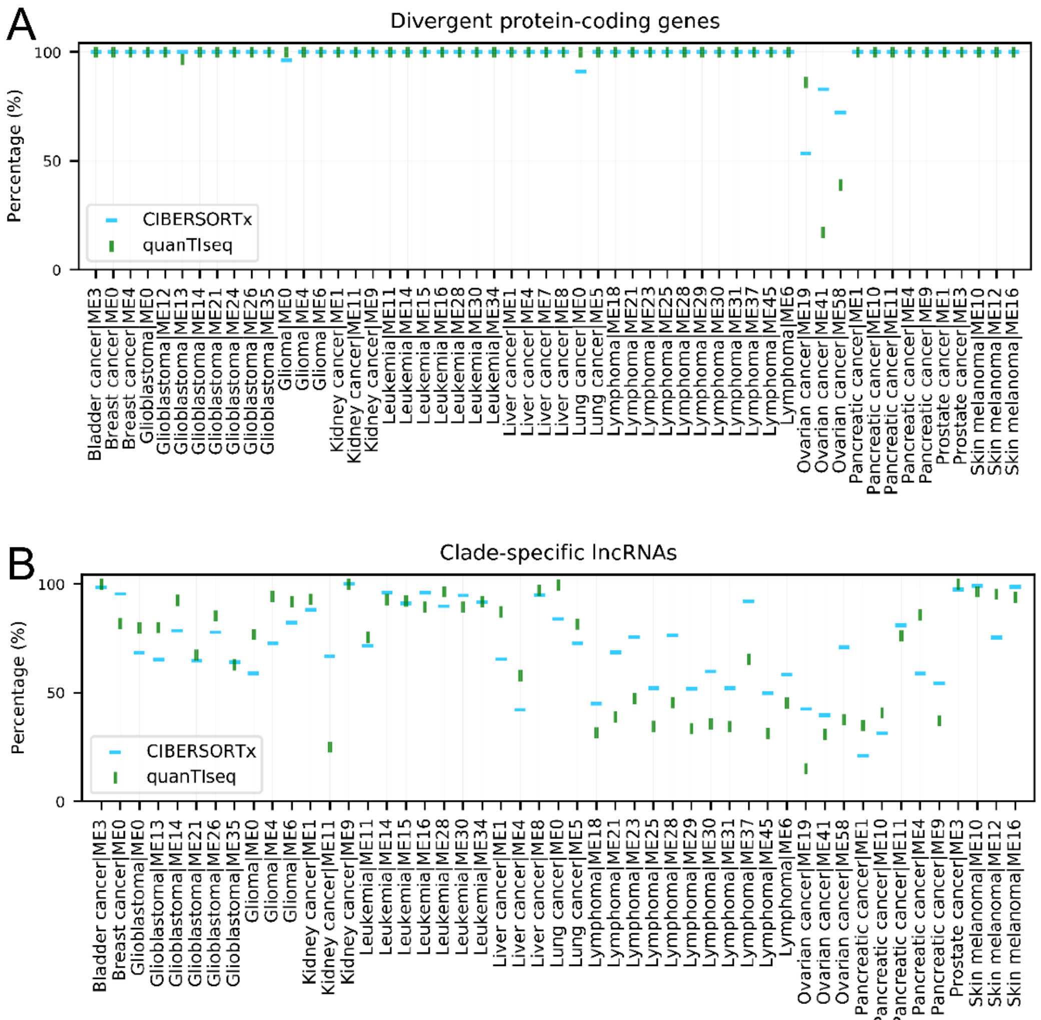


Supplementary Figure 5-1. NX values of 98% of TDGs and 70% of LS lncRNAs within immune divergence-associated modules (ID-associated modules) are significantly correlated with infiltration levels of at least one immune cell type (FDR < 0.05). (A) Percentage of TDGs showing significant correlations. (B) Percentage of LS lncRNAs showing significant correlations. Immune cell fractions were estimated using *CIBERSORTx* and *quanTIseq*.

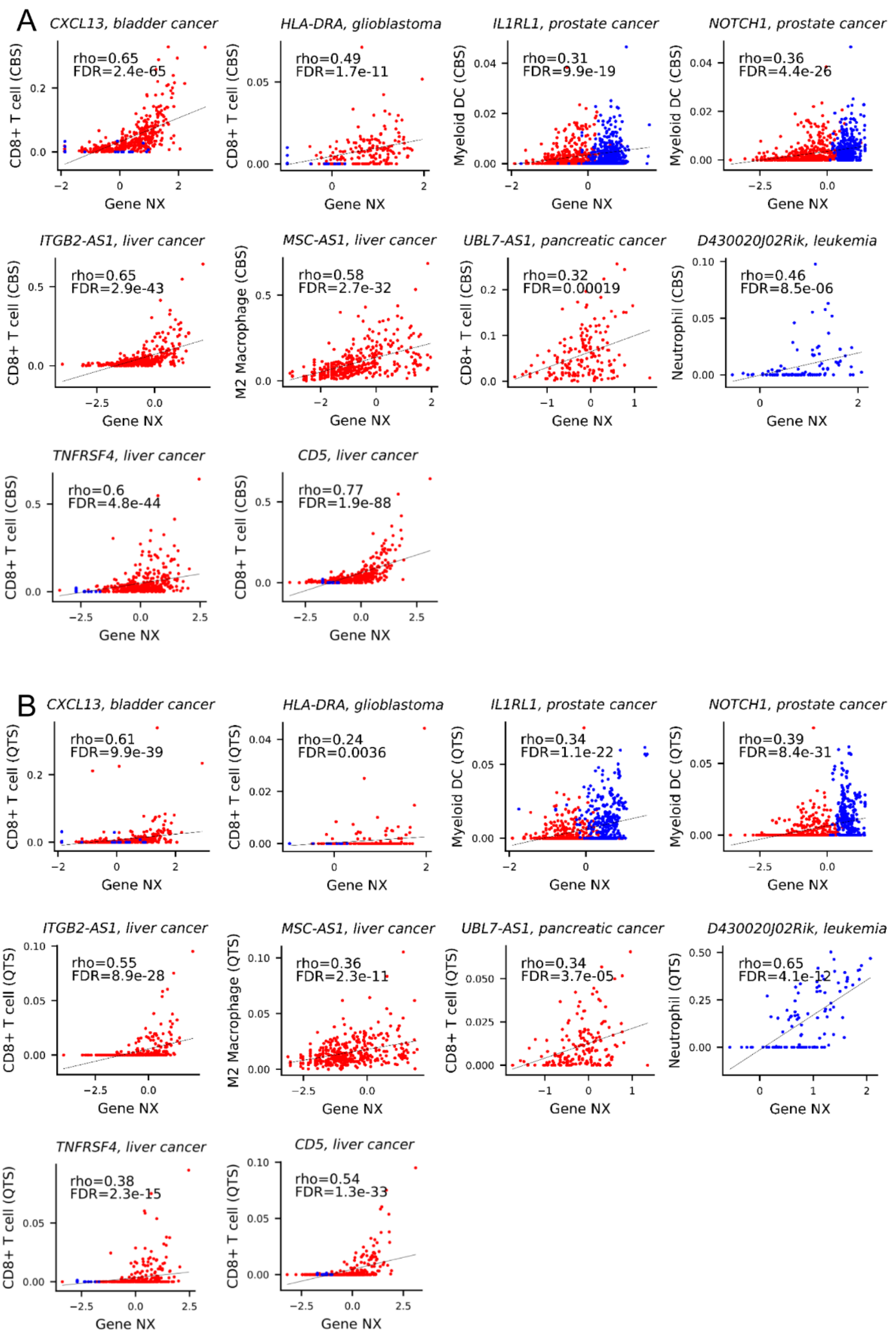


Supplementary Figure 5-2. Scatter plots illustrating correlations between TDG NX values and immune cell infiltration. Vertical axes indicate estimated immune cell abundance (*CIBERSORTx* or *quanTIseq*); horizontal axes indicate gene NX values. Red and blue dots represent human and mouse samples, respectively. (A) The results based on the *CIBERSORTx*. (B) The results based on the *quanTIseq*.

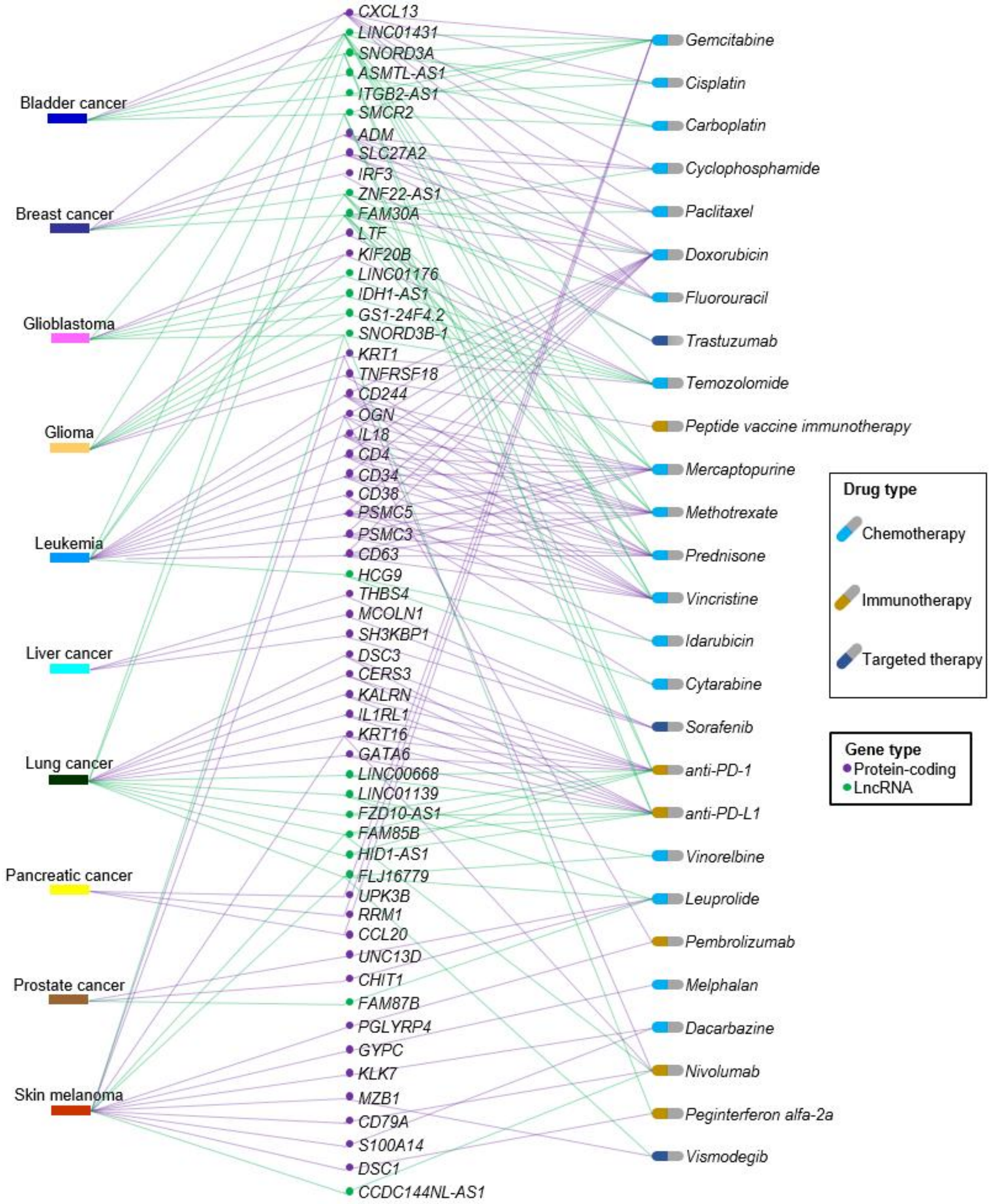


Supplementary Figure 6-1. Examples of tumors, LS lncRNAs (green lines and dots), TDGs (purple lines and dots), and anti-cancer drug resistance. Many TDGs with DBSs of PS lncRNAs (e.g., *DSC3*, *KRT16*, and *CERS3*, and FZD10-AS1, LINC00668, and HID1-AS1) are targets of anti-cancer drugs (e.g., anti-PD-1/PD-L1 therapy for lung cancer).

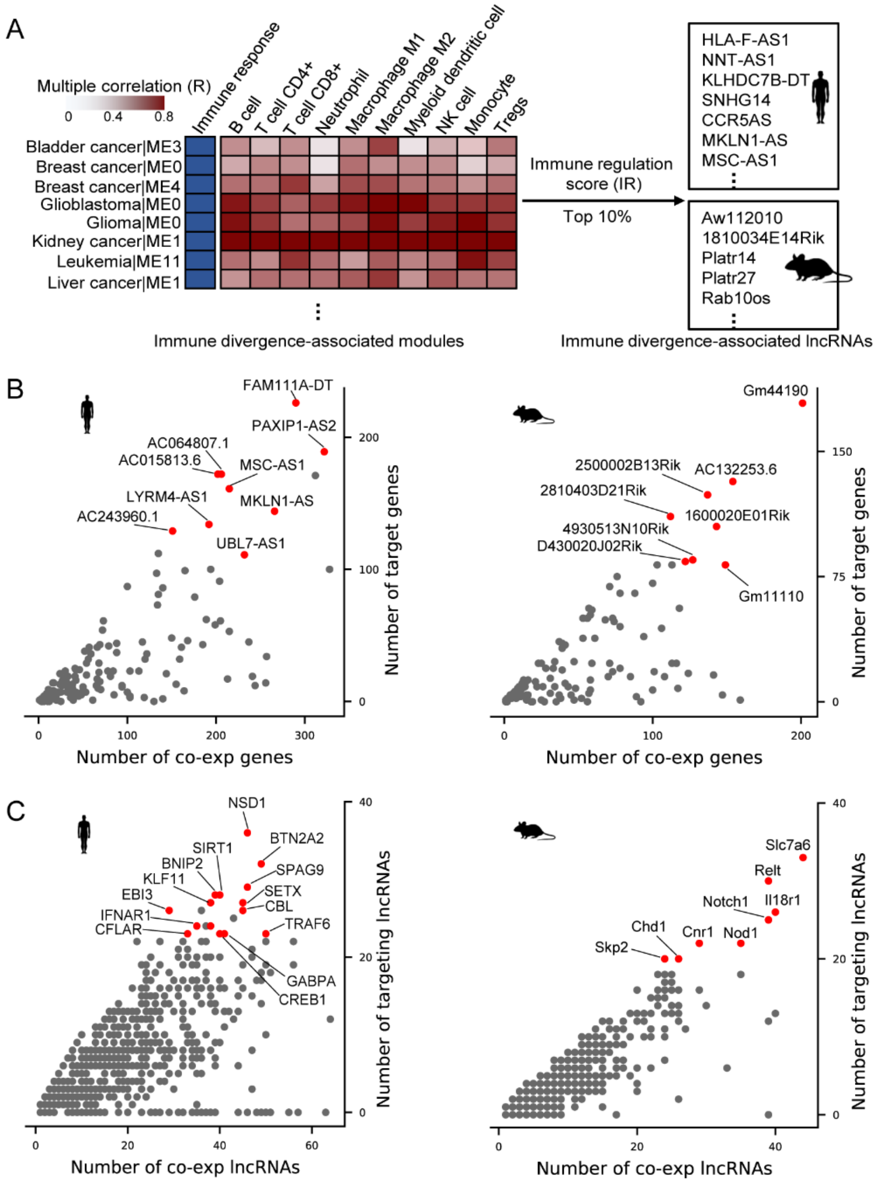


Supplementary Figure 6-2. Disproportionate, many-to-many topological distribution of LS lncRNA-mediated immune regulation. (A) Identification of ID-associated modules and specific ID-associated lncRNAs. Modules were ranked by their multiple correlation coefficients with specific immune cell infiltration proportions, enabling the identification of the top 10% of lncRNAs (ranked by Immune Regulation [IR] score) as prominent regulators. (B) Scatter plots illustrating the regulatory reach of individual LS lncRNAs in humans (left) and mice (right). Red dots highlight top "hub" lncRNAs possessing the highest volume of target TDGs and co-expressed genes within these immune modules. (C) Scatter plots illustrating the target density of individual TDGs. Red dots highlight essential "hub" target genes that are heavily targeted by multiple LS lncRNAs. Identified human target hubs with previously validated experimental roles in tumor immunity include *IFNAR1* (Zitvogel et al., 2015), *CFLAR* (Humphreys et al., 2018), *BTN2A2* (Sarter et al., 2016). Distinctly targeted murine hubs include *Il18r1*, *Notch1* (Roper et al., 2021), and *Relt* (Choi et al., 2018).

### References


1. Affo, S., Yu, L.X., and Schwabe, R.F. (2017). The Role of Cancer-Associated Fibroblasts and Fibrosis in Liver Cancer. Annu Rev Pathol *12*, 153-186.
2. Alexandrov, L.B., Nik-Zainal, S., Wedge, D.C., Campbell, P.J., and Stratton, M.R. (2013). Deciphering signatures of mutational processes operative in human cancer. Cell Rep *3*, 246-259.
3. Basso, K., and Dalla-Favera, R. (2015). Germinal centres and B cell lymphomagenesis. Nat Rev Immunol *15*, 172-184.
4. Brennan, C.W., Verhaak, R.G., McKenna, A., Campos, B., Noushmehr, H., Salama, S.R., Zheng, S., Chakravarty, D., Sanborn, J.Z., Berman, S.H.*, et al.* (2013). The somatic genomic landscape of glioblastoma. Cell *155*, 462-477.
5. Chan, J.E., Pan, C.H., Rub, J., Guzman, G., Krause, K., Brown, E., Zhang, Z., Styers, H., Hartmann, G., Li, Z.*, et al.* (2026). Critical role for a high-plasticity cell state in lung cancer. Nature *651*, 231-241.
6. Chen, J., Li, Y., Yu, T.S., McKay, R.M., Burns, D.K., Kernie, S.G., and Parada, L.F. (2012). A restricted cell population propagates glioblastoma growth after chemotherapy. Nature *488*, 522-526.
7. Choi, B.K., Kim, S.H., Kim, Y.H., Lee, D.G., Oh, H.S., Han, C., Kim, Y.I., Jeon, Y., Lee, H., and Kwon, B.S. (2018). RELT negatively regulates the early phase of the T-cell response in mice. Eur J Immunol *48*, 1739-1749.
8. Dankort, D., Curley, D.P., Cartlidge, R.A., Nelson, B., Karnezis, A.N., Damsky, W.E., Jr., You, M.J., DePinho, R.A., McMahon, M., and Bosenberg, M. (2009). Braf(V600E) cooperates with Pten loss to induce metastatic melanoma. Nat Genet *41*, 544-552.
9. Gao, Q., Zhu, H., Dong, L., Shi, W., Chen, R., Song, Z., Huang, C., Li, J., Dong, X., Zhou, Y.*, et al.* (2019). Integrated Proteogenomic Characterization of HBV-Related Hepatocellular Carcinoma. Cell *179*, 1240.
10. Grizzi, F., Hegazi, M., Zanoni, M., Vota, P., Toia, G., Clementi, M.C., Mazzieri, C., Chiriva-Internati, M., and Taverna, G. (2023). Prostate Cancer Microvascular Routes: Exploration and Measurement Strategies. Life (Basel) *13*.
11. Hanahan, D. (2022). Hallmarks of Cancer: New Dimensions. Cancer Discov *12*, 31-46.
12. Herschkowitz, J.I., Simin, K., Weigman, V.J., Mikaelian, I., Usary, J., Hu, Z., Rasmussen, K.E., Jones, L.P., Assefnia, S., Chandrasekharan, S.*, et al.* (2007). Identification of conserved gene expression features between murine mammary carcinoma models and human breast tumors. Genome Biol *8*, R76.
13. Hingorani, S.R., Wang, L., Multani, A.S., Combs, C., Deramaudt, T.B., Hruban, R.H., Rustgi, A.K., Chang, S., and Tuveson, D.A. (2005). Trp53R172H and KrasG12D cooperate to promote chromosomal instability and widely metastatic pancreatic ductal adenocarcinoma in mice. Cancer Cell *7*, 469-483.
14. Humphreys, L., Espona-Fiedler, M., and Longley, D.B. (2018). FLIP as a therapeutic target in cancer. FEBS J *285*, 4104-4123.
15. Lin, C.Y., Kleinbrink, E.L., Dachet, F., Cai, J., Ju, D., Goldstone, A., Wood, E.J., Liu, K., Jia, H., Goustin, A.S.*, et al.* (2016). Primate-specific oestrogen-responsive long non-coding RNAs regulate proliferation and viability of human breast cancer cells. Open Biol *6*.
16. Miyai, M., Tomita, H., Soeda, A., Yano, H., Iwama, T., and Hara, A. (2017). Current trends in mouse models of glioblastoma. J Neurooncol *135*, 423-432.
17. Pfefferle, A.D., Herschkowitz, J.I., Usary, J., Harrell, J.C., Spike, B.T., Adams, J.R., Torres-Arzayus, M.I., Brown, M., Egan, S.E., Wahl, G.M.*, et al.* (2013). Transcriptomic classification of genetically engineered mouse models of breast cancer identifies human subtype counterparts. Genome Biol *14*, R125.
18. Phillips, J.W., Pan, Y., Tsai, B.L., Xie, Z., Demirdjian, L., Xiao, W., Yang, H.T., Zhang, Y., Lin, C.H., Cheng, D.*, et al.* (2020). Pathway-guided analysis identifies Myc-dependent alternative pre-mRNA splicing in aggressive prostate cancers. Proc Natl Acad Sci U S A *117*, 5269-5279.
19. Rambow, F., Marine, J.C., and Goding, C.R. (2019). Melanoma plasticity and phenotypic diversity: therapeutic barriers and opportunities. Genes Dev *33*, 1295-1318.
20. Robertson, A.G., Kim, J., Al-Ahmadie, H., Bellmunt, J., Guo, G., Cherniack, A.D., Hinoue, T., Laird, P.W., Hoadley, K.A., Akbani, R.*, et al.* (2017). Comprehensive Molecular Characterization of Muscle-Invasive Bladder Cancer. Cell *171*, 540-556 e525.
21. Roper, N., Velez, M.J., Chiappori, A., Kim, Y.S., Wei, J.S., Sindiri, S., Takahashi, N., Mulford, D., Kumar, S., Ylaya, K.*, et al.* (2021). Notch signaling and efficacy of PD-1/PD-L1 blockade in relapsed small cell lung cancer. Nat Commun *12*, 3880.
22. Sarter, K., Leimgruber, E., Gobet, F., Agrawal, V., Dunand-Sauthier, I., Barras, E., Mastelic-Gavillet, B., Kamath, A., Fontannaz, P., Guery, L.*, et al.* (2016). Btn2a2, a T cell immunomodulatory molecule coregulated with MHC class II genes. J Exp Med *213*, 177-187.
23. Venteicher, A.S., Tirosh, I., Hebert, C., Yizhak, K., Neftel, C., Filbin, M.G., Hovestadt, V., Escalante, L.E., Shaw, M.L., Rodman, C.*, et al.* (2017). Decoupling genetics, lineages, and microenvironment in IDH-mutant gliomas by single-cell RNA-seq. Science *355*.
24. Zitvogel, L., Galluzzi, L., Kepp, O., Smyth, M.J., and Kroemer, G. (2015). Type I interferons in anticancer immunity. Nat Rev Immunol *15*, 405-414.
